# Molecular recording of cellular protein kinase activity with chemical labeling

**DOI:** 10.1101/2024.09.11.611894

**Authors:** De-en Sun, Siu Wang Ng, Yu Zheng, Shu Xie, Niklas Schwan, Paula Breuer, Dirk C. Hoffmann, Julius Michel, Daniel D. Azorin, Kim E. Boonekamp, Frank Winkler, Wolfgang Wick, Michael Boutros, Yulong Li, Kai Johnsson

## Abstract

Protein kinases control most cellular processes and aberrant kinase activity is involved in numerous diseases. To investigate the link between specific kinase activities and cellular phenotypes in heterogeneous cell populations and *in vivo*, we introduce molecular recorders of kinase activities for later analysis. Based on split-HaloTag and a phosphorylation-dependent molecular switch, our recorders become rapidly labeled in the presence of a specific kinase activity and a fluorescent HaloTag substrate. The kinase activity in a given cell controls the degree of fluorescent labeling whereas the recording window is set by the presence of the fluorescent substrate. We have designed specific recorders for four protein kinases, including protein kinase A. We apply our protein kinase A recorder for the sorting of heterogeneous cell populations and subsequent transcriptome analysis, in genome-wide CRISPR screens to discover regulators of PKA activity and for the tracking of neuromodulation in freely moving mice.

## Introduction

Protein kinase signaling cascades are essential in nearly all cellular processes. Aberrant kinase signaling, which disrupts the balance of protein phosphorylation, is tightly linked to tumorigenesis and neuronal dysfunction^1^. A prototypical kinase is cAMP-dependent protein kinase (i.e., protein kinase A, or PKA), a major cAMP effector, which integrates different signaling pathways, including neuromodulation, metabolism and proliferation to control a vast number of physiological processes^2^. Monitoring the activity of specific kinases thus can provide important insights into cellular physiology, an example being the measurements of PKA activities as a readout for neuromodulation in the brain^3–13^. Currently, phosphorylation state-specific antibodies are widely used to analyze protein phosphorylation levels at a single timepoint in cell lysates and fixed samples. However, the inaccessibility of the antigen to the antibody can compromise the sensitivity and quantifiability of the approach^14^. Genetically encoded kinase activity reporters with fluorescence or luminescence readouts allow real-time monitoring of kinase activity *ex vivo* and *in vivo*^15^. While these sensors offer exquisite spatiotemporal resolution, the restricted field of view of optical approaches and the limited penetration of light into tissue poses some restrictions with respect to number of cells and tissue regions investigated. Furthermore, long-term *in vivo* imaging remains laborious and requires custom-built imaging equipment. A recent optogenetic approach, KINACT, relies on reporter gene expression that is induced both by kinase activity and illumination with blue light^16^. Converting transient kinase signals into a “permanent” mark separates recording from analysis and thus facilitates the parallel investigation of large number of cells. However, the approach requires reporter gene expression for one day after illumination, and still suffers from the same limitations as other optical approaches. Directly recording kinase activity with high temporal resolution in a scalable manner or in deep tissues without the need for illumination thus remains challenging.

To address these challenges and complement the aforementioned approaches, we developed split-HaloTag recorders for <u>ki</u>nase activity-dependent <u>pro</u>tein <u>la</u>beling (Kinprola). Kinprola enables the accumulation of an irreversible label in the presence of both a specific kinase activity and a fluorescent HaloTag substrate. Kinprola allows scalable recording of kinase activities for later analysis in different cellular compartments, and its modular design enables the generation of Kinprola variants for different kinases. We demonstrate the versatility of Kinprola for applications in transcriptome analysis, in functional genomic screening and for the tracking of neuromodulation in freely moving mice.

## Results

### Development of Kinprola

We first focused on the generation of a recorder for PKA, Kinprola_PKA_, in which reversible phosphorylation of Kinprola_PKA_ by PKA would activate its labeling activity. To develop Kinprola_PKA_, we utilized the split-HaloTag system recently developed by our group^17^. The system is comprised of a truncated, circularly permutated HaloTag (cpHaloΔ) that retains the overall fold of HaloTag but exhibits almost no activity, and a decapeptide (Hpep) that can bind to cpHaloΔ and restore its activity towards chloroalkane (CA) substrates. We envisioned generating Kinprola_PKA_ by connecting cpHaloΔ and Hpep through a linker comprising the forkhead-associated domain 1 (FHA1) and a PKA-specific substrate peptide (PKAsub). Specific binding of FHA1 to phosphorylated PKAsub should then result in a conformational change of Kinprola_PKA_ that enables the activation of cpHaloΔ by binding to Hpep (Fig.1a and Extended Data Fig. 1a). We used structural information on the FHA1–phosphothreonine peptide complex^18^ to design circularly permuted FHA1 variants and tested, together with wild-type FHA1, their performance in Kinprola_PKA_. We identified circularly permuted FHA1 variants with new C and N termini at positions 53 and 54, respectively, that showed a strong dependence of labeling rates on phosphorylation of Kinprola_PKA_. Additionally, we incorporated the FHA1 mutation N49Y which is known to enhance protein thermostability^19^ (Extended Data Fig. 1b and Supplementary Table 1). Finally, we systematically optimized the length and composition of the linker sequences and screened different Hpep variants. The final version of Kinprola_PKA_ showed no significant labeling with a fluorescent tetramethyl-rhodamine HaloTag substrate (TMR-CA) in its non-phosphorylated form, but exhibited a more than thousandfold increase in labeling speed after being phosphorylated by PKA catalytic subunit (PKAcat) (i.e., second-order rate constant after phosphorylation (k_TMR-CA_) = 1.37×10^5^ M^−1^s^−1^; Fig. 1b, Extended Data Fig. 2 and Supplementary Table 2). When mutating the phospho-acceptor site threonine to alanine (T/A) in the PKAsub, resulting in Kinprola_PKA_T/A_, no increase in labeling rate was observed after incubation with PKAcat and ATP (Fig. 1b and Extended Data Fig. 2). Kinprola_PKA_ was also rapidly labeled in a PKA-dependent manner by several other spectrally distinct fluorescent substrates (Extended Data Fig. 2, Supplementary Fig. 1 and Supplementary Table 2).

**Fig.1.**
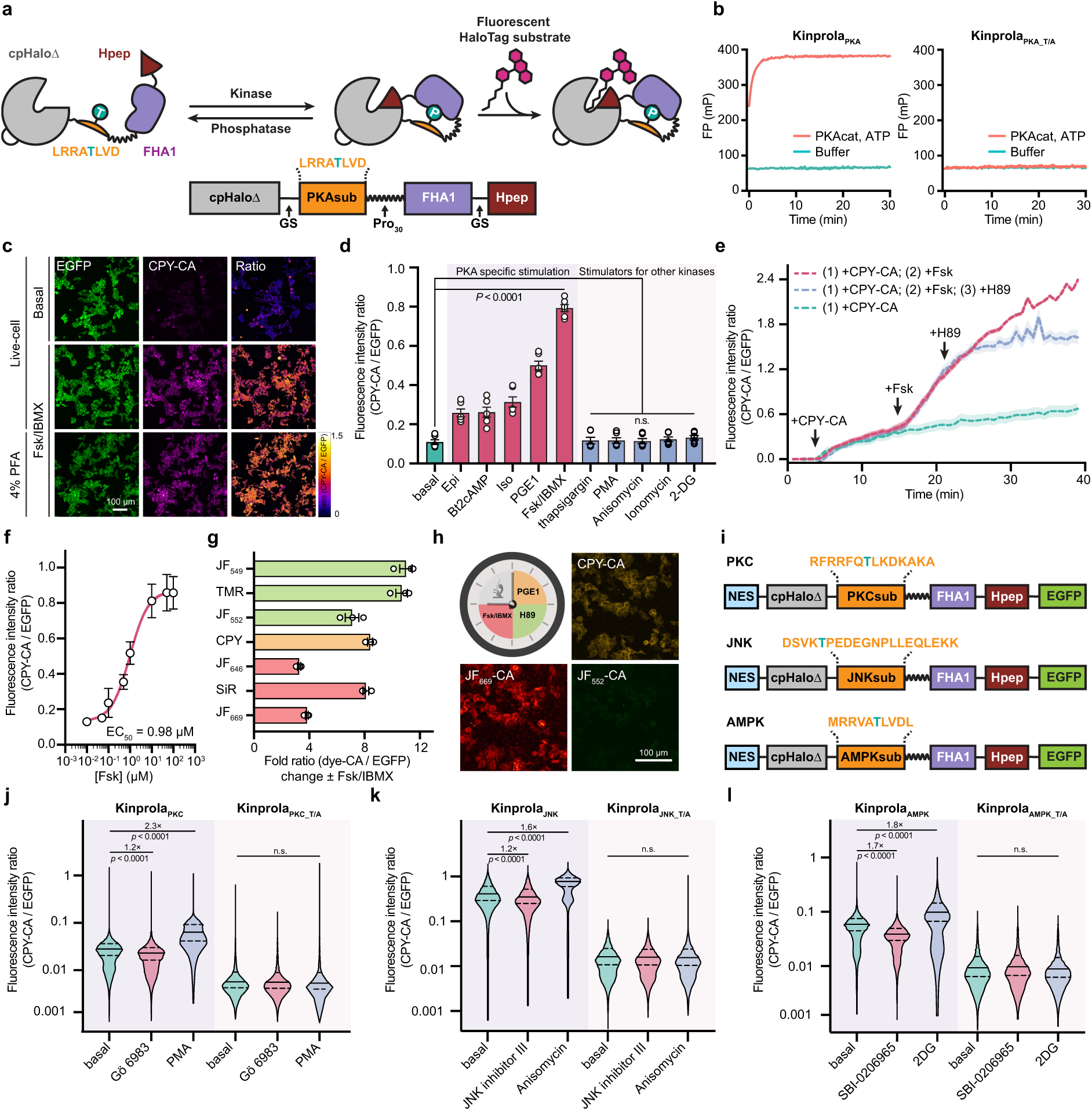
| Design and characterization of the Kinprola recorder. (**a**) Schematic diagram depicting the design of the Kinprola_PKA_ recorder, consisting of the recording moiety cpHaloΔ–Hpep and sensing moiety FHA1–PKAsub. Phosphorylation on the PKAsub activates the Kinprola_PKA_ recorder, inducing the accumulation of fluorescence labeling. (**b**) Labeling kinetics of Kinprola_PKA_ and Kinprola_PKA_T/A_ (200 nM) with TMR-CA (50 nM) in the presence or absence of PKAcat (25 ng μL^−1^) and ATP (500 μM) measured by fluorescence polarization (FP), data from three technical replicates. (**c**) Fluorescence images of HEK293 cells stably expressing Kinprola_PKA_ treated with 50 μM Fsk/100 μM IBMX for 30 min in the presence of 25 nM CPY-CA. Live-cell and 4% PFA-fixed cell images are shown. Representative images from three wells of cell culture. (**d**) Flow cytometry analysis of HEK293 cells stably expressing Kinprola_PKA_ treated with different stimulators for 30 min in the presence of 25 nM CPY-CA. Error bars indicate mean ± SEM (from three independent experiments with duplicates). (**e**) Time-lapse fluorescence traces recorded on HEK293 cells stably expressing Kinprola_PKA_ successively treated with 50 μM Fsk and 20 μM H89 in the presence of 25 nM CPY-CA. Cells only treated with Fsk or without treatment were acquired for comparison. Signal background from free CPY-CA was calibrated using co-cultured cells without Kinprola_PKA_ expression. Ratios indicate CPY-CA/EGFP. Representative traces from three independent experiments with replicates. (**f**) Flow cytometry analysis of HEK293 cells stably expressing Kinprola_PKA_ treated with varying concentrations of Fsk in the presence of 25 nM CPY-CA for 30 min. Error bars indicate mean ± SEM (from three independent experiments with replicates). The curve was fitted with a sigmoidal function to determine the EC_50_ value. (**g**) Flow cytometry analysis of HeLa cells stably expressing Kinprola_PKA_ labeled with different HaloTag fluorescent substrates (25 nM, 30 min) in the presence or absence of 50 μM Fsk/100 μM IBMX stimulation. Error bars indicate mean ± SEM (from three independent experiments with duplicates). (**h**) Recording of three successive periods of PKA activity in HEK293 cells stably expressing Kinprola_PKA_. Cells were allowed to rest for 2 h between each treatment. Imaging was performed after the third incubation period. Representative images from three independent experiments with replicates. (**i**) Domain structures of Kinprola recorders for PKC, JNK and AMPK. (**j**-**l**) Flow cytometry analysis of labeling in HeLa cells transiently expressing different Kinprolas (25 nM CPY-CA, 30 min) in the presence of activators or inhibitors for PKC, JNK and AMPK, respectively. Error bars indicate median with interquartile range (from three independent experiments with triplicates). Statistical significance was calculated with one-way ANOVA with Dunnett’s Post hoc test (**d**,**j**-**l**) and *p* values are given for comparison, n.s., not significant (*p* > 0.05). Scale bars: 100 μm (**c**,**h**). Kinprolas for other kinases can be created by substituting the PKAsub with substrate peptides specific for other kinases such as protein kinase C (PKC)^23^, c-Jun N-terminal kinases (JNKs)^24^ and AMP-activated protein kinase (AMPK)^25^. Without additional engineering, these Kinprola variants recorded the activities of their cognate kinases during drug stimulation or inhibition in the cytosol of mammalian cells lines, with minimal responses observed in T/A mutant negative controls (Fig. 1i-l and Extended Data Fig. 6a,b). Furthermore, Kinprola_PKA_ can be (simultaneously) used in different cellular compartments. For this, Kinprola_PKA_ variants with different localization tags were simultaneously expressed in cytoplasm and nucleus of single cells. Different fluorescent proteins were introduced to normalize the labeling intensity separately (Extended Data Fig. 6c). Fsk/IBMX stimulation in the presence of CPY-CA, resulted in increased fluorescent labeling both in the nucleus and the cytosol compared to fluorescent labeling observed in the absence of Fsk/IBMX (Extended Data Fig. 6d,e).

To record cytosolic PKA activities in cultured mammalian cells, we fused a nuclear export signal (NES) sequence and EGFP to Kinprola_PKA_. EGFP was introduced to normalize the fluorescence signal of labeled Kinprola_PKA_ for differences in expression levels of the recorder. We also constructed constitutively active Kinprola_on_ by replacing cpHaloΔ with full length cpHalo, and constitutively inactive Kinprola_off_ by deleting Hpep. Similar to Kinprola_PKA_T/A_, Kinprola_off_ only accumulated background labeling (Extended Data Fig. 3a). HEK293 cells expressing Kinprola_PKA_ were incubated with a fluorescent carbopyronine HaloTag substrate (CPY-CA) in the presence or absence of the adenylyl cyclase activator forskolin (Fsk) and the pan-phosphodiesterase inhibitor 3-isobutyl-1-methylxanthine (IBMX), a stimulation that induces high PKA activity by raising cellular cAMP levels^12,20^. Incubation with Fsk/IBMX led to strong labeling whereas very little labeling was observed in its absence. Furthermore, the fluorescent labeling resisted subsequent chemical fixation (Fig. 1c and Supplementary Fig. 2). Flow cytometry analysis revealed an 8-fold change in normalized fluorescence intensity (CPY-CA/EGFP) between Fsk/IBMX-treated and untreated cells (Fig. 1d and Supplementary Fig. 3). Labeling times as short as 5 minutes and CPY-CA concentrations as low as 5 nM were sufficient to detect a difference between Fsk/IBMX-treated and untreated cells (Extended Data Fig. 3). In contrast, Kinprola_on_ showed negligible differences with or without Fsk/IBMX treatment, and neither Kinprola_off_ nor Kinprola_PKA_T/A_ showed significant labeling (Extended Data Fig. 3). Kinprola_PKA_ labeling was dose-dependent on Fsk, and directly raising intracellular cAMP concentrations with the cell-permeable cAMP analog (Bt_2_cAMP) also increased Kinprola_PKA_ labeling (Fig. 1d,f and Extended Data Fig. 4). The cAMP/PKA pathway acts as a central downstream mediator of multiple G protein-coupled receptor (GPCR) signaling pathways and Kinprola_PKA_ also strongly responded to several key Gα_s_-coupled receptor agonists such as epinephrine (Epi), isoproterenol (Iso) and prostaglandin E1 (PGE1) in a dose-dependent manner (Fig. 1d and Extended Data Fig. 4). Furthermore, incubation with the PKA inhibitor H89 further suppressed the already low basal labeling of Kinprola_PKA_ (Extended Data Fig. 4). Various stimulators of kinases other than PKA did not increase Kinprola_PKA_ labeling, confirming that Kinprola_PKA_ specifically reports on PKA activation (Fig. 1d). While Kinprola_PKA_ was designed to record periods of PKA activity for later analysis, it can also be used for real-time recording. Addition of Fsk to HEK293 cells expressing Kinprola_PKA_ resulted in an increase in Kinprola_PKA_ labeling as measured by time-lapse fluorescence microscopy. In these experiments, the rate of Fsk-induced labeling can be attenuated to the basal labeling level by the addition of the PKA inhibitor H89. This demonstrates that Kinprola_PKA_ responds to sudden changes in cellular PKA activity (Fig. 1e). Furthermore, a variety of different fluorescent HaloTag substrates can be used for the Fsk/IBMX-dependent fluorescent marking of Kinprola_PKA_-expressing cells (Fig. 1g and Supplementary Fig. 1) and we leveraged this to distinguish multiple recording periods in a single cell. Specifically, cells expressing Kinprola_PKA_ were first incubated with CPY-CA and PGE1 (moderate stimulation of PKA), then with the fluorescent Janelia Fluor 552 HaloTag substrate (JF_552_-CA)^21^ in the presence of H89 (inhibition of PKA), and finally with JF_669_-CA^22^ and Fsk/IBMX (strong stimulation of PKA). Post hoc imaging showed the expected pattern of moderate labeling with CPY, weak labeling with JF_552_, and strong labeling with JF_669_ (Fig. 1h). By switching the combination of fluorescent substrates and drugs, the differences in labeling efficiencies for the three periods were observed to be independent of which fluorescent substrate was used for which recording period (Extended Data Fig. 5).

### Stably marking and selecting cell subpopulations by Kinprola_PKA_ for transcriptome analysis

Recording transient kinase activity for later analysis is particularly valuable for correlating cellular phenotypes with kinase signaling in large and heterogeneous cell populations. An example of a heterogeneous cell combination is glioblastoma, the most frequent and aggressive adult-type diffuse glioma^26,27^. Glioblastoma cells (GBCs) exhibit diverse phenotypic and behavioral characteristics. For instance, the formation of tumor microtubes diversifies GBC invasiveness, which also positively correlates with their proliferation^28–31^. Recently, using Caprola_6_, a split-HaloTag recorder for cytosolic calcium transients, we observed heterogeneous calcium signaling signatures in GBC subpopulations^17,31^. To investigate whether GBC subpopulations selected using Kinprola_PKA_ with varying PKA activities, we expressed Kinprola_PKA_ in patient-derived GBCs and cultured them in a 2D monoculture under serum-free stem-like conditions, in which GBCs retain their capacity for tumor microtubes formation^31,32^. Following labeling with CPY-CA, GBCs were sorted into high, medium and low normalized labeling intensity groups using fluorescence-activated cell sorting (FACS) and subjected to bulk RNA sequencing (RNA-Seq) (Fig. 2a and Supplementary Fig. 4). As a control, GBCs expressing Kinprola_on_ underwent the same procedure to eliminate potential labeling differences due to CPY-CA permeability heterogeneity (Supplementary Fig. 4).

**Fig.2.**
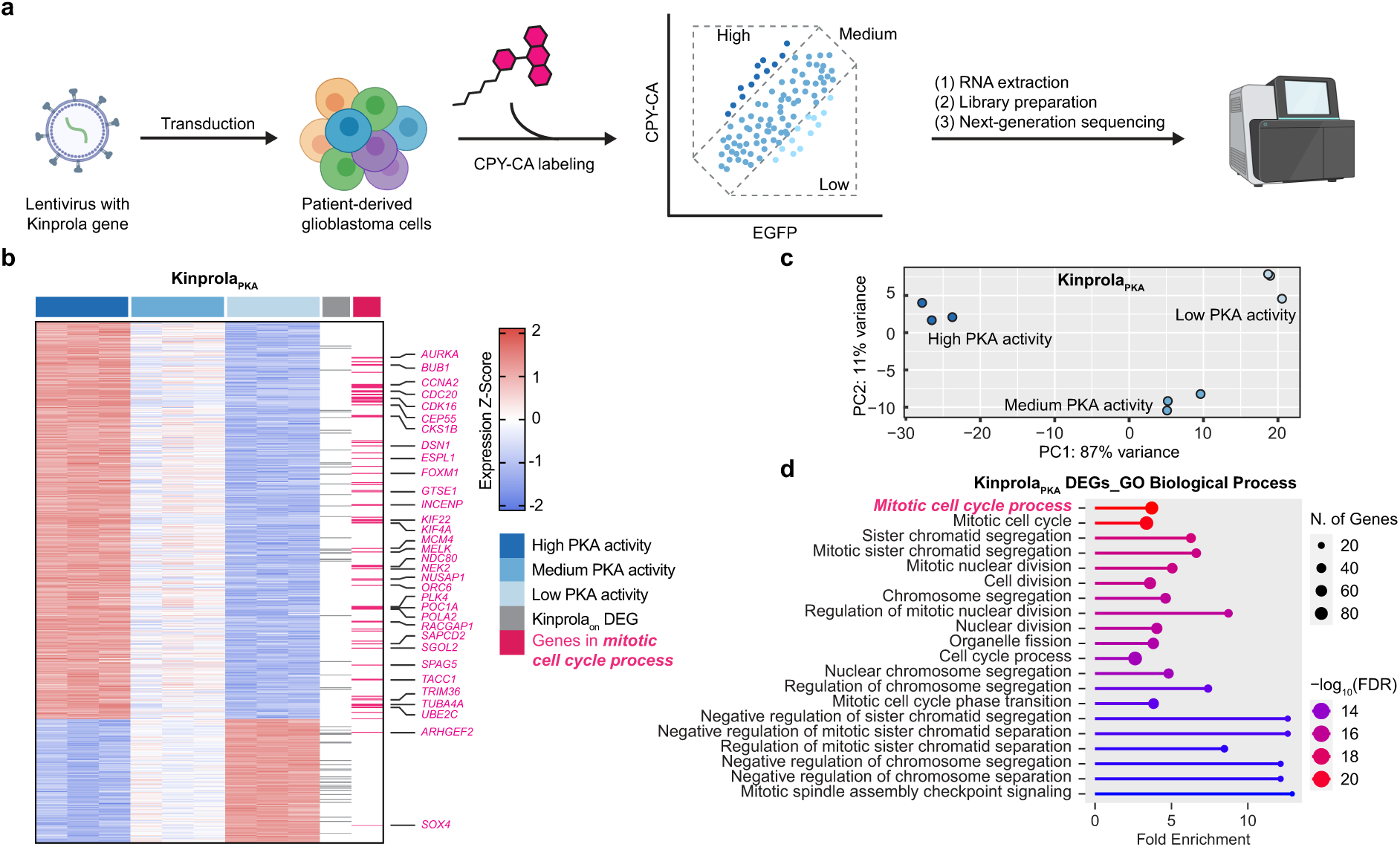
| Kinprola enables the selection of cell subpopulations based on PKA activity for subsequent transcriptome analysis. (**a**) Schematic diagram depicting the strategy of Kinprola_PKA_ recording (100 nM CPY-CA, 30 min) in glioblastoma cells (GBCs) for subsequent bulk RNA-Seq analysis. (**b**) Transcriptional profiles of the three sorted groups of Kinprola_PKA_-expressing GBCs. DEGs identified by RNA-Seq analysis are color coded according to the Z-score. Genes in the GO term “mitotic cell cycle process” are highlighted in magenta and representative gene symbols are listed. Overlapped DEGs identified in both Kinprola_PKA_ and Kinprola_on_ groups are indicated in gray. RNA-Seq data was obtained from triplicates. (**c**) Dot plots of the first principal coordinate analysis on the three sorted groups of Kinprola_PKA_-expressing GBCs. (**d**) Biological process GO terms of Kinprola_PKA_-identified DEGs. The top twenty GO terms ordered by false discovery rate (FDR) are shown.

The transcriptomic profiles of three groups were clearly separated in Kinprola_PKA_, and 737 differentially expressed genes (DEGs) between all groups in pairwise group analysis were identified (Fig. 2b,c). Among the 737 Kinprola_PKA_-identified DEGs, only 55 DEGs were overlapped with 327 DEGs identified by Kinprola_on_ (Extended Data Fig. 7a,b), suggesting their unique signatures. Subsequent Gene ontology (GO) analysis, conducted without the coinciding DEGs, revealed that GO terms associated with proliferation, such as mitotic cell cycle process and extracellular matrix were over-represented in the 682 Kinprola_PKA_ specific DEGs (Fig. 2d and Extended Data Fig. 7c). In contrast, the molecular function of DEGs identified by Kinprola_on_ was associated with transporters and other features (Extended Data Fig. 7d,e). This observation is consistent with previous studies on PKA activity oscillations during the cell mitotic process^33–35^, suggesting the impact of PKA activity for GBC proliferation. Taking together, these experiments demonstrates the capability of Kinprola for stably marking, selecting and analyzing cell subpopulations within heterogeneous networks in a high-throughput and scalable manner.

### Combining Kinprola_PKA_ with CRISPR knockout screening to identify regulators of PKA signaling

To demonstrate the capability of Kinprola_PKA_ for identifying potential regulators of PKA, we combined a pooled CRISPR knockout screening approach with Kinprola_PKA_, which can select cells based on their relative PKA activities after genetic perturbation. Firstly, transgenes for Kinprola_PKA_ and Cas9 were introduced into RKO colon cancer cells by lentiviral integration. Cells stably expressing the two transgenes exhibited robust responses to both drug stimulation and inhibition compared to cells expressing Kinprola_on_ and the T/A mutant (Extended Data Fig. 8a). To conduct pooled genetic perturbation, the stable cells were transduced with a lentiviral genome-wide sgRNA library^36^ at a multiplicity of infection (MOI) of 0.2 to 0.3, followed by puromycin selection of transduced cells, expansion, and labeling with CPY-CA at day 6 after transduction. The labeled cells were then sorted into three populations based on their normalized labeling intensity (high 25%, lowest 25%, and medium) (Fig. 3a and Extended Data Fig. 8b). Next-generation sequencing was performed on each sorted sample to determine the abundance of each sgRNA present in the three cell subpopulations. By comparing the sgRNA representation in each subpopulation, sgRNAs that significantly altered Kinprola_PKA_ labeling were inferred. The screen was independently performed twice to identify differentially regulated gene targets (Extended Data Fig. 8c-e). Of the 18,659 protein coding genes targeted by the sgRNA library, a total of 340 hits from the “high” *vs.* “low” comparison were selected with an FDR threshold of 0.05 (Fig. 3b). 45 genes were identified that decreased normalized labeling intensity, which represent knockouts that decreased PKA activity, whereas 295 genes were identified that increased PKA activity after knockout. Notably, canonical regulators in the GPCR-cAMP-PKA signaling pathway, including the catalytic subunit α of PKA (*PRKACA*), the heterotrimeric G protein Gα_s_ subunit (*GNAS*) and the adenylyl cyclase 7 (*ADCY7*), were among the identified hits for which knockout decreased PKA activity. These three regulators were also found to be enriched in the “low” *vs.* “medium” comparison, but not in the “high” *vs.* “medium” comparison (Fig. 3b and Extended Data Fig. 8f-h). Gene set enrichment analysis (GSEA) was subsequently performed to identify relevant biological processes that were enriched among the hits. GO terms related to stress response and metabolic process were over-represented in hits for which knockout increased PKA activity, whereas terms associated with GPCR signaling pathways, second messenger-mediated signaling, and nucleosome organization were over-represented in collections of genes for which knockout decrease PKA activity (Fig. 3c, Extended Data Fig. 8i,j and Supplementary Fig. 5).

**Fig.3.**
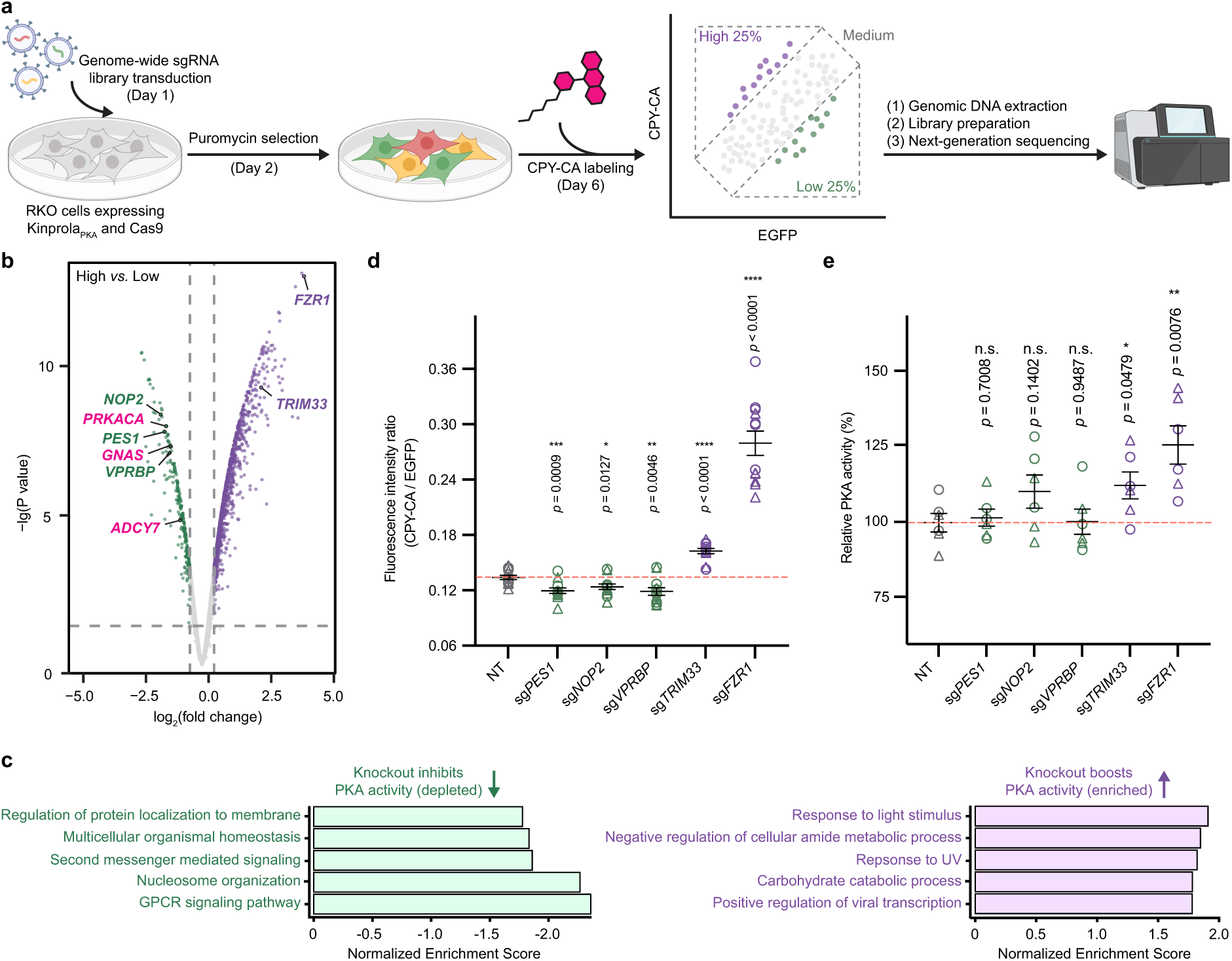
| Kinprola for dissecting regulators of PKA signaling through CRISPR knockout screening. (**a**) Schematic diagram depicting the strategy of cell subpopulation selection using KinprolaPKA in RKO cells from pooled CRISPR knockout screen analysis. (**b**) Volcano plot of gene enrichment from comparison of “high” *vs.* “low” fractions. Dashed lines depict FDR cutoff (0.05) for hit gene selection. Putative hits and canonical regulators including *PRKACA*, *GNAS* and *ADCY7* are highlighted. CRISPR knockout screen data were obtained from two independent experiments. (**c**) Gene set enrichment analysis (GSEA) of the top 5 categories among “high” *vs.* “low” comparison. (**d**) Dot plot showing normalized Kinprola_PKA_ labeling intensity (CPY-CA/EGFP) in cells with individual knockouts of putative regulators. NT group denotes cells with non-targeting (NT) sgRNAs. The dots with circle and triangle shape represent two different sgRNAs targeting the same gene. Error bars indicate mean ± SEM (from six independent experiments with duplicates). (**e**) Validation of putative regulators in cell lysates using an ELISA-based PKA colorimetric activity assay. PKA activities of each group were normalized by NT group. The dots with circle and triangle shape represent two different sgRNAs targeting the same gene. Error bars indicate mean ± SEM (from three independent experiments with duplicates). The same cell line was used for both the screen (**b**,**c**) and the single sgRNA knockouts (**d**,**e**). Statistical significance was calculated with unpaired two-tailed Welch’s *t* test and *p* values are given for comparison (**d**,**e**).

We then examined three genes for which knock-out decreased PKA activity, i.e., *PES1*, *NOP2*, and *VPRBP*, and two genes for which knock-out increased PKA activity, i.e., *TRIM33* and *FZR1*. For each of the five genes, we used two individual sgRNAs for knockouts and examined their effect on PKA activity as measured by Kinprola_PKA_ in the cell line used in the CRISPR screen. Consistent with the results from the CRISPR screen, knockout of *PES1*, *NOP2*, and *VPRBP* decreased PKA activity, and knockout of *TRIM33* and *FZR1* increased PKA activity as compared to non-targeting (NT) sgRNAs (Fig. 3d). Furthermore, we attempted to measure PKA activity in cell lysates of these five knockout lines. Using an ELISA-based PKA colorimetric activity assay, we were able to detect a significant increase in PKA activity in lysates of cells in which *FZR1* was knocked out. For the four other knock-outs, only subtle or nonsignificant changes were observed in cell lysates (Fig. 3e). This can be attributed to the low sensitivity of the underlying assay and the non-physiological conditions during the measurement. *FZR1* encodes cdh1, an important component of the anaphase promoting complex/cyclosome (APC/C), which controls cell cycle fate decisions^37^. Knockout of *FZR1* leads to unscheduled cell cycle progression, subsequently triggering replicative stress and DNA damage responses^38^. In addition, previous studies showed that PKA activity oscillates throughout the cell cycle, and negatively regulates APC/C by phosphorylating several of its components to ensure proper activation^33,35^. Our screen results thus indicate a potential regulatory connection between *FZR1* function and PKA activity. Overall, our CRISPR screen highlights the influence of diverse biological processes on the regulation of PKA signaling and outlines how Kinprola can be used in genetic screens to unravel genes regulating kinase activity.

### Recording PKA activation in primary neurons, acute brain slices and freely moving mice

PKA integrates multiple GPCR signaling pathways and plays a key role in neuronal excitability and plasticity. Consequently, monitoring cellular PKA activity provides a valuable readout for neuromodulatory events^3–13^. To investigate if Kinprola_PKA_ can be used to track PKA activity changes in the nervous system, we firstly expressed Kinprola in cultured primary rat hippocampal neurons. Compared with Kinprola_on_ and the T/A mutant, neurons expressing Kinprola_PKA_ exhibited a robust increase in labeling relative to basally active neurons when stimulated with Fsk/Rolipram (Rol) or Iso. Conversely, PKA inhibition by H89 or synaptic transmission silencing between neurons with glutamate receptor antagonists NBQX/APV resulted in decreased labeling compared to basally active neurons (Fig. 4a,b and Extended Data Fig. 9a,b). Additionally, neurons expressing Kinprola_PKA_ responded to stimulation by the neuromodulator norepinephrine, with the response effectively blocked by co-treatment with the β-adrenergic receptor antagonist propranolol (Extended Data Fig. 9c,d). In NBQX/APV-silenced neurons, Kinprola_PKA_ effectively recorded the increase in cytosolic PKA activity elicited by electrically evoked action potentials (Fig. 4c,d and Extended Data Fig. 9e,f). Furthermore, the fluorescent Kinprola_PKA_ labeling signal in live neurons remained detectable for at least three days after recording (Supplementary Fig. 6).

**Fig.4.**
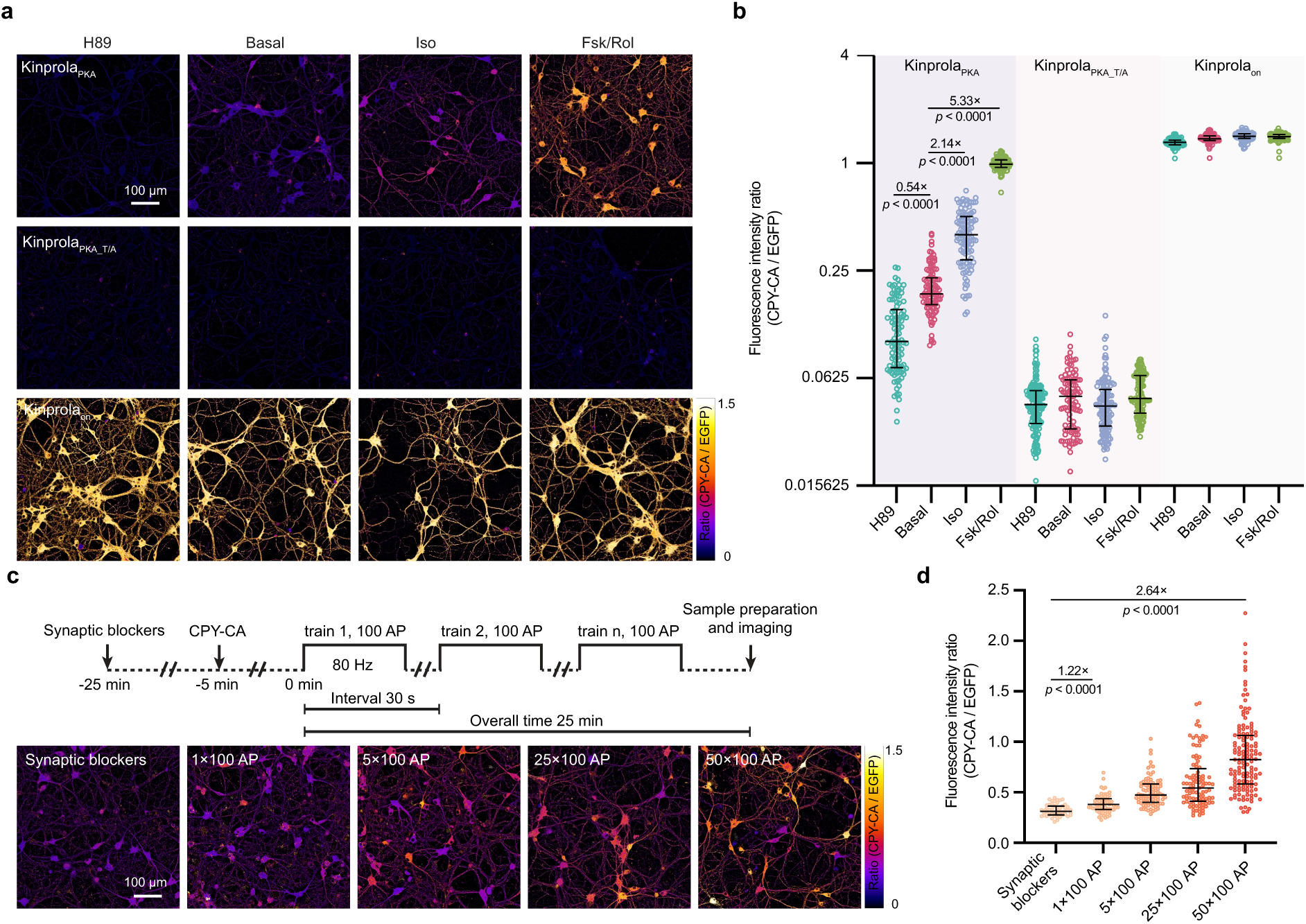
| Kinprola enables rapid recording of PKA activation in cultured neurons. (**a**) Ratiometric fluorescence (CPY-CA/EGFP) images of primary rat hippocampal neurons expressing Kinprola_PKA_, Kinprola_PKA_T/A_ and Kinprola_on_ labeled with CPY-CA (25 nM, 45 min) in the presence of H89, Iso, Fsk/Rol or vehicle. Representative images from three independent experiments with duplicates. (**b**) Dot plot comparison of normalized fluorescence intensity described in (**a**). n ≥ 91 neurons per group, and error bars indicate median with interquartile range. (**c**) Ratiometric fluorescence (CPY-CA/EGFP) images of primary rat hippocampal neurons expressing Kinprola_PKA_ labeled with 125 nM CPY-CA upon defined electrical field stimulation. Representative images from three independent experiments with duplicates. (**d**) Dot plots comparison of normalized fluorescence intensity described in (**c**). n ≥ 51 neurons per group. Error bars indicate median with interquartile range. Statistical significance was calculated with unpaired two-tailed Welch’s *t* test (**b**,**d**), and *p* values are given for comparison. Scale bars: 100 μm (**a**,**c**).

Next, we characterized the performance of Kinprola_PKA_ in acute mouse brain slices (Fig. 5a). Adeno-associated viruses (AAVs) with transgenes of Kinprola_PKA_ or its T/A mutant were stereotactically injected into bilateral nucleus accumbens (NAc) of mice, a striatal region that receives extensive dopaminergic input. Two weeks post-injection, acute brain slices were prepared, perfused with Fsk/Rol to evoke PKA activity and Kinprola_PKA_ labeling was followed through time-lapse fluorescence microscopy. Rapid fluorescence elevation in CPY-CA channel was observed in Fsk/Rol-perfused slices expressing Kinprola_PKA_, whereas the fluorescence changes were significantly smaller in basally active slices (Fig. 5b,c,e). In contrast, no obvious response was detected in slices expressing the T/A mutant whether perfused with Fsk/Rol or vehicle (Fig. 5b,c,e). These experiments demonstrate rapid and specific Kinprola_PKA_ labeling upon PKA activation in a complex and near-native context. After real-time recording, free CPY-CA was washed out and the slices were fixed and mounted. The distinct labeling differences between Fsk/Rol-perfused and vehicle-perfused slices were well preserved in the post hoc imaging and even enhanced relative to those observed in live-cell imaging as washing out free CPY-CA reduced background fluorescence (Fig. 5d,f).

**Fig.5.**
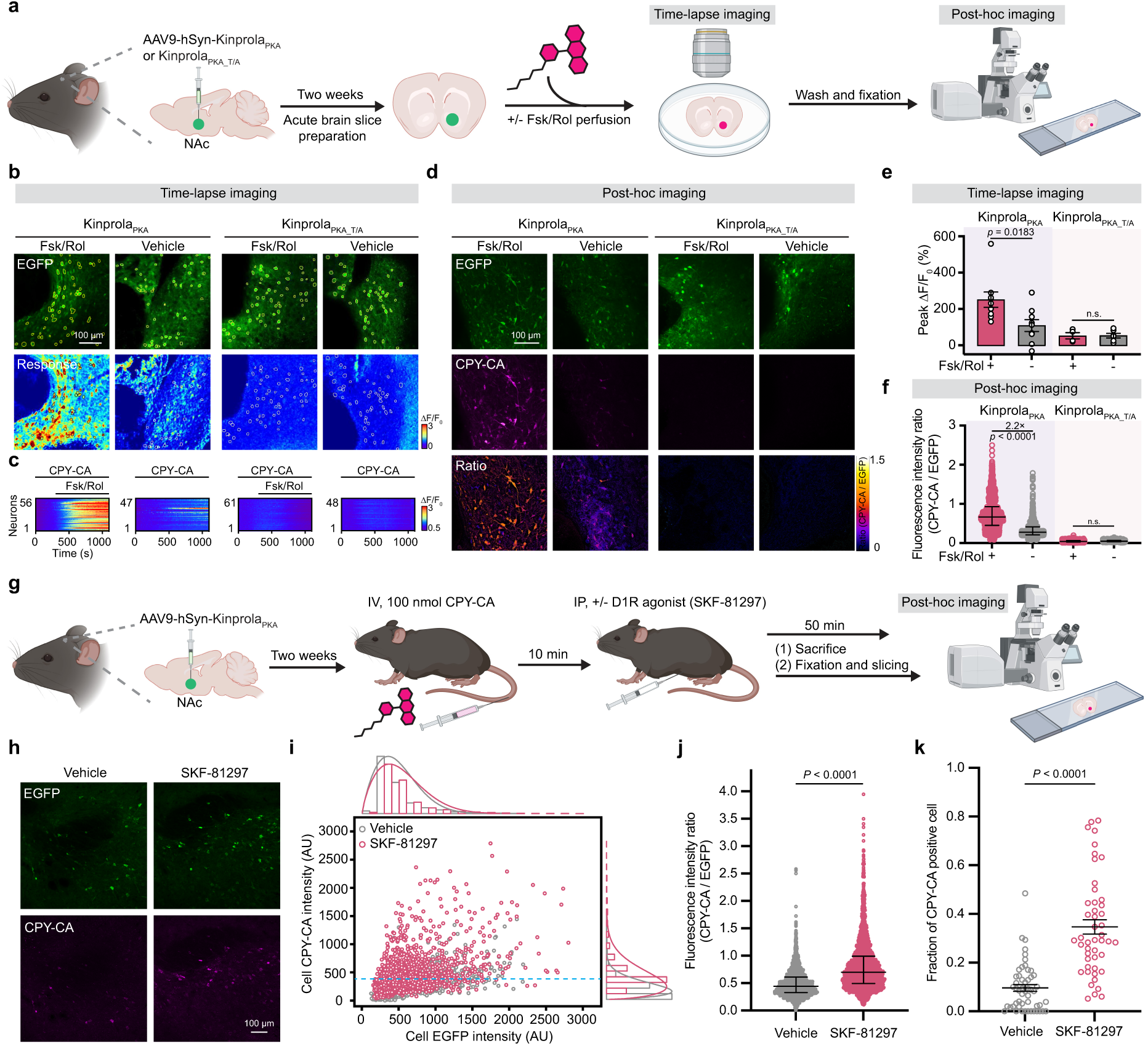
| Kinprola records neuromodulation-induced PKA activation in acute mouse brain slices and freely moving mice. (**a**) Schematic diagram for panels **b**-**f** depicting the experimental design of Kinprola recording pharmacologically induced PKA activation in the nucleus accumbens (NAc) of acute mouse brain slices. (**b**,**c**) Time-lapse imaging of acute mouse brain slices expressing Kinprola_PKA_ or Kinprola_PKA_T/A_ during Fsk/Rol-induced PKA activation. The slices were first perfused with 250 nM CPY-CA to get a baseline, then perfused with 50 μM Fsk/1 μM Rol or vehicle in the presence of 250 nM CPY-CA. Fluorescence intensity traces of single neurons, segmented by circles based on Kinprola expression in each slice shown in (**b**), are presented in (**c**). (**d**) Representative fluorescence images of fixed mouse brain slices expressing Kinprola_PKA_ or Kinprola_PKA_T/A_ perfused under time-lapse imaging conditions in (**b**,**c**). (**e**) Summary of the peak change in CPY-CA fluorescence intensity during time-lapse imaging in (**b**,**c**). n = 9 slices from 5 mice (9/5), 9/5, 4/3 and 6/3 for Fsk/Rol-treated Kinprola_PKA_, vehicle-treated Kinprola_PKA_, Fsk/Rol-treated Kinprola_PKA_T/A_, and vehicle-treated Kinprola_PKA_T/A_. Error bars indicate mean ± SEM. (**f**) Dot plot comparison of fluorescence intensity ratios (CPY-CA/EGFP) of single neurons described in (**d**). n = 1045, 1294, 438 and 614 neurons for Fsk/Rol-treated Kinprola_PKA_, vehicle-treated Kinprola_PKA_, Fsk/Rol-treated Kinprola_PKA_T/A_, and vehicle-treated Kinprola_PKA_T/A_. Error bars indicate median with interquartile range. (**g**) Schematic diagram for panels **h**-**k** depicting the experimental design of Kinprola_PKA_ recording neuromodulation-induced PKA activation during D1/D5 receptor agonist (SKF-81297) treatment *in vivo*. (**h**) Representative fluorescence images of NAc region expressing Kinprola_PKA_ labeled with CPY-CA after SKF-81297 or vehicle injection. (**i**) Scatter plot of mean CPY-CA *vs*. EGFP fluorescence for single neurons expressing Kinprola_PKA_ segmented in SKF-81297-injected mice (n = 2512 neurons in 47 slices from 4 mice) or vehicle-injected mice (n = 2593 neurons in 52 slices from 4 mice). The horizontal dashed line indicates the 90th percentile threshold value of all CPY-CA neurons in the vehicle group. (**j**) Dot plot comparison of normalized fluorescence intensity (CPY-CA/EGFP) of single neurons described in (**i**). n = 2593 neurons pooled from 4 mice for vehicle group, and n = 2512 neurons pooled from 4 mice for SKF-81297 group. Error bars indicate median with interquartile range. (**k**) Dot plot comparison of fraction of all EGFP positive neurons that are CPY-CA positive described in (**i**). (Defined as having a CPY-CA fluorescence value greater than the 90th percentile of neurons in the vehicle group). n = neurons in 52 slices from 4 mice for vehicle group, and neurons in 47 slices from 4 mice for SKF-81297 group. Error bars indicate mean ± SEM. Statistical significance was calculated with one-way ANOVA with Tukey’s Post hoc test (**e**,**f**), or unpaired two-tailed Welch’s *t* test (**j**,**k**), and *p* values are given for comparison. Scale bars: 100 μm (**b**,**d**,**h**).

Finally, we applied Kinprola_PKA_ to record neuromodulation-induced PKA activity changes in the NAc of freely moving mice. Dopaminergic signaling through abundant type 1 dopamine receptors (D1Rs) in the NAc can activate PKA and subsequently modulate a multitude of brain functions, including reward signaling and reinforcement learning^3,10,11,39^. To assess whether Kinprola_PKA_ can record PKA activity in response to D1R activation, mice expressing Kinprola_PKA_ first received an intravenous tail vein injection (IV) of CPY-CA. After 10 minutes, SKF-81297, a potent and selective D1/D5R agonist known to increase signaling via the cAMP/PKA pathway^8,10,11,40,41^ was administered via intraperitoneal (IP) injection. 50 minutes later, mice were sacrificed and NAc slices were processed for post hoc imaging (Fig. 5g). The normalized fluorescence intensity of single neurons expressing Kinprola_PKA_ significantly increased following SKF-81297 injection compared to the fluorescence labeling observed after vehicle injection (Fig. 5h-k and Extended Data Fig. 10). In SKF-81297-injected mice, around 35% neurons expressing Kinprola_PKA_ per slices in the NAc exhibited strong CPY-CA labeling, compared to around 10% after injection of vehicle (Fig. 5k). These experiments demonstrate the ability of Kinprola_PKA_ to directly and rapidly record PKA activity for monitoring neuromodulation with cellular resolution *in vivo* for later analysis in deep tissues of freely moving mice, providing a scalable and complementary strategy to current real-time PKA activity biosensors.

## Discussion

Genetically encoded fluorescent kinase activity reporters have been extensively utilized to monitor kinase activities in real-time for elucidating the connections between intracellular signaling cascades and neuromodulation in the nervous system. However, such reporters face limitations when challenged to record kinase activities with high spatiotemporal resolution in a scalable manner or in deep tissues of freely moving animals. These limitations are mainly due to the inherent constraints of light microscopy. A strategy that potentially addresses these limitations would involve separating the recording period from its analysis by converting transient protein kinase activities into a “permanent” mark for later analysis. To achieve this, we developed Kinprola, a chemigenetic approach for recording protein kinase activity based on our recently reported split-HaloTag system^17^. Kinprola accumulates an irreversible fluorescent mark in the presence of both a specific protein kinase activity and a fluorescent substrate. Thus, the light delivery required for monitoring the activity with fluorescent biosensors is replaced by the delivery of a fluorescent substrate. The recording window is time-gated by applying or washing out the fluorescence substrate, typically spanning from a few minutes to hours. Kinprola rapidly responds to the cellular phosphorylation state, enabling successive recordings of kinase activity during different periods through the use of spectrally distinct substrates.

The design principle of Kinprola was inspired by the classical kinase activity reporters, in which the reversible binding of a phosphorylated substrate peptide to a specific phosphoamino acid binding domain triggers proximity between two tethered fluorescent proteins forming a FRET pair^15^. In Kinprola, these two fluorescent proteins are replaced by the two components of a split-HaloTag system. Phosphorylation of Kinprola then leads to reconstitution of active HaloTag, which in the presence of a fluorescent HaloTag substrate leads to its irreversible labeling. Using this modular design principle, we have generated recorders for PKA, PKC, JNK and AMPK. Kinprola may be further extended to record the activity of other types of kinases, such as extracellular signal-regulated kinase (ERK) and tyrosine kinases, by using appropriate substrate peptides and phosphoamino acid binding domains.

The large majority of the experiments reported here were performed with Kinprola_PKA_. A key feature of Kinprola_PKA_ is the persistence of the fluorescent mark, which remains detectable for days in live neurons and is resistant to fixation procedures. This allows the sorting of cells based on their kinase activity histories from large and heterogeneous cell populations for later analysis. Transcriptome analysis of GBCs subpopulations sorted according to Kinprola labeling highlights the important role of PKA in cell cycle regulation, which potentially correlates with glioblastoma proliferation and invasion. Furthermore, a Kinprola_PKA_-based CRISPR knockout screening identified numerous putative non-canonical regulators that influence PKA activity, potentially broadening our understanding of PKA signaling and aiding in therapeutic target identification. Of note, using different cell sorting strategies, the putative hits identified by CRISPR knockout screening in RKO cells did not significantly correlate with DEGs identified by RNAseq in GBCs, where no genetic perturbation was introduced. This demonstrates that Kinprola recording can be tailored to address various biological questions by adjusting the recording parameters and experimental context.

Kinprola_PKA_ can record cytosolic PKA activation on a minute scale during pharmacological or electrophysiological stimulation in primary neurons and acute brain slices. Its sensitivity and temporal resolution under these conditions approximately align with the timescale of endogenous PKA signaling, estimated from the dynamics of PKA sensors and the cAMP dissociation rate from PKA regulatory subunits (K_d_ around 0.15 min^-1^)^9,42,43^. Kinprola is also a promising tool for recording PKA activity *in vivo* as demonstrated by the labeling of Kinprola_PKA_ in NAc neurons with elevated PKA activity provoked by a selective D1/D5R agonist in freely moving mice. The fast *in vivo* clearance kinetics of fluorescent HaloTag substrates in mice (t_1/2_ around 15 min) restricts the recording period of Kinprola_PKA_ after a single injection of CPY-CA to minute timescales^44,45^. Extending the recording period of Kinprola *in vivo* would thus require either different delivery methods of the HaloTag substrates or the development of HaloTag substrates with more favorable pharmacokinetic properties.

PKA activity has been monitored as a central surrogate for intracellular neuromodulation throughout the nervous system, Kinprola should thus become a complementary tool to existing fluorescent PKA activity reporters for dissecting mechanisms related to neuromodulatory activity *ex vivo* and *in vivo*. In the future, it should be possible to stably mark neurons with elevated PKA activity using Kinprola in the nervous system for transcriptome analysis and pharmacological or genetic screenings, enabling systematic studies of the molecular features of signaling pathways during neuromodulation.

## Supporting information

Sun et al_Kinprola_Supplemental Files

## Acknowledgements

We are grateful to the Flow Cytometry Core Facility of the German Cancer Research Center (DKFZ, Heidelberg) and the Genomics and Proteomics Core Facility of the DKFZ for support, C-Tech Deep Sequencing Core Facility at BioQuant (Heidelberg University) for sequencing, and the Omics IT and Data Management Core Facility of the DKFZ for processing RNA-Seq data and data storage. We thank J. Hiblot for discussions; A. Andres-Pons (European Molecular Biology Laboratory, Heidelberg) for providing the HeLa Kyoto Flp-In cell line; L. D. Lavis (Janelia Research Campus, Ashburn, Virginia) for providing Janelia Fluor dyes; A. Bergner, J. Kress, B. Koch, B. Réssy, D. Schmidt and E. D’Este for reagents or materials; S. Wendler for maintaining glioblastoma stable cells. We thank the mass spectrometry facility (S. Fabritz, T. Rudi, and J. Kling) of the MPIMR for its support, and M. Tarnawski for nanoDSF measurement. We are grateful to B. Luo and S. Li for assistance with mice experiments. This work was funded by the Max Planck Society (D.S., N.S., P.B., K.J.), École Polytechnique Fédérale de Lausanne (EPFL) (K.J.), Deutsche Forschungsgemeinschaft (DFG) SFB TRR 186 (K.J.), GRK 2099 (S.N.), SFB 1324 (M.B.), the National Natural Science Foundation of China (31925017, Y.L.), and the New Cornerstone Science Foundation through the New Cornerstone Investigator Program (Y.L.). D.S. was supported by a Humboldt Research Fellowship and an interinstitutional postdoctoral fellowship of The Health + Life Science Alliance Heidelberg Mannheim.

## Author contributions

K.J. and D.S. conceived the project. D.S. conducted most of the experiments unless specified otherwise. D.S., N.S. and P.B. together designed and performed the mammalian cell experiments. D.S., S.N., K.E.B. and M.B. together designed and performed the CRISPR screen. D.S., D.C.H., J.M., D.D.A., F.W. and W.W. together designed and performed the RNA-Seq. D.S., Y.Z., S.X. and Y.L. together designed and performed mice-related experiments. D.S. and K.J. wrote the manuscript with input from all authors.

## Data availability

All data are available in the paper or the supplementary materials. Plasmids of interest from the study will be deposited at Addgene; accession codes will be provided in the supplementary materials. Raw RNA-Seq data has been deposited in NCBI’s Gene Expression Omnibus^46^ and are accessible through GEO Series accession number GSE269419 (https://www.ncbi.nlm.nih.gov/geo/query/acc.cgi?acc=GSE269419). Raw sequencing data of CRISPR screen will be available from the GEO. Reagents and materials are available from the corresponding authors upon request. depicting the process for developing the Kinprola_PKA_ recorder. PAABD: phosphoamino acid binding domain. PKAsub: PKA substrate peptide. Hpep: Halo peptide. (**b**) Amino acid sequence of the Kinprola_PKA_ recorder, with the targeting sequence, domains, linkers, phosphorylation site and the mutated site indicated.

---

**Extended Data Fig. 1.**
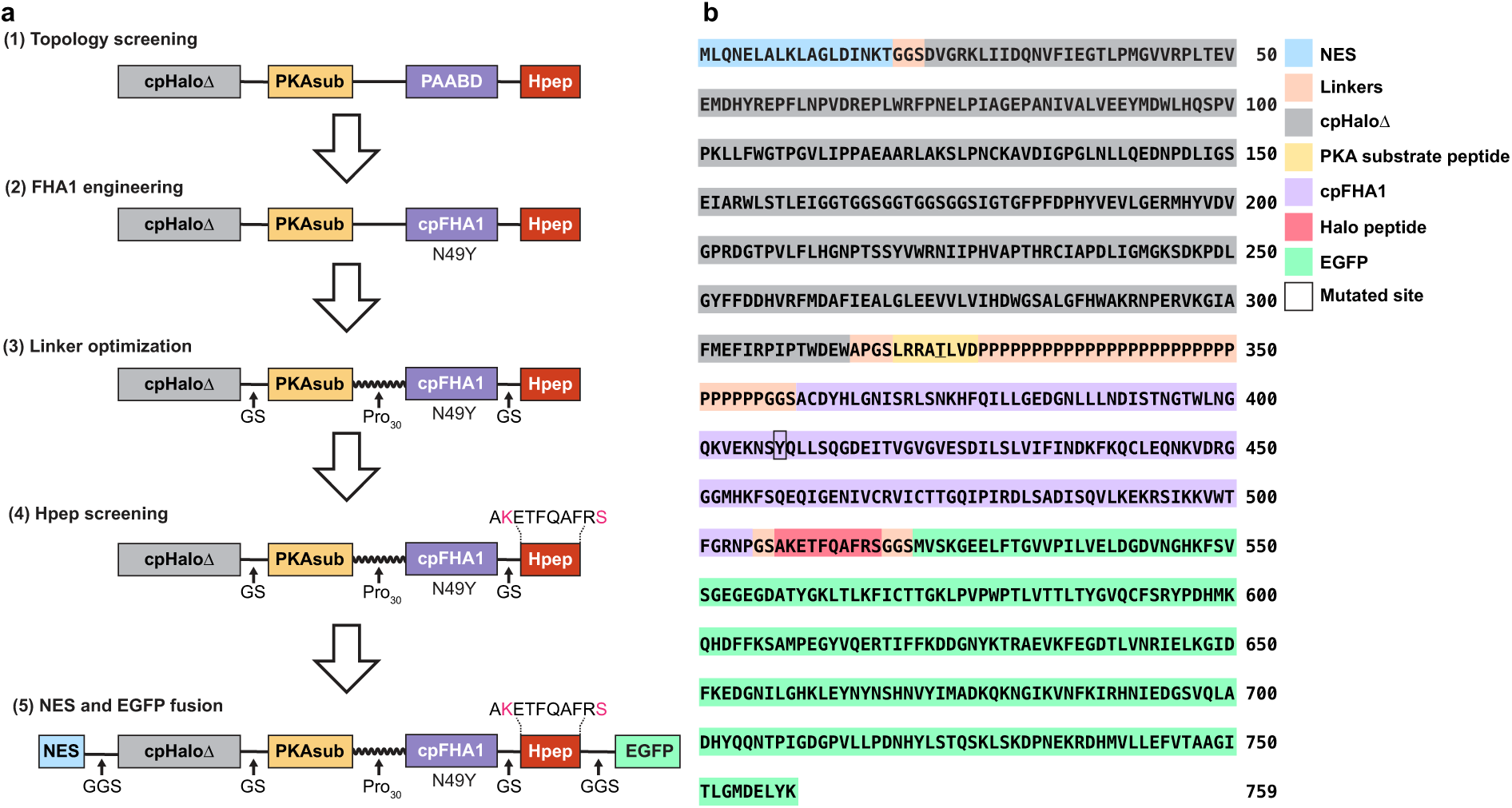
| Strategy for designing, optimizing, and screening Kinprola. (**a**) Flowchart depicting the process for developing the Kinprola_PKA_ recorder. PAABD: phosphoamino acid binding domain. PKAsub: PKA substrate peptide. Hpep: Halo peptide. (**b**) Amino acid sequence of the Kinprola_PKA_ recorder, with the targeting sequence, domains, linkers, phosphorylation site and the mutated site indicated.

**Extended Data Fig. 2.**
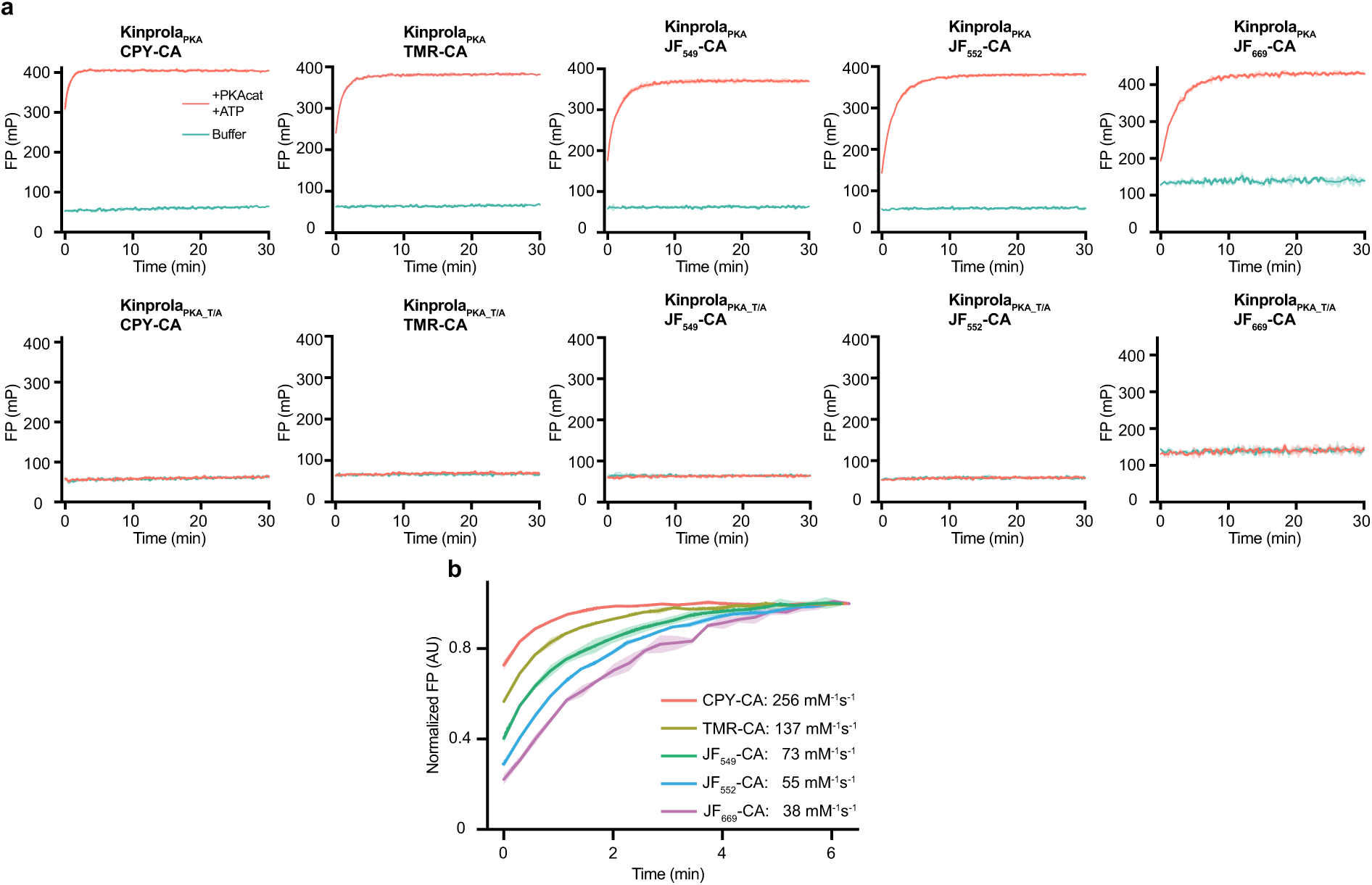
| (related to Fig. 1) Labeling kinetics of Kinprola with different fluorescent HaloTag substrates. (**a**) Labeling kinetics of Kinprola_PKA_ and Kinprola_PKA_T/A_ (200 nM) with different fluorescent HaloTag substrates (50 nM) in the presence or absence of PKAcat (25 ng μL^-1^) and ATP (500 μM), measured by fluorescence polarization (FP). Labeling kinetics with TMR-CA is represented in **Fig. 1b**. (**b**) Comparison of labeling kinetics between different fluorescent HaloTag substrates described in (**a**). Fluorescence polarization values were normalized to their unbound and fully bound values (normalized FP). AU, arbitrary units. Data from three technical replicates.

**Extended Data Fig. 3.**
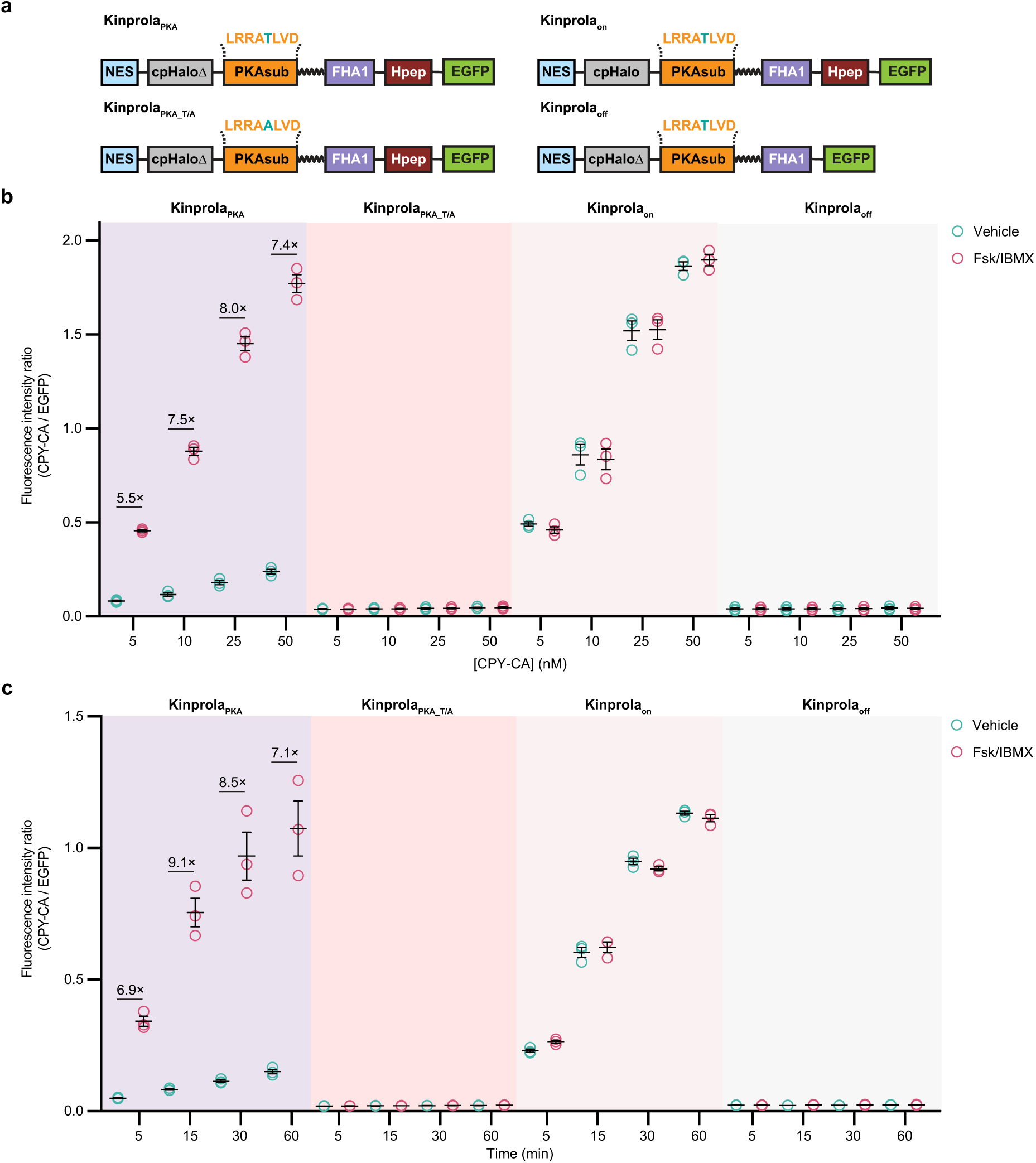
| (related to Fig. 1) Time and fluorescent substrate-dependency of Kinprola labeling. (**a**) Domain structures of Kinprola_PKA_, Kinprola_PKA_T/A_, Kinprola_on_, and Kinprola_off_. (**b**) Flow cytometry analysis of Kinprola labeling at different CPY-CA concentrations in the presence or absence of 50 μM Fsk/100 μM IBMX stimulation for 30 min. Fluorescence intensity ratios (CPY-CA/EGFP) of CPY-labeled HeLa cells stably expressing Kinprola_PKA_, Kinprola_PKA_T/A,_ Kinprola_on_ and Kinprola_off_ are shown. Error bars indicate mean ± SEM (from three independent experiments with triplicates). (**c**) Flow cytometry analysis of Kinprola labeling with 25 nM CPY-CA over different time periods in the presence or absence of 50 μM Fsk/100 μM IBMX stimulation. Fluorescence intensity ratios (CPY-CA/EGFP) of CPY-labeled HeLa cells stably expressing Kinprola_PKA_, Kinprola_PKA_T/A,_ Kinprola_on_ and Kinprola_off_ are shown. Error bars indicate mean ± SEM (from three independent experiments with triplicates).

**Extended Data Fig. 4.**
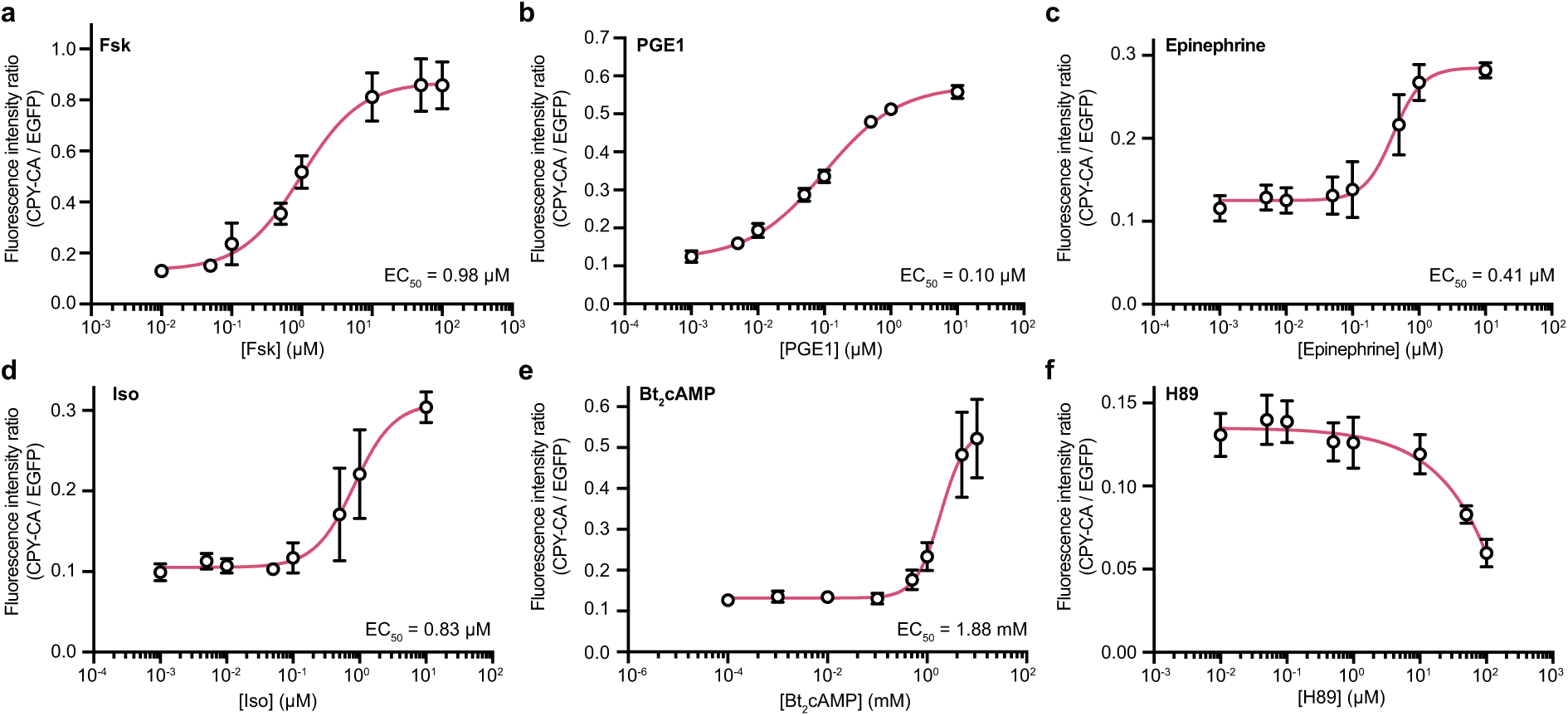
| (related to Fig. 1) Labeling of HEK293 cells stably expressing Kinprola_PKA_ in the presence of varying concentrations of different PKA modulators. Cells were labeled with 25 nM CPY-CA for 30 min in the presence of PKA with different concentration and subsequently analyzed by flow cytometry. Data are shown as mean ± SEM (from three independent experiments with replicates). Curves were fitted with the sigmoidal function to determine EC_50_ values. Dose curve of Fsk is represented in **Fig. 1f**.

**Extended Data Fig. 5.**
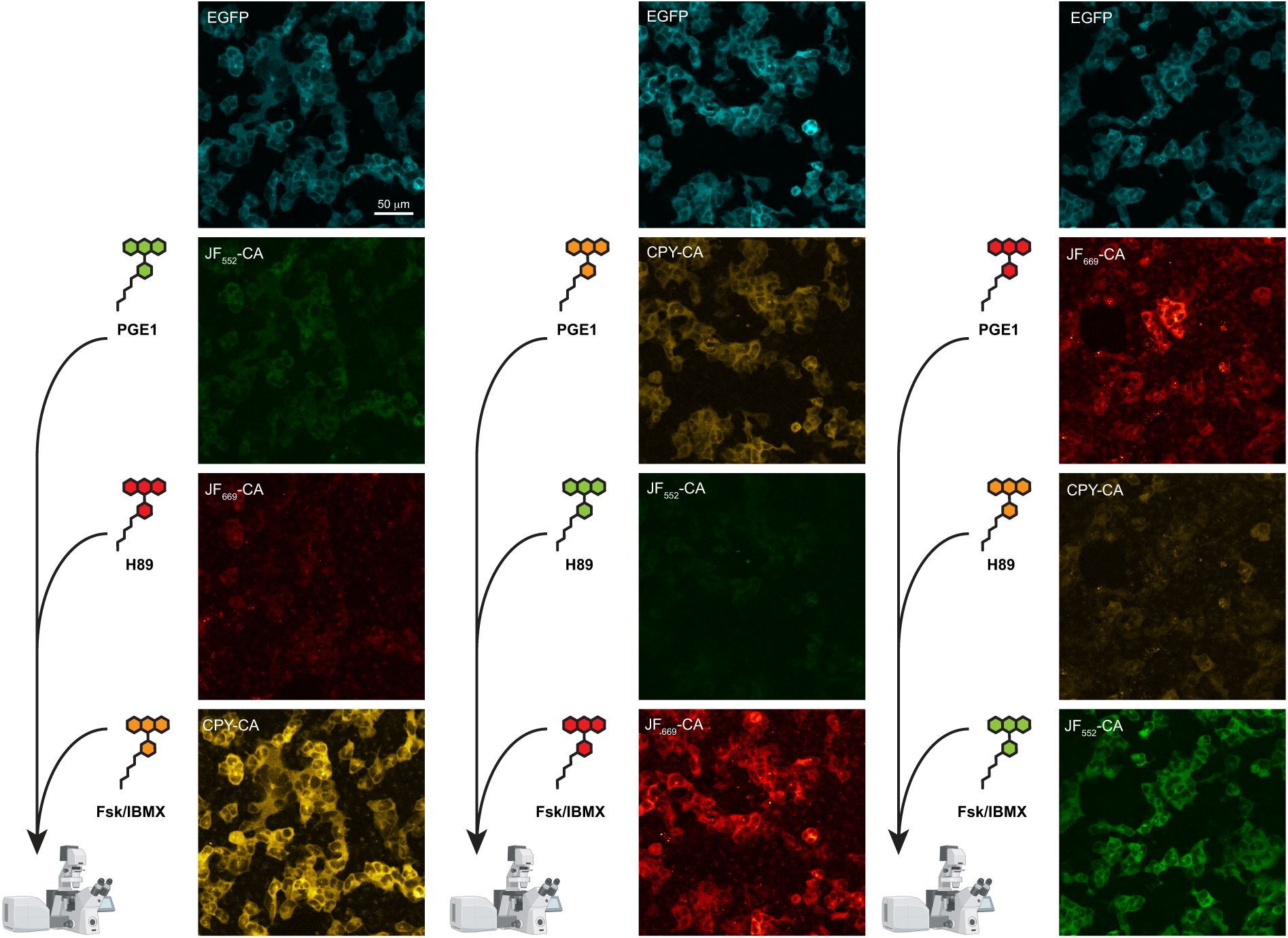
| (related to Fig. 1) Recording of three successive periods of PKA activity in HEK293 cells stably expressing KinprolaPKA. Fluorescence images depicting the successive recordings of PKA activity in Kinprola_PKA_-expressing HEK293 cells. First treatment: PGE1 stimulation; second treatment: H89 inhibition; third treatment: Fsk/IBMX stimulation, in the presence of JF_552_-CA or JF_669_-CA or CPY-CA. Cells were allowed to rest for 2 h between each treatment. Imaging was performed after the third incubation period. The extent of labeling is proportional to the drug treatment, independent of the fluorescent substrate. Labeling degree: H89 inhibition < PGE1 stimulation < Fsk/IBMX stimulation. Representative images from three independent experiments. Partial images are represented in **Fig. 1h**. Scale bar: 50 μm.

**Extended Data Fig. 6.**
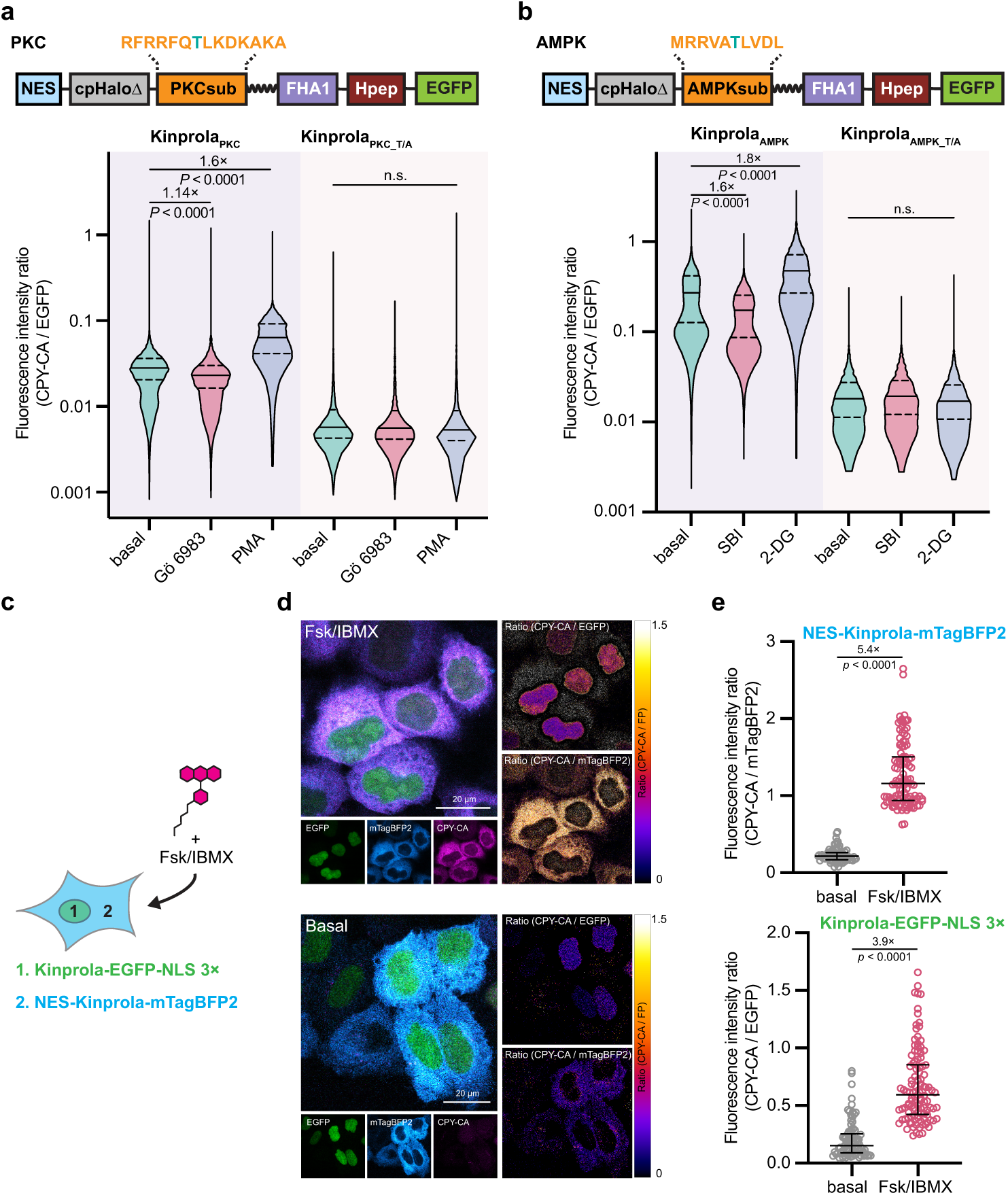
| (related to Fig. 1) Kinprola for recording the activities of different kinases and compartmentalized PKA activity in different subcellular locations. (**a**,**b**) Flow cytometry analysis of Kinprolas labeling with 25 nM CPY-CA in the presence of corresponding activators or inhibitors for 30 min in HEK293 cells. Error bars indicate median with interquartile range (from three independent experiments with triplicates). (**c**) Schematic diagram depicting Kinprola_PKA_ multiplexed recording in different cellular localization. (**d**) Fluorescence images of Hela cells expressing Kinprola_PKA_ in nucleus and cytoplasm by adeno-associated viruses (AAVs) transduction, with or without Fsk/IBMX stimulation in the presence of 50 nM CPY-CA for 30 min. Representative images from three independent experiments with replicates. Scale bars: 20 μm. (**e**) Dot plot comparison of normalized fluorescence intensities (CPY-CA/mTagBFP2) and (CPY-CA/EGFP) described in (**d**). Error bars indicate median with interquartile range. Statistical significance was calculated with one-way ANOVA with Dunnett’s Post hoc test (**a**,**b**) or unpaired two-tailed Welch’s *t* test (**e**) and *p* values are given for comparison.

**Extended Data Fig. 7.**
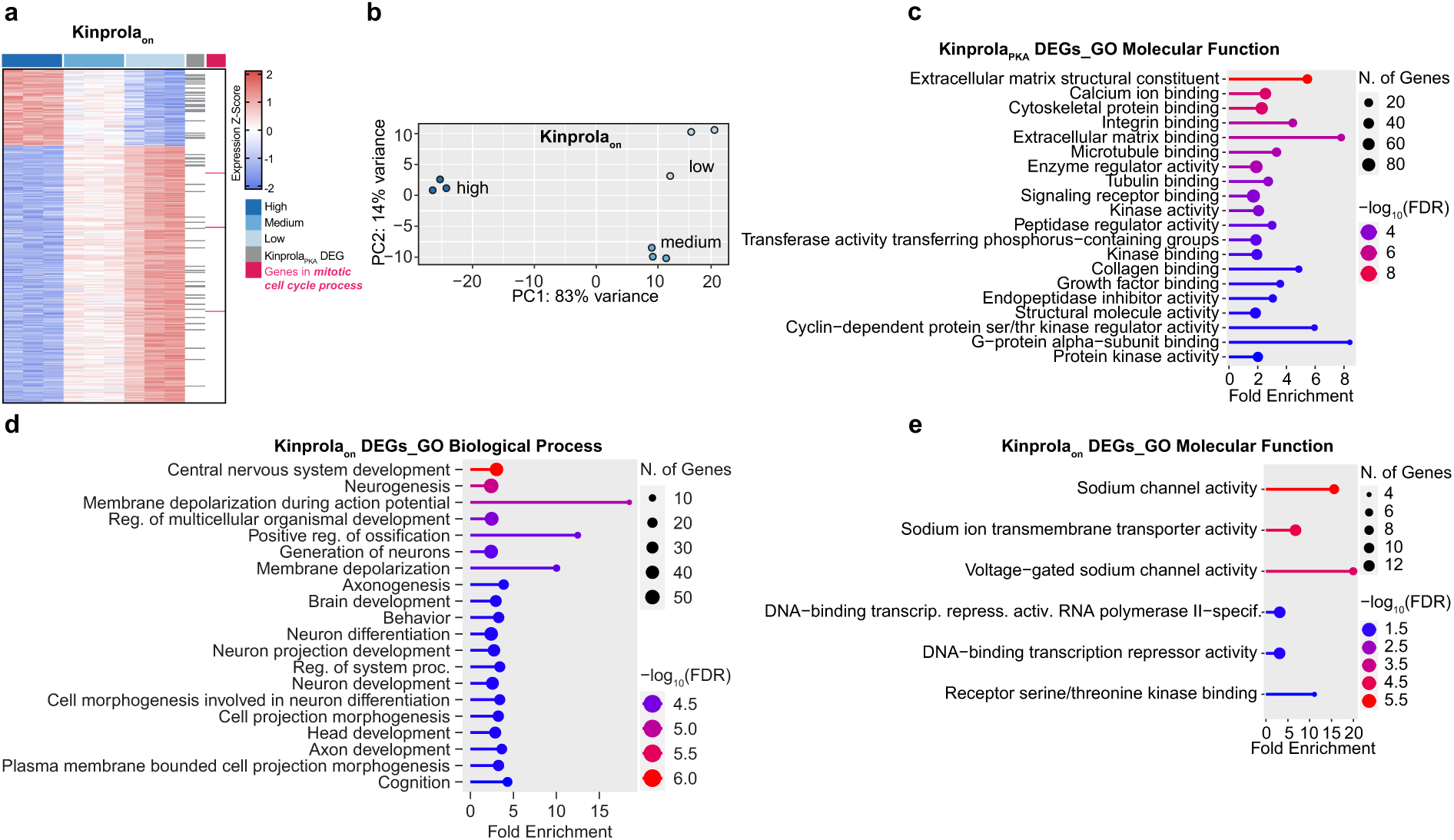
| (related to Fig. 2) Transcriptome analysis of GBC subpopulations labeled and selected by Kinprola. (**a**) Transcriptional profiles of the three GBC subpopulations selected by Kinprola_on_ labeling. DEGs identified by RNA-Seq analysis are color coded according to the Z-score. Genes in the GO term “mitotic cell cycle process” are highlighted in magenta, and overlapped DEGs identified in both Kinprola_on_ and Kinprola_PKA_ experiments are indicated in gray. (**b**) Dot plots of the first principal coordinate analysis on the three sorted groups of Kinprola_on_-expressing GBCs. (**c**) Molecular function GO terms of Kinprola_PKA_-identified DEGs. (**d**) Biological process Gene Ontology (GO) terms of Kinprola_on_-identified DEGs. (**e**) Molecular function GO terms of Kinprola_on_-identified DEGs.

**Extended Data Fig. 8.**
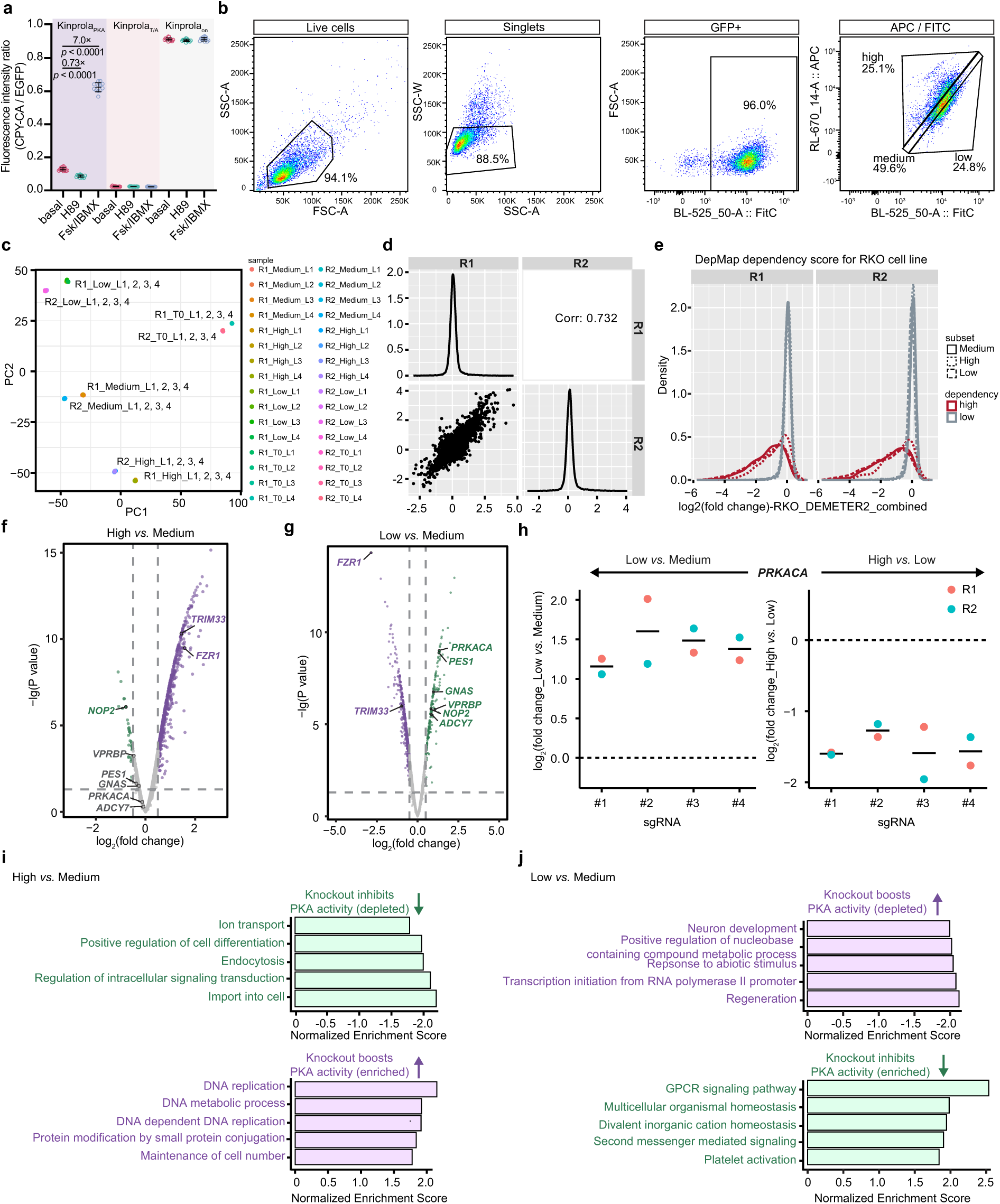
| (related to Fig. 3) Pooled CRISPR knockout screen analysis of RKO cell subpopulations sorted based on Kinprola_PKA_ labeling. (**a**) Fluorescence intensity ratios (CPY-CA/EGFP) quantification of RKO cells stably expressing Kinprola_PKA_, Kinprola_PKA_T/A_, and Kinprola_on_ labeled with CPY-CA (125 nM CPY-CA, 30 min) in the presence of 20 μM H89 inhibition, 50 μM Fsk/100 μM IBMX stimulation or vehicle. Statistical significance was calculated with unpaired two-tailed Welch’s *t* test, and *p* values are given for comparison. Error bars indicate mean ± SEM (from three independent experiments with triplicates). (**b**) Gating and sorting strategies of CPY-CA labeled RKO cells stably expressing Kinprola_PKA_ for pooled CRISPR knockout screen. (**c**) Principal component analysis of different sorted populations in two independent replicate screens (R1, R2). (**d**) Gene-level reproducibility analysis of two independent replicate screens (R1, R2). (**e**) Core essential genes (high dependency) were strongly depleted over the course of screening with HD CRISPR sub-library A, compared to nonessential genes (low dependency). Core-essential genes of RKO cells with dependency score were obtained from the Cancer Dependency Map Project (DepMap). (**f**)-(**g**) Volcano plots showing gene enrichment by comparing “high”, “medium” and “low” subpopulations sorted in (**b**). Dashed lines indicate cutoff for hit genes (FDR 0.05). Putative hits and canonical regulators including *PRKACA*, *GNAS*, and *ADCY7* are highlighted. (**h**) Enrichment comparison of four sgRNAs targeting *PRKACA* in different sorted groups. All four sgRNAs were enriched in the “low” group and depleted in the “high” group of two independent replicate screens (R1, R2). (**i**) Gene set enrichment analysis (GSEA) of the top 5 categories among “high” *vs.* “medium” comparison. (**j**) GSEA top 5 categories among “low” *vs.* “medium” comparison.

**Extended Data Fig. 9.**
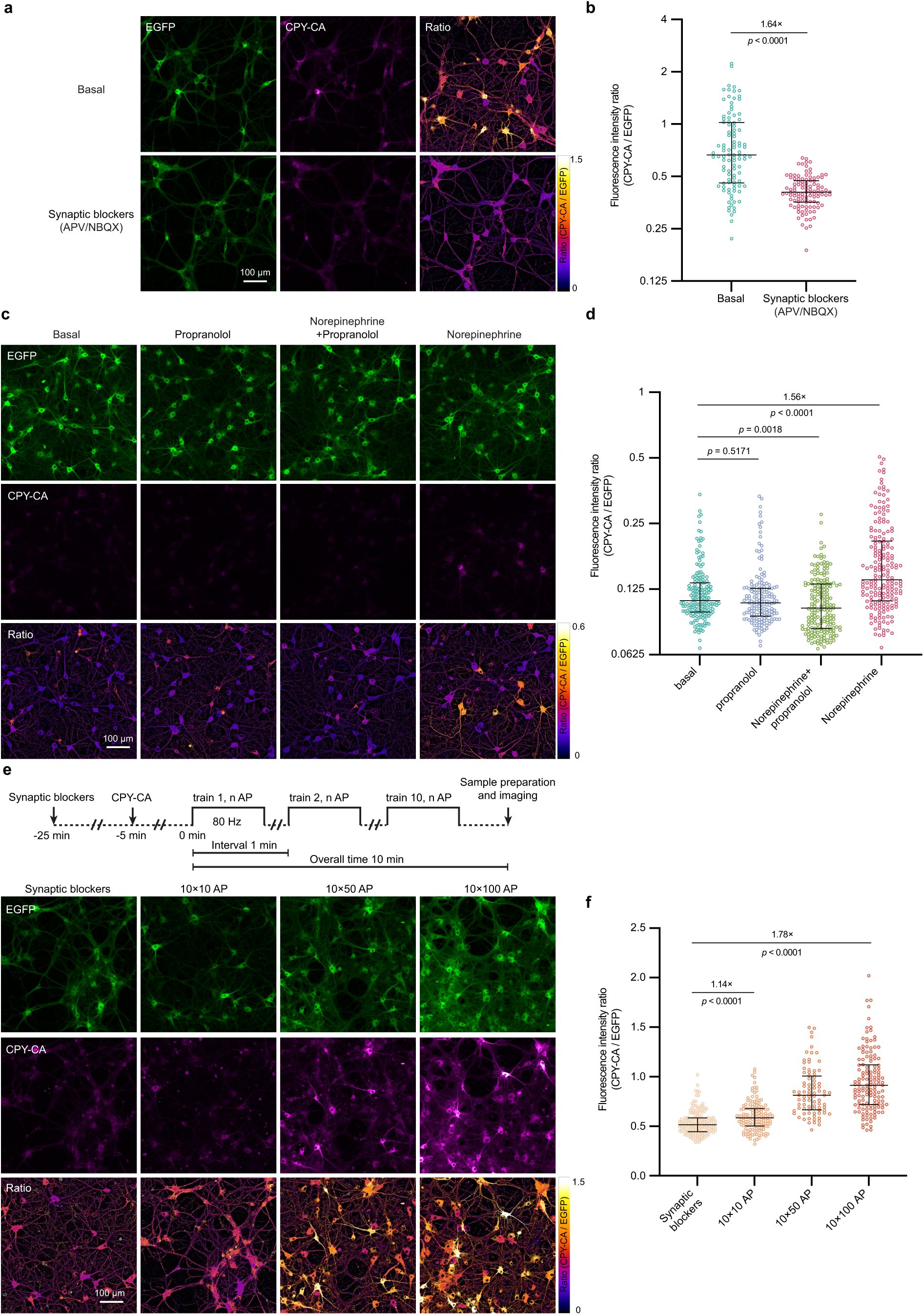
| (related to Fig. 4) Recording PKA activity in primary hippocampal neurons expressing Kinprola_PKA_ during pharmacological treatment and electrophysiological stimulation. (**a**) Fluorescence images of primary rat hippocampal neurons expressing Kinprola_PKA_ labeled with CPY-CA (125 nM, 45 min) in the presence of synaptic blockers 25 μM APV/10 μM NBQX (25 μM/10 μM) or vehicle. The ratio indicates CPY-CA/EGFP. Representative images from three independent experiments with duplicates. (**b**) Quantification of Kinprola_PKA_ labeling from experiments described in (**a**) (n ≥ 101 cells). (**c**) Fluorescence images of primary rat hippocampal neurons expressing Kinprola_PKA_ labeled with CPY-CA (25 nM, 45 min) in the presence of 1 μM norepinephrine, 1 μM propranolol, 1 μM norepinephrine/1 μM propranolol or vehicle. The ratio indicates CPY-CA/EGFP. Representative images from three independent experiments with duplicates. (**d**) Quantification of Kinprola_PKA_ labeling from experiments described in (**c**) (n ≥ 168 cells). (**e**) Fluorescence images of primary rat hippocampal neurons expressing Kinprola_PKA_ labeled with 125 nM CPY-CA upon defined electrical field stimulation. Representative images from three independent experiments with duplicates. (**f**) Dot plots comparison of normalized fluorescence intensity described in (**e**). n ≥ 84 neurons per group. Error bars indicate median with interquartile range (**b**,**d**,**f**). Statistical significance was calculated with unpaired two-tailed Welch’s *t* test (**b**,**d**,**f**), and *p* values are given for comparison. Scale bars: 100 μm (**a**,**c**,**e**).

**Extended Data Fig. 10.**
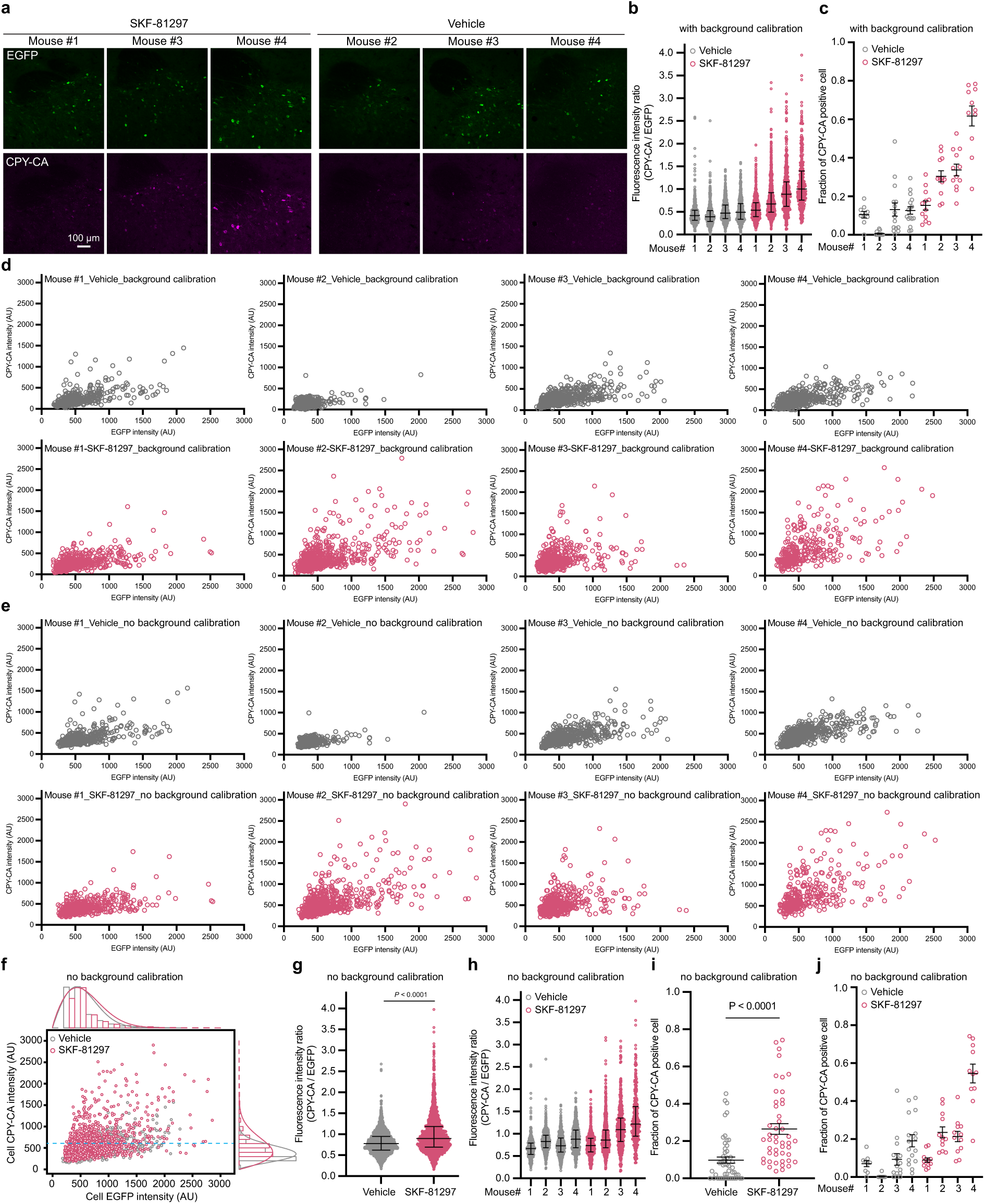
| (related to Fig. 5) Kinprola_PKA_ records neuromodulation-induced PKA activation in freely moving mice. (**a**) Additional representative fluorescence images of the nucleus accumbens (NAc) region expressing Kinprola_PKA_ labeled with CPY-CA after SKF-81297 or vehicle injection from another 3 mice. (**b**) Dot plot comparison of fluorescence intensity ratios (CPY-CA/EGFP) of neurons per mouse described in **Fig. 5j**. Error bars indicate median with interquartile range. (**c**) Dot plot comparison of fraction of all EGFP-positive neurons that are CPY-CA positive of a single slice described in **Fig. 5k**. (**d**) Scatter plot of mean CPY-CA *vs.* EGFP fluorescence for single EGFP-positive neurons segmented in individual SKF-81297-injected or vehicle-injected mouse described in **Fig. 5i**. AU, arbitrary units. (**e**) Scatter plot of mean CPY-CA *vs.* EGFP fluorescence for single EGFP-positive neurons segmented in individual SKF-81297-injected or vehicle-injected mouse, raw intensity value without background calibration. (**f**) Scatter plot of mean CPY-CA *vs.* EGFP fluorescence for single EGFP-positive neurons segmented in all SKF-81297-injected or vehicle-injected mice described in **Fig. 5i**, raw intensity value without background calibration. The horizontal dashed line indicates the 90th percentile threshold value of all CPY-CA neurons in the vehicle group. (**g**) Dot plot comparison of fluorescence intensity ratios (CPY-CA/EGFP) of neurons described in (**f**), raw intensity value without background calibration. Error bars indicate median with interquartile range. (**h**) Dot plot comparison of fluorescence intensity ratios (CPY-CA/EGFP) of neurons in individual mouse described in (**g**). Error bars indicate median with interquartile range. (**i**) Dot plot comparison of fraction of all EGFP positive neurons that are CPY-CA positive described in (**f**). Threshold using the dashed line in (**f**). Error bars indicate mean ± SEM. (**j**) Dot plot comparison of fraction of all EGFP-positive neurons that are CPY-CA positive in slices per mouse described in (**i**). Error bars indicate mean ± SEM. Statistical significance was calculated with unpaired two-tailed Welch’s *t* test (**g**,**i**), and *p* values are given for comparison. Scale bar: 100 μm (**a**).

## Online methods

### General information

All reagents were purchased from commercial suppliers and used without further purification. Detailed information on the reagents and resource is listed in **Supplementary Table 3**. Fluorescent HaloTag substrates were synthesized according to literature procedures^47,48^. Janelia Fluor (JF) dyes were generously provided by L. D. Lavis (Janelia Research Campus, Ashburn, Virginia). Fluorescent substrates were dissolved in dry DMSO to create stock solutions, and subsequently diluted in the respective buffers to ensure the final DMSO concentration was within 1% (vol/vol). The chemical structures and photophysical properties of these substrates are presented in **Supplementary Fig. 1**. For drug treatment, an equal volume of DMSO was used as the vehicle control unless otherwise specified. The composition of common buffers used in this study is detailed in **Supplementary Table 4**.

### Molecular cloning

For protein production in *E. coli*, the pET51b(+) vector (Novagen) was used. For mammalian cell expression, the pcDNA5/FRT vector (ThermoFisher Scientific) was used. Unless otherwise specified, molecular cloning was performed using Gibson assembly^49^, In-Fusion Cloning or the Q5 site-directed mutagenesis kit, following the manufacturer’s protocols. Primers were synthesized by Sigma-Aldrich or Eurofins. DNA amplification was carried out by PCR using KOD-hot-start DNA polymerase master mix or Q5 high-fidelity DNA polymerase. PCR products were purified using the QIAquick PCR purification kit or the NucleoSpin Gel and PCR Clean-up kit. Cloning products generated by Gibson assembly were electroporated into *E. coli* strain 10G. For cloning virus-related plasmids, NEB stable competent *E. coli* and One Shot Stbl3 chemically competent *E. coli* were used. Plasmids were purified with the QIAprep Spin Miniprep Kit. For cell transfection purpose, endotoxin was removed using the GeneJET Endo-Free Plasmid Maxiprep Kit. All the sequences were verified by Sanger sequencing, and the integrity of inverted terminal repeats recombination sites (ITRs) of virus-related plasmids was also verified. All the plasmids used in this study can be found in **Supplementary Table 5**.

### Single sgRNA plasmid cloning

For cloning individual sgRNA sequences into the HDCRISPRv1 vector^36^, the vector was sequentially digested with BfuAI and BsrGI-HF, followed by dephosphorylation using Quick CIP. The digested vector was then purified by gel electrophoresis. Annealed oligos were ligated into digested vector using the Quick Ligation Kit according to the manufacturer’s protocol. All the sgRNA sequences can be found in **Supplementary Table 6**.

### Protein expression and purification

Proteins were expressed in *E. coli* strain BL21(DE3). Lysogenic broth (LB) cultures were grown at 37°C until the optical density at 600 nm (OD 600 nm) reached 0.8. Protein expression was induced by adding 0.5 mM isopropyl-β-D-1-thiogalactopyranoside (IPTG) and the cultures were then grown at 16°C overnight. The cell cultures were harvested by centrifugation (4,500 g, 10 min, 4°C) and lysed by sonication on ice in IMAC lysis buffer (see **Supplementary table 4**). The cell lysate was cleared by centrifugation (70,000 g, 20 min, 4°C). Proteins were purified using HisPur Ni-NTA Superflow Agarose or on the ÄktaPure FPLC system (Cytiva) with IMAC wash and elution buffer (see **Supplementary table 4**) and concentrated using Ultra-15 centrifugal filter units with a molecular weight cut-off smaller than the protein size, followed by buffer exchange into the activity buffer (see **Supplementary table 4**, final imidazole concentration < 0.1 mM). The purification His_10_ tag was removed by overnight cleavage with TEV protease at 30°C as previously described^50^. Cleaved proteins were purified using a HisTrap FF crude column (Cytiva) on the ÄktaPure FPLC system (Cytiva) by collecting the flow-through. The proteins were further purified by size exclusion chromatography (HiLoad 26/600 Superdex75, Cytiva) using the activity buffer. The purified proteins were either flash frozen in liquid nitrogen and stored at −80°C, or mixed 1:1 (vol/vol) with 90% (wt/vol) glycerol in activity buffer and stored at −20°C. The final protein concentration ranged from 100 to 500 μM. The correct size and purity of proteins were assessed by SDS-PAGE and liquid chromatography-mass spectrometry (LC-MS). Protein amino acid sequences are listed in **Supplementary Note 1**.

### *In vitro* phosphorylation of Kinprola_PKA_ protein

In a 500 µL kinase assay buffer system (see **Supplementary table 4**), 10 µM of purified Kinprola_PKA_ protein was mixed with 10 µg of PKAcat and 200 µM of ATP, then incubated at 30°C for 1 h with shaking at 300 rpm. The phosphorylated proteins were purified by size-exclusion chromatography and assessed by LC-MS.

### Protein thermostability measurement

Protein thermostability was measured at 0.5 mg mL^−1^ in activity buffer using a nanoscale differential scanning fluorimeter (NanoDSF) Prometheus NT 48 device (NanoTemper Technologies). The temperature range for measurement was from 20°C to 95°C with a heating rate of 1°C min^−1^. Changes in the ratio of the fluorescence intensities at 350 nm and 330 nm were monitored. The indicated melting temperature (mean of two technical replicates) corresponds to the point of inflection (maximum of the first derivative). The melting temperatures of the proteins are listed in **Supplementary Table 1**.

### Kinprola labeling kinetics

The labeling kinetics of Kinprola was measured by recording fluorescence polarization over time at 30 °C using a Tecan microplate reader. The measurements were conducted in black, non-binding, flat bottom, 96-well polystyrene plates (OptiPlate, PerkinElmer). Experiments were performed in kinase assay buffer (see **Supplementary table 4**) with or without PKAcat and ATP, in technical triplicates. In a 200 μL assay system, final concentrations were as follows: 200 nM Kinprola protein, 25 ng μL^-1^ PKAcat, 500 μM ATP and 50 nM fluorescent HaloTag substrate (CPY-CA, TMR-CA, JF_549_-CA, JF_552_-CA, or JF_669_-CA). Kinprola protein and fluorescent substrate were prepared separately in 50 μL aliquots, and PKAcat + ATP was prepared in 100 μL. Kinprola protein was first incubated with kinase assay buffer containing PKAcat + ATP for 30 min at 30 °C. The reactions were then started by adding 50 μL of the fluorescent HaloTag substrate. Control experiments were conducted with the same procedure but in kinase assay buffer without PKAcat and ATP. Data analysis was performed as previously described^17^. Kinprola labeling kinetics parameters are listed in **Supplementary Table 2**.

### Cell culture

HeLa Kyoto Flp-In (provided by Dr. Amparo Andres-Pons, EMBL, Heidelberg), human embryonic kidney 293 (HEK293) Flp-In T-REx cells, and HEK293T cells were cultured in Dubeco’s Modified Eagle Medium (DMEM) supplemented with 10% (vol/vol) fetal bovine serum (FBS) in a humidified 5% CO_2_ incubator at 37°C. RKO cells were cultured in RPMI + GlutaMAX medium supplemented with 10% (vol/vol) FBS,100 units mL^-1^ penicillin, and 100 µg mL^-1^ streptomycin. Primary rat hippocampal neurons were cultured in NeuroBasal medium supplemented with 1× GlutaMAX, 1× B-27, 100 units mL^-1^ penicillin, and 100 µg mL^-1^ streptomycin. The patient-derived glioblastoma cell line (PDGCL) S24^31,51^ was cultured as non-adherent neurospheres in PDGCL medium, consisting of DMEM/F-12, 1× B-27 supplement, 5 μg mL^−1^ insulin, 5 μg mL^-1^ heparin, 20 ng mL^-1^ epidermal growth factor, and 20 ng mL^-1^ basic fibroblast growth factor. Cell lines were regularly tested for mycoplasma contamination and were mycoplasma-free.

### Transient transfection of cells

Transient transfection was performed using Lipofectamine 3000 transfection reagent unless otherwise specified. For transfecting a single well of a 96-well plate, 100 ng of DNA was first diluted into 10 μL of OptiMEM I, and mixed with 0.2 μL of P3000. Separately, 0.2 μL of Lipofectamine 3000 was diluted with 10 μL of OptiMEM I. The two solutions were then mixed and incubated for 15 min at room temperature. The prepared DNA–Lipofectamine complex was added to cells at 50–70% confluency. The medium was changed after 12 h of incubation in a humidified 5% CO_2_ incubator at 37°C. The cells were then cultured under the same conditions for another 12–36 h before further treatment.

### Stable cell line establishment

HeLa and HEK293 stable cell lines were generated using the Flp-In system. Briefly, HEK293 Flp-In T-Rex or HeLa Kyoto Flp-In cells were cultured to 80% confluency in T-25 cell culture flasks. Cells were then co-transfected with 440 ng of a pCDNA5/FRT plasmid encoding the gene-of-interest (GOI) and 3560 ng of the pOG44 Flp-recombinase expression plasmid (Invitrogen) using Lipofectamine 3000 transfection reagent as described above. The cell culture medium was changed 12 h post-transfection, and the medium was replaced with cell culture medium supplemented with 100 μg mL^-1^ hygromycin B 24 h post-transfection to select cells that stably integrated the GOI into the genome. After 48–72 h of selection, cells were recovered in fresh cell culture medium until reaching confluency. Cells with high expression levels of GOI were sorted in bulk population based on EGFP fluorescence intensity (blue laser, 488 nm with 530/30 bandpass filter) using fluorescence-activated cell sorting (FACS) with a FACSMelody cell sorter (BD Biosciences). A list of established stable cell lines can be found in **Supplementary Table 5**.

### Recombinant adeno-associated viruses (rAAVs) preparation

rAAVs used for cell experiments were produced as previously described^52^. Briefly, HEK293 cells were transfected with plasmids pRV1 (containing AAV2 Rep and Cap sequences), pH21 (containing AAV1 Rep and Cap sequences), pFD6 (adenovirus helper plasmid) and the AAV plasmid containing the recombinant expression cassette driven by the hSyn1 or CAG promoter and flanked by AAV2 packaging signals (ITRs). Transfection was performed using polyethylenimine (PEI) 25,000. Five days post-transfection, the culture medium and cells were harvested by centrifugation at 1,000 g for 5 min at 4°C. The cells were lysed using TNT extraction buffer (see **Supplementary table 4**). Cell debris was removed by centrifugation at 3,000 g for 5 min at 4°C. The supernatant was treated with 50 U mL^−1^ benzonase nuclease for 30–60 min at 37°C, with mixing by inverting every 20 min. rAAVs were purified from the medium and supernatant via Äkta-Quick FPLC system (Cytiva) with AVB Sepharose HiTrap columns (Cytiva). The columns were first equilibrated with PBS (pH 7.4), and the virus particles were then eluted with 50 mM glycine-HCl (pH 2.7). Purified virus particles were concentrated and buffer exchanged to PBS (pH 7.3) using Amicon Ultra centrifugal filters (Millipore) with a molecular weight cutoff of 100 kDa. The rAAVs were aliquoted, flash-frozen, and stored at −80°C until further use. The rAAV titer was quantified by quantitative PCR (qPCR) as previously described^53^. For mice brain injection and expression, AAV9-hSyn-Kinprola_PKA_ and AAV9-hSyn-Kinprola_PKA_T/A_ were packaged at BrainVTA.

### Lentivirus production

Lentivirus were produced as previously described^54^. Briefly, the lentiviral packaging vector psPAX2 and the lentiviral envelope vector pMD2.G were co-transfected with the respective lentiviral expression vector (in a ratio of 5.25:3.15:10) using TransIT-LT1 transfection reagent and Opti-MEM, according to the manufacturer’s protocol, into low-passage (< 15) HEK293T cells at 70–80% confluency. Approximately 16 h post-transfection, the medium was replaced with fresh culture medium and supernatant was harvested 48 h post-transfection by filtration through a 0.45 μm low protein binding PES membrane. The harvested lentiviral supernatant was aliquoted and stored at −80°C. For determining the multiplicity of infection (MOI), RKO cells were transduced with varying amounts of lentiviral supernatant in the presence of 10 μg mL^-1^ polybrene according to the manufacturer’s protocol. 24 h post-transduction, cells were selected with 2 μg mL^-1^ puromycin for another 48 h, and the number of surviving cells was compared to a non-transduced control sample. The titer of HD CRISPR sub-library A in RKO cells was determined to be 2.19×10^7^ transduction units (TU) mL^-1^.

### Cell fixation

After treatment and labeling, cells were washed with pre-warmed PBS and fixed with 4% (wt/vol) methanol-free formaldehyde (PFA) in PBS at 37°C for 15 min. Subsequently, the cells were washed three times with PBS for further imaging.

### Recording PKA activities in cells during pharmacological treatment

In general, cells expressing Kinprola_PKA_ variants were seeded into chambered coverslips or 96-well culture dishes and grown to approximately 80% confluency. Cells were treated with various compounds (50 μM forskolin (Fsk)/100 μM IBMX, 1 μM prostaglandin E1 (PGE1), 10 μM isoproterenol (Iso), 1 μM epinephrine (Epi), 1 mM Bt_2_cAMP, 20 μM H89, 100 nM thapsigargin, 100 ng mL^-1^ phorbol 12-myristate 13-acetate (PMA), 10 μM anisomycin, 1 μM ionomycin, 40 mM 2-deoxy-D-glucose (2-DG)) in the presence of 25 nM CPY-CA for 30 min in a humidified incubator at 37°C with 5% CO_2_ atmosphere. After treatment and labeling, cells were then washed with PBS and incubated with medium supplemented with 5 μM recombinant HaloTag protein for 10 min. Subsequently, cells were washed again with PBS and either fixed for fluorescence imaging or detached with transparent TrypLE Express Enzyme for flow cytometry analysis.

### Real-time recording of labeling signal integration in HEK293 cells

HEK293 cells stably expressing Kinprola_PKA_ were seeded into poly-D-lysine-coated 8-well imaging chambered coverslips and cultured in 150 μL of transparent medium per well. During time-lapse imaging, five baseline frames were first captured. Then, the medium was replaced with 150 μL of medium containing 25 nM CPY-CA, followed by a 10-min imaging acquisition period. Afterwards, 150 μL of medium containing 25 nM CPY-CA and 100 μM Fsk was added, and another 5-min imaging acquisition was performed. Finally, 150 μL of medium containing 25 nM CPY-CA, 50 μM Fsk, and 60 μM H89 was added for approximately 15-min of imaging acquisition. The final concentrations of CPY-CA, Fsk and H89 in the imaging medium were maintained constant at 25 nM, 50 μM and 20 μM, respectively. Signal background from free CPY-CA was subtracted using co-cultured cells without Kinprola_PKA_ expression.

### Successive recordings during pharmacological treatment in HEK293 cells

HEK293 cells stably expressing Kinprola_PKA_ were seeded into poly-D-lysine-coated 96-well imaging plates. For successive recordings with spectrally distinguishable fluorescent HaloTag substrates, cells were sequentially treated as follows: (1) co-treated with 10 μM PGE1 and fluorophore substrate 1 for 30 min; (2) co-treated with 10 μM H89 and fluorophore substrate 2 for 30 min; (3) co-treated with 50 μM Fsk/100 μM IBMX and fluorophore substrate 3 for 20 min. Between different recordings, cells were washed with medium supplemented with 5 μM recombinant HaloTag protein for 10 min, followed by two washes with medium, and then allowed to rest for 2 h in fresh medium in a humidified incubator at 37°C with 5% CO_2_ atmosphere. For these recordings, the following substrates were used: 25 nM CPY-CA, 100 nM JF_552_-CA and 100 nM JF_669_-CA. After the final recording session, cells were fixed for further imaging as described above.

### Recording activities of different kinases in cells during pharmacological treatment

HeLa or HEK293 cells were seeded into 96-well culture dishes and grown to approximately 50% confluency. For HEK293 cells, the 96-well culture dishes were pre-coated with poly-D-lysine. Kinprola variants plasmids were then transfected into cells using Lipofectamine 3000. 24 h post-transfection, cells were treated with various compounds (for Kinprola_PKC_ and Kinprola_PKC_T/A_, 100 ng mL^-1^ PMA, 1 μM Gö 6983; for Kinprola_JNK_ and Kinprola_JNK_T/A_, 10 μM anisomycin, 10 μM JNK inhibitor III; for Kinprola_AMPK_ and Kinprola_AMPK_T/A_, 40 mM 2-DG, 30 μM SBI-0206965 (SBI) in the presence of 25 nM CPY-CA for 30 min in a humidified incubator at 37°C with 5% CO_2_ atmosphere. After treatment, cells were then washed with PBS and incubated with medium supplemented with 5 μM recombinant HaloTag protein for 10 min. Subsequently, cells were washed again with PBS and detached with transparent TrypLE Express Enzyme for flow cytometry analysis.

### Simultaneously recording PKA activities in different cellular compartments during pharmacological treatment

HeLa cells were transduced using AAVs under a CAG promoter with NES-Kinprola_PKA_-mEGFP and Kinprola_PKA_-mTagBFP2-NLS 3×. 36 h post-transduction, cells were stimulated with 50 μM Fsk/100 μM IBMX in the presence of 50 nM CPY-CA for 30 min in a humidified incubator at 37°C with 5% CO_2_ atmosphere. After treatment, cells were then washed with PBS and incubated with medium supplemented with 5 μM recombinant HaloTag protein for 10 min. Subsequently, cells were washed again with PBS and fixed for fluorescence imaging.

### Generation of glioblastoma cells stably expressing Kinprola

The Kinprola expression cassette was cloned into a pLKO.1-puro vector for lentivirus production. Lentivirus transduction of glioblastoma S24 cells was performed as previously described^28^. Successfully transduced S24 cells were selected using 1 μg mL^-1^ puromycin. EGFP-positive cells were subsequently sorted using a FACSMelody cell sorter (BD Biosciences, excitation 488 nm, filter 530/30 nm) and propagated for further use.

### RNA-Seq data generation

RNA-Seq sample preparation was conducted as previously described with modifications^17^. Glioblastoma S24 cells stably expressing Kinprola_PKA_ or Kinprola_on_ were plated at a density of 4×10^6^ cells onto Matrigel-coated T-25 cell culture flasks in growth factor-devoid PDGCL medium supplemented with 50 mM glucose (high-glucose medium, HGM). These serum-free stem-like conditions preserve both the gene expression and biological properties of the original tumor such as diffuse growth and network formation^55^. 48 h after seeding, Kinprola_PKA_-expressing S24 cells were labeled with 100 nM CPY-CA for 30 min at 37°C in a humidified incubator with 5% CO_2_ atmosphere, while Kinprola_on_-expressing S24 cells were labeled for a decreased time of 2.5 min as a control. Afterwards, cells were rinsed with HGM and incubated with HGM containing 4 μM recombinant HaloTag protein for 10 min. The cells were then rinsed with PBS, detached using accutase, resuspended in cold PBS, and subjected to FACS sorting on a FACSAria Fusion Special Order System (BD Biosystems). Kinprola_PKA_ or Kinprola_on_-expressing S24 cells were sorted into three groups based on high (around 5%), medium, and low (around 5%) normalized fluorescence intensity (CPY-CA/EGFP). Three replicate samples per group were collected, and RNA was isolated with the Arcturus PicoPure Frozen RNA Isolation Kit, according to the manufacturer’s instructions. On-column DNase digestion was performed using the RNase Free DNase Set and RNA integrity was verified using the high-sensitivity RNA ScreenTape System (Agilent) and the 4150 Tapestation System (Agilent). Library preparation and RNA sequencing of the three replicates per condition were performed on a NovaSeq6000 device (Illumina) by the Genomics and Proteomics Core Facility at the German Cancer Research Center (DKFZ, Heidelberg).

### RNA-Seq data analysis

RNA-Seq reads were aligned with STAR^56^ (v.2.5.3a) against the GRCh38 human reference genome. A gene-count matrix was generated using featureCounts^57^ in Subread (v.1.5.3) against GENCODE^58^ (v.32). Pairwise differentially expressed genes (DEGs) between groups were identified with generalized linear models using DeSeq2^59^ (v1.38.3) (i.e., DEGs in “high” *vs.* “medium”, “high” *vs.* “low” and “medium” *vs.* “low”). DEGs were retained with a false discovery rate (FDR) < 0.05, a raw count > 9 in all samples and replicates, a log_2_ fold change (log_2_FC) > 0.5 and < −0.5, and those recurrently identified in all three comparisons. The analysis yielded 737 DEGs in Kinprola_PKA_ data and 326 DEGs in Kinprola_on_ data. Expression levels (log_2_ fragments per kilobase per million mapped fragments) of DEGs were z-score scaled across samples and visualized with GraphPad Prism (version 10.2.1). Multidimensional scaling plots of RNA-Seq datasets with the first principal coordinate (PCoA1) were plotted using the plotPCA function after normalization with the vst() function in DESeq2^59^. ShinyGO^60^ (RRID:SCR_019213, v.0.741 and v.0.80) was used for gene ontology (GO) enrichment analysis. FDR and fold enrichment were calculated by comparing DEGs lists with a background of all protein-coding genes in the human genome.

### Generation of RKO cells stably expressing Kinprola and Cas9

The Kinprola expression cassette was cloned into a lenti-EF1α-Cas9-T2A-Blasticidin expression plasmid (see **Supplementary note 1**). Lentivirus production was performed as described above. Successfully transduced RKO cells were selected using 15 μg mL^−1^ Blasticidin for 7 days. EGFP-positive cells were sorted using a FACSMelody cell sorter (BD Biosciences, excitation 488 nm, filter 530/30 nm) and propagated for further use.

### Recording PKA activities during pharmacological treatment in RKO cells

RKO cells stably expressing Kinprola_PKA_, Kinprola_PKA_T/A_, and Kinprola_on_ were seeded at 8×10^3^ cells per well into 96-well plates. Two days post-seeding, cells were incubated with vehicle or with compounds (50 μM Fsk/100 μM IBMX or 20 μM H89) in the presence of 125 nM CPY-CA for 30 min at 37°C in a humidified incubator with 5% CO_2_ atmosphere. Cells were then rinsed with fresh medium, incubated with medium containing 5 μM recombinant HaloTag protein for 10 min, rinsed with PBS, detached using transparent TrypLE Express Enzyme, resuspended into PBS containing 2% (vol/vol) FBS, and finally analyzed via flow cytometry.

### Lentiviral CRISPR sgRNA library preparation

The HD CRISPR library sub-library A was constructed on an HD CRISPR vector as previously described^36^. Briefly, lentivirus was produced using HEK293T cells as the host. Together with psPAX2 and pMD2.G packaging plasmids, the sgRNA plasmid library was transfected into HEK293T cells using TransIT-LT1 transfection reagent. 48 h post-transfection, virus-containing supernatant was filtered through a 0.45 μm low protein binding PES membrane, aliquoted and stored at −80°C until further use.

### Pooled CRISPR knockout screen

A total of 2.5×10^8^ RKO cells stably expressing Kinprola_PKA_ and Cas9 were transduced with the lentiviral HD CRISPR sub-library A and 10 μg mL^−1^ polybrene in 5-layer cell culture multi-flasks to achieve an initial library coverage of around 500-fold upon infection at an MOI of 0.2 to 0.3 (ensuring the large majority of cells receive only one sgRNA for gene editing). 24 h post-transduction, the transduced cells were selected with 2 μg mL^−1^ puromycin. One flask was cultured without puromycin selection for MOI determination. Then, 48 h post-selection, cells were passaged and the MOI was determined by calculating the ratio of live cells with and without puromycin selection. Cells were re-seeded at a 500-fold library coverage and cultured for another two days. To prepare labeled cells for sorting, cells were incubated with 125 nM CPY-CA for 30 min at 37°C in a humidified incubator with 5% CO_2_ atmosphere. Cells were further rinsed with PBS, and washed with medium containing 2 μM recombinant HaloTag protein for 15 min. Then the cells were detached with accutase, pelleted by centrifugation at 300 g for 5 min, resuspended in ice-cold PBS supplemented with 2% (vol/vol) FBS and 5 mM EDTA, filtered with cell strainers, and stored on ice until sorting. The labeled cells were sorted into three groups with the high (around 25%), medium (around 50%) and low (around 25%) normalized fluorescence intensity (CPY-CA/EGFP). Once sorted, the cells were pelleted by centrifugation at 300 g for 5 min, washed with PBS, and stored dry at −20°C until genomic DNA extraction.

### Genomic DNA extraction, library preparation, and sequencing

These steps were performed as previous described with modifications^36^. Briefly, genomic DNA was isolated from frozen cell pellets using the QIAamp DNA Blood & Cell Culture DNA Maxi Kit according to the manufacturer’s instruction and purified by ethanol precipitation. DNA concentration in all subsequent steps was measured using Nanodrop or Qubit dsDNA HS and BR Assay Kits. For PCR amplification of the sgRNA cassette, 100 μg of genomic DNA was used with unique indexing primers for each sample. Illumina adapters and indices were added in the same one-step PCR reaction using the KAPA HiFi HotStart ReadyMix. The PCR product was purified using the QIAQuick PCR Purification Kit and further gel purified to remove genomic DNA contamination using the QiaQuick Gel Extraction kit. Quality and purity of the amplified PCR product were determined using the Agilent 2100 Bioanalyzer system. DNA concentrations from all conditions were adjusted and pooled at equimolar ratios. The sequencing reaction was performed on an Illumina NextSeq 500/550 system with a High-Output Kit v2.5 (75 cycles) to read the 20-nt sgRNA sequence and quantify the number of copies, by the Deep Sequencing Core Facility (Bioquant, Heidelberg University).

### Pooled CRISPR knock screen data processing and analysis

Absolute sgRNA read counts were collected and demultiplexed using the MAGeCK version 0.5.9.4 package^61^. To process the raw data, read counts were first normalized and log-transformed. Fold changes between conditions were then determined by subtracting the log-normalized read count of the control samples from that of the corresponding treated sample. Replicates were collapsed by arithmetic mean for each gene and each sgRNA. Statistical significance of differential gene expression was calculated using the LIMMA package^62^. *p* values were corrected for multiple testing by Benjamini−Hochberg correction. An FDR cut-off of 0.05 was applied to hit selection. The hit list was further filtered by eliminating RKO core-essential genes according to dependency score available in the Cancer Dependency Map Project (DepMap, https://depmap.org/).

### Hits validation by individual retests of select sgRNAs

Two non-targeting (NT) sgRNAs and two sgRNAs targeting each hit of interest were cloned into the parental HDCRIPSRv1^36^ vector for sgRNA expression under a U6 promoter (see **Supplementary table 6**). RKO cells stably expressing Kinprola_PKA_ and Cas9 were seeded into 6-well plates at a density of 5×10^5^ cells per well. After 24 h, the cells were transfected with plasmids containing the sgRNAs with jetOPTIMUS transfection reagent. For a single well transfection, 3 μg of plasmid and 3 μL of jetOPTIMUS transfection reagent in 200 μL of jetOPTIMUS buffer were used. The cells were changed into new culture medium after 6 h. 24 h after transfection, the cells were selected with new culture medium supplemented with 1.25 μg mL^−1^ puromycin. After 48 h of selection, the cells were labeled with 125 nM CPY-CA for 30 min, and prepared as described above for flow cytometer analysis. SYTOX blue dead cell stain was used for gating out dead cells according to manufacturer’s protocol.

### ELISA-based colorimetric PKA activity measurement

PKA activity in cell lysate was measured using an ELISA-based PKA colorimetric activity kit according to the manufacturer’s instructions. Briefly, sgRNA-transfected cells, prepared as described above, were collected using a cell scraper, and lysed in activated cell lysis buffer (see **Supplementary table 4**) for 30 min on ice with occasional vortexing, followed by centrifugation at 10,000 rpm for 10 min at 4°C. The supernatants were then frozen as single-use aliquots at −80°C. Protein concentrations were quantified using a Bicinchoninic acid assay (BCA) before the following colorimetric assay. The colorimetric assay was performed according to the manufacturer’s protocol, and the optical density was measured using a Tecan microplate reader.

### Primary rat hippocampal neurons preparation

All procedures were conducted in strict accordance with the Animal Welfare Act of the Federal Republic of Germany (Tierschutzgesetz der Bundesrepublik Deutschland, TierSchG) and the Animal Welfare Laboratory Animal Regulations (Tierschutzversuchsverordnung). According to these regulations, no ethical approval from an ethics committee is required for euthanizing rodents when the organs or tissues are used for scientific purposes. The euthanasia procedure for rats in this study was supervised by animal welfare officers of the Max Planck Institute for Medical Research and was carried out and documented in compliance with the TierSchG (permit number assigned by the Max Planck Institute for Medical Research: MPI/T-35/18). Primary rat hippocampal neurons were prepared from isolated hippocampi obtained from postnatal P0– P1 Wistar rats of both sexes, following the established protocols as previously described^63^. Neurons were seeded onto poly-L-ornithine (100 μg mL^−1^ in water) and laminin (1 μg mL^−1^ in HBSS) 24-well or 96-well glass bottom imaging plates, and maintained in a humidified cell culture incubator with 5% CO_2_ at 37°C.

### rAAV transduction

At day 6, neurons were refreshed with one-third of the medium. On day 7, neurons were transduced with purified rAAVs (serotype 2/1) at concentrations ranging from 10^9^ to 10^10^ genome copies mL^−1^. Cultures were allowed to express transgenes for 7 days. One-third of the medium was changed every three days during this period.

### Recording PKA activities in cultured neurons during pharmacological treatment

Neurons seeded in 96-well glass bottom imaging plates were used at 14–16 days in vitro (DIV). PKA modulators (50 μM Fsk/2 μM Rolipram (Rol), 1 μM Iso, 1 μM propranolol, 1 μM norepinephrine, or 20 μM H89) were applied along with 25 nM CPY-CA to neuronal cultures. The treatments were conducted in a humidified cell culture incubator with 5% CO_2_ at 37°C for 45 min. Afterwards, neurons were rinsed with warm NeuroBasal medium, followed by incubation with warm NeuroBasal medium supplemented with 5 μM recombinant HaloTag protein for 15 min. Neurons were then washed with warm HBSS and fixed with 4% (wt/vol) PFA for 15 min at 37°C. After fixation, neurons were washed with HBSS and stored at 4°C for further imaging.

### Recording PKA activities in cultured neurons during electric field stimulation

Cultured neurons in 24-well glass bottom imaging plates were prepared for electric field stimulation using a custom-build 24-well cap stimulator equipped with platinum electrodes connected to a stimulation control unit, as previously described^64^. Prior to stimulation, neurons were treated with a synaptic blocker solution (25 μM APV/10 μM NBQX in NeuroBasal medium) for 25 min at 37 °C in a humidified cell culture incubator with 5% CO_2_. The neuron cultures were then transferred to a widefield microscope stage housed in an environmental chamber set to 37°C with 5% CO_2_. The cap stimulator was positioned on top of the neuron cultures. Neurons were pre-incubated with 125 nM CPY-CA for 5 min prior to stimulation. The stimulation patterns were set to 80 Hz frequency, 100 mA intensity and 1 ms pulse width, generating defined trains of action potentials in the presence of CPY-CA. Following electric field stimulation, neurons were washed and fixed for further imaging, following the procedures described above.

### Stability measurement of fluorescent Kinprola_PKA_ labeling signal in cultured neurons

Neurons were incubated with 125 nM CPY-CA for 1 h at 37°C in a humidified cell culture incubator with 5% CO_2_, and then washed as described above. Experiments were conducted in three 24-well imaging plates as replicates. The fluorescent intensities of basally active neurons were recorded and measured every 24h for three days. Between measurements, neurons were maintained at 37°C in a humidified cell culture incubator with 5% CO_2_.

### Animals

All procedures for animal surgery and experimentation were performed using protocols approved by the Institutional Animal Care and Use Committee at Peking University. Mice were group- or pair-housed in a temperature-controlled (18–23°C) and humidity-controlled (40–60%) room with a 12-h light/dark cycle. Food and water were available ad libitum.

### Expression of Kinprola in mice brain

Male C57BL/6N mice (6–8 weeks of age) were anesthetized with an intraperitoneal injection of 2,2,2-tribromoethanol (Avertin; 500 mg kg^−1^ of body weight). The AAV9-hSyn-Kinprola_PKA_ (500 nL, 2.5×10^12^ viral genomes (vg) mL^-1^, BrainVTA) and AAV9-hSyn-Kinprola_PKA_T/A_ (300 nL, 6×10^12^ vg mL^-1^, BrainVTA) viruses were injected into the bilateral nucleus accumbens (NAc) separately (AP: +1.4 mm relative to Bregma; ML: ±1.2 mm relative to Bregma; DV: −4.0 mm from the dura) at a rate of 50 nL min^−1^. The experiments were performed 2−3 weeks after virus injection.

### Acute mouse brain slices preparation

the mice were anesthetized with 2,2,2-tribromoethanol (Avertin, 500 mg kg^−1^ body weight) and perfused with ice-cold oxygenated slicing buffer (see **Supplementary table 4**). The brains were then dissected, and coronal slices of 300 μm thickness were obtained using a VT1200 vibratome (Leica) in ice-cold oxygenated slicing buffer. These slices were then transferred in oxygenated artificial cerebrospinal fluid (ACSF, see **Supplementary table 4**) and allowed to recover for at least 30 min at 34°C.

### Recording PKA activities in acute mouse brain slices during pharmacological treatment

The acute brain slices were moved to a custom-made perfusion chamber and placed on the stage of an upright LSM 710 confocal microscope (ZEISS). During time lapse imaging experiments, the slices were perfused with oxygenated ACSF containing 250 nM CPY-CA with or without 50 μM Fsk/2 μM Rol. After imaging, the slices were transferred to oxygenated ACSF containing 2 μM recombinant HaloTag protein for 10 min. Subsequently, the brain slices were fixed with 4% (wt/vol) PFA at 4°C overnight, followed by washing with 0.2% (vol/vol) Tween 20 in PBS for 2 h. For further imaging on a SP8X confocal microscope (Leica), the fixed brain slices were mounted using Fluoromount-G (SouthernBiotech) in a custom-made imaging chamber. For chamber preparation, a coverslip (no. 1.5, ∼0.17 mm thick, Paul Marienfeld) were cut to the desired size to serve as a spacer. Two spacers were stacked (∼0.34 mm thick) and two pairs of these stacks were placed at either side of a coverslip to form an imaging chamber, which was then sealed with epoxy prior to imaging.

### Recording neuromodulation-induced PKA activation in mice brain during D1/D5R agonist SKF-81297 treatment

CPY-CA solution for *in vivo* administration were prepared as previously described^65^. Briefly, 100 nmol of CPY-CA was first dissolved in 20 μL DMSO. Then, 20 μL of a Pluronic F-127 solution (20% (wt/wt) in DMSO) was added and mixed by pipetting. This stock solution was diluted into 100 μL sterile saline. The dye solution was prepared freshly prior to injection to avoid freeze-thaw cycles. Mice were placed into the separated clean cages without food and water for 1 h of habituation over two consecutive days. Two weeks post-viral expression, mice were first injected with 100 nmol of CPY-CA solution via tail vein (intravenous, IV). After 10 min, the mice received intraperitoneal (IP) injections of SKF-81297 (10 mg kg^−1^, diluted in saline, 300 μL) or vehicle (equal volume of DMSO diluted in saline, 300 μL). Mice were placed in separated clean cages without food and water following the injection and were sacrificed 50 min after the IP injections. Mice were perfused with cold PBS supplemented with 50 μg mL^−1^ heparin, followed with cold 4% (wt/vol) PFA in PBS. Brains were dissected and fixed overnight at 4°C in 4% (wt/vol) PFA in PBS. The brains were then dehydrated with 30% (wt/vol) sucrose solution, embedded into OCT compound, and sectioned in the coronal plane at 40 μm thickness using a CM1900 cryostat (Leica). The brain slices were washed three times with 0.2% (vol/vol) Tween 20 in PBS, and mounted using Fluoromount-G for further imaging.

### Flow cytometry

Unless otherwise specified, the labeled cells were detached from culture plates using transparent TrypLE Express Enzyme and suspended in PBS containing 2% (vol/vol) FBS and transferred into U-shaped-bottom 96-well microplates. Cell samples were subjected to the autosampler of a BD LSRFortessa X-20 flow cytometry analyzer. Fluorescence recording parameters were set as follows: EGFP (Ex: 488 nm, Em: 530/30 nm), fluorophore excitation in 525–570 nm range (Ex: 561 nm excitation, Em: 586/15 nm) and fluorophore excitation in 620–680 nm range (Ex: 640 nm, Em: 670/30 nm). Photomultiplier tube detectors were adjusted to prevent signal saturation. The same recording parameters were used consistently throughout a set of experiment.

### Flow cytometry analysis

Raw data obtained from flow cytometry was imported into FlowJo suite (version 10.10.0) and processed as follows. First, live (SSC-A/FSC-A) and single cell (SSC-H/SSC-A) gates were gated and cells with EGFP fluorescence intensities below certain attribute unit (e.g., 10^3^ attribute unit) were excluded from further analysis to minimize background noise. Fluorescence intensity ratios were calculated for each cell by dividing the fluorescence intensities of certain fluorophores by the fluorescence intensities of EGFP. The same gating strategies were used consistently throughout a set of experiment. Quantitative assessment and statistical analysis were performed using either R package^66^ or GraphPad Prism (version 10.2.1).

### Microscopy

Fluorescence imaging for cultured cells and primary neurons was performed on a commercial Leica Stellaris 5 confocal microscope with a supercontinuum white light laser (470–670 nm) and hybrid photodetectors for single-molecule detection (HyD SMD). Laser power output was set to 85% of maximum power and regularly calibrated. The microscope stage was maintained in an environmental chamber (set to 37°C, 5% CO_2_). Before imaging, the imaging plate was equilibrated on the microscope stage for 30 min to avoid thermal drifting during image acquisition. Unless specified, the following imaging settings were used: a HC PL APO CS2 ×20/0.75-NA (numerical aperture) air/water objective, 581.82×581.82 μm, scan speed 400 MHz, Z-stacks with 2 μm step size. For time-lapse imaging, the settings were used: scan speed 600 MHz, 45 s per frame, Z-stacks with 2 μm step size and the physical length of 10 μm. Time-lapse imaging of acute mouse brain slices was performed on a LSM 710 upright microscope with a W N-Achroplan ×20/0.5-NA M27 water objective. The settings were used: 425.1×425.22 μm, pinhole 150 μm, zoom 1.0, pixel dwell time 2.55 μs, average 1; for EGFP channel, excitation 488 nm, emission 498–550 nm; for CPY-CA channel, excitation 633 nm, emission 638–747 nm. Fluorescence imaging of fixed brain slices was performed on a Leica SP8X laser scanning confocal system with a HC PL APO CS2 ×40/1.30-NA oil objective, a supercontinuum white light laser (470–670 nm) and HyD detectors. Unless specified, the settings were used: 387.5×387.5 μm, zoom 0.75, scan speed 400 MHz, Z-stacks with 2 μm step size and the physical length of 38 μm. Imaging acquisition parameters for each image were listed in **Supplementary Table 7**.

### Imaging processing and analysis

All images were processed and analyzed using ImageJ/Fiji (version 2.9.0/1.54h)^67^. Unless specified, Z-stack images (12 or 16 bit) were first converted into maximum intensity projections (MIP). From the channel indicating Kinprola expression (mostly EGFP channel), ROIs were delineated manually or segmented using Cellpose 2.0^68^ and mean fluorescence intensities from individual ROIs were derived for multiple fields-of-view. Unhealthy neurons were excluded from analyses. For tissue imaging, background signal was determined by average mean fluorescence intensities from 1–3 cell-free regions and subtracted for background correction. *BRET-Analyzer (v1.0.8)* plugin was employed for presenting ratiometric projections^69^. Specifically, the EGFP channel underwent thresholding (autoTh-Chastagnier method, Gaussian radius 1 and 2 using default value of 5 and 15), and fluorescence channels were divided (e.g., CPY-CA/EGFP) to generate ratiometric images.

### Data representation, reproducibility and statistical analysis

Numerical data was analyzed and plotted using the Excel (version 16.78.3), R package^66^, OriginPro 2020b (OriginLab) and GraphPad Prism (version 10.2.1). Schemes and figures were assembled in Adobe Illustrator 2024, using some elements adopted from BioRender (https://www.biorender.com/) through Max Planck Gesellschaft’s license. For fluorescence images presentation, brightness and contrast were adjusted identically for each channel. Unless specified, all in vitro measurements were performed in three technical replicates. All cell experiments were performed at least in three biological replicates. Statistical significance was determined by performing a two-tailed unpaired t-test with Welch’s correction, or one-way ANOVA with Dunnett’s or Tukey’s Post hoc test using the GraphPad Prism. The following notations apply for all statistical analyses: n.s. (not significant) *p* ≥ 0.05, * *p* < 0.05, ** *p* < 0.01, *** *p* < 0.001 and **** *p* < 0.0001. *p* values were provided for comparison.

