## Supplementary material for "Molecular recording of cellular protein kinase activity with chemical labeling": Sun et al_Kinprola_Supplemental Files

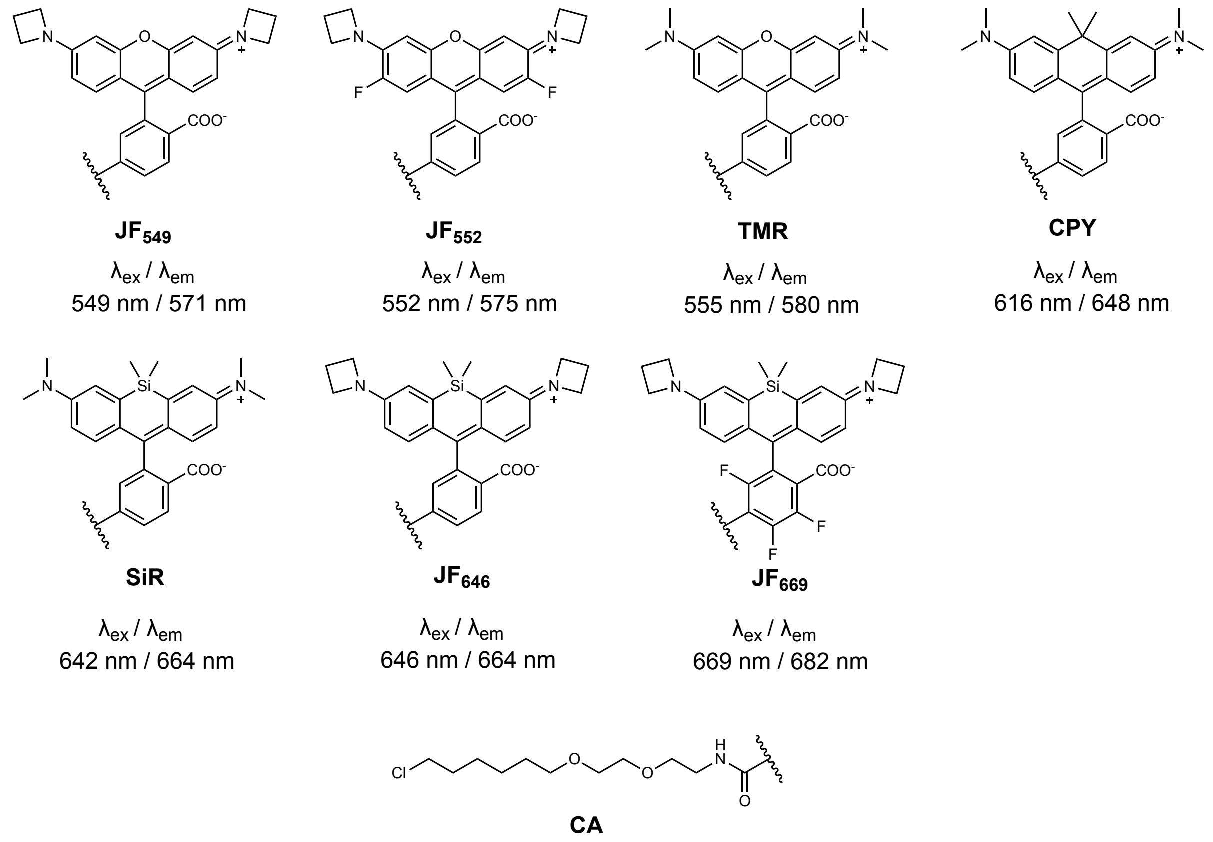


**Supplementary Fig. 1 | Fluorescent HaloTag substrates used in this study.** Chemical structures of fluorescent HaloTag substrates with excitation and emission maxima. Janelia Fluor (JF) fluorescent substrates were kind gifts of L. D. Lavis (Janelia Research Campus, Ashburn, Virginia). TMR-CA, CPY-CA and SiR-CA were synthesized in house according to literature procedures.

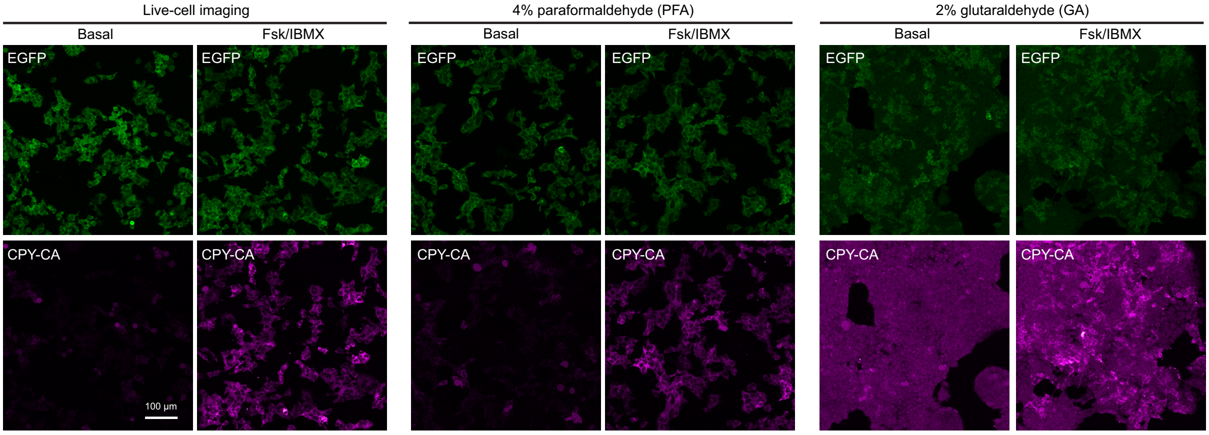


**Supplementary Fig. 2 | (related to Fig. 1) Kinprola-recorded fluorescent signal can be preserved by chemical fixation.** Fluorescence images of HEK293 cells stably expressing Kinprola$_{PKA}$ labeled with 25 nM CPY-CA for 30 min in the presence or absence of 50 µM Fsk/100 µM IBMX stimulation. The cells were then fixed with 4% paraformaldehyde (PFA) or 2% glutaraldehyde (GA) and subsequently imaged. Representative images from three wells of cell culture. Scale bar: 100 µm.

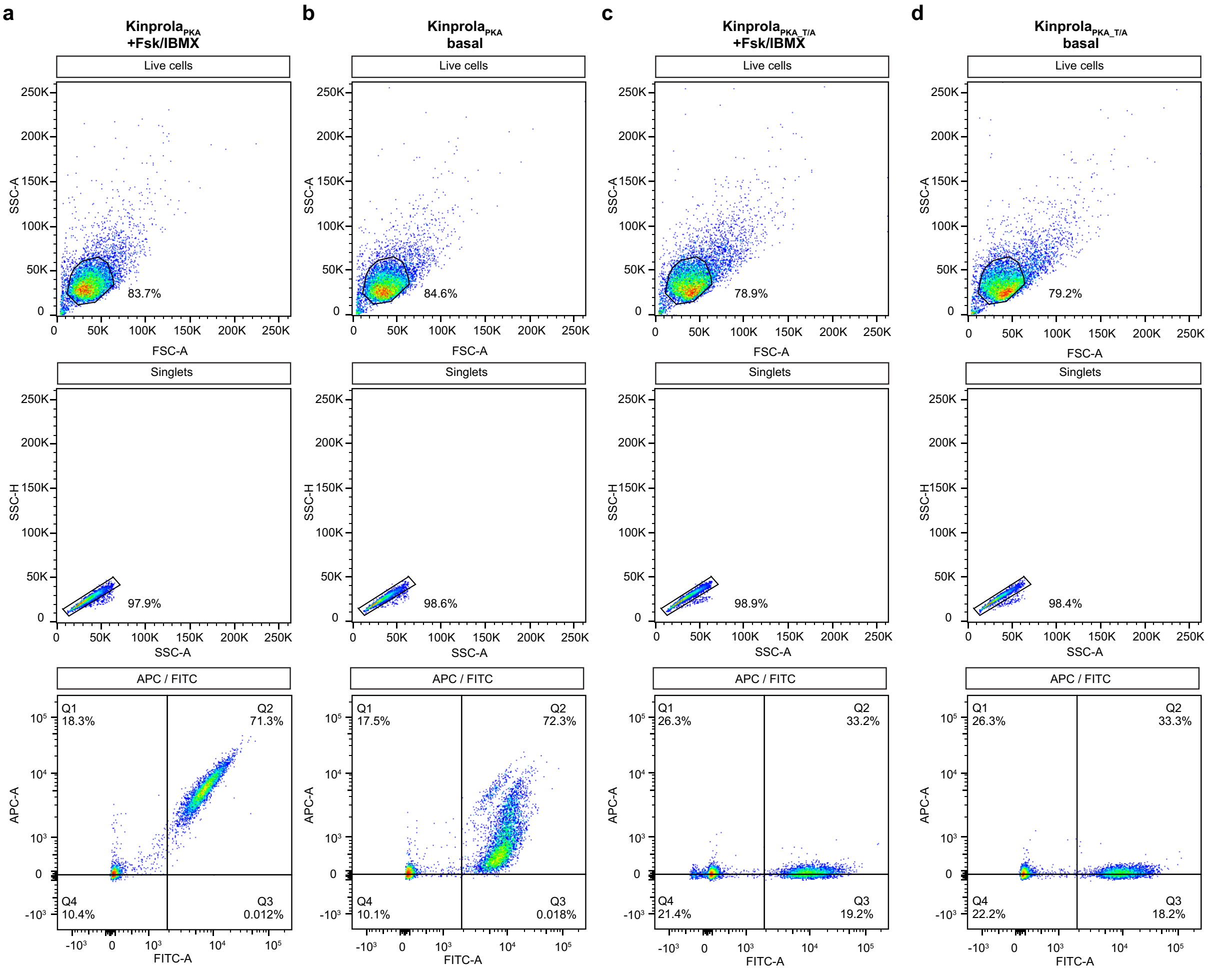


**Supplementary Fig. 3 | (related to Fig. 1) Representative flow cytometry gating strategy for HEK293 cells expressing Kinprola.** (**a**) Hierarchical gating of live cells (FSC-A *vs.* SSC-A), singlets (SSC-A *vs.* SSC-H) and positive cells (APC *vs.* FITC) for HEK293 cells expressing Kinprola$_{PKA}$ incubated with 25 nM CPY-CA for 30 min with 50 µM Fsk/100 µM IBMX stimulation. (**b**) Hierarchical gating for HEK293 cells expressing Kinprola$_{PKA}$ incubated with 25 nM CPY-CA for 30 min without stimulation (basal). (**c**) Hierarchical gating for HEK293 cells expressing Kinprola$_{PKA_T/A}$ incubated with 25 nM CPY-CA for 30 min with 50 µM Fsk/100 µM IBMX stimulation. (**d**) Hierarchical gating for HEK293 cells expressing Kinprola$_{PKA_T/A}$ incubated with 25 nM CPY-CA for 30 min without stimulation (basal).

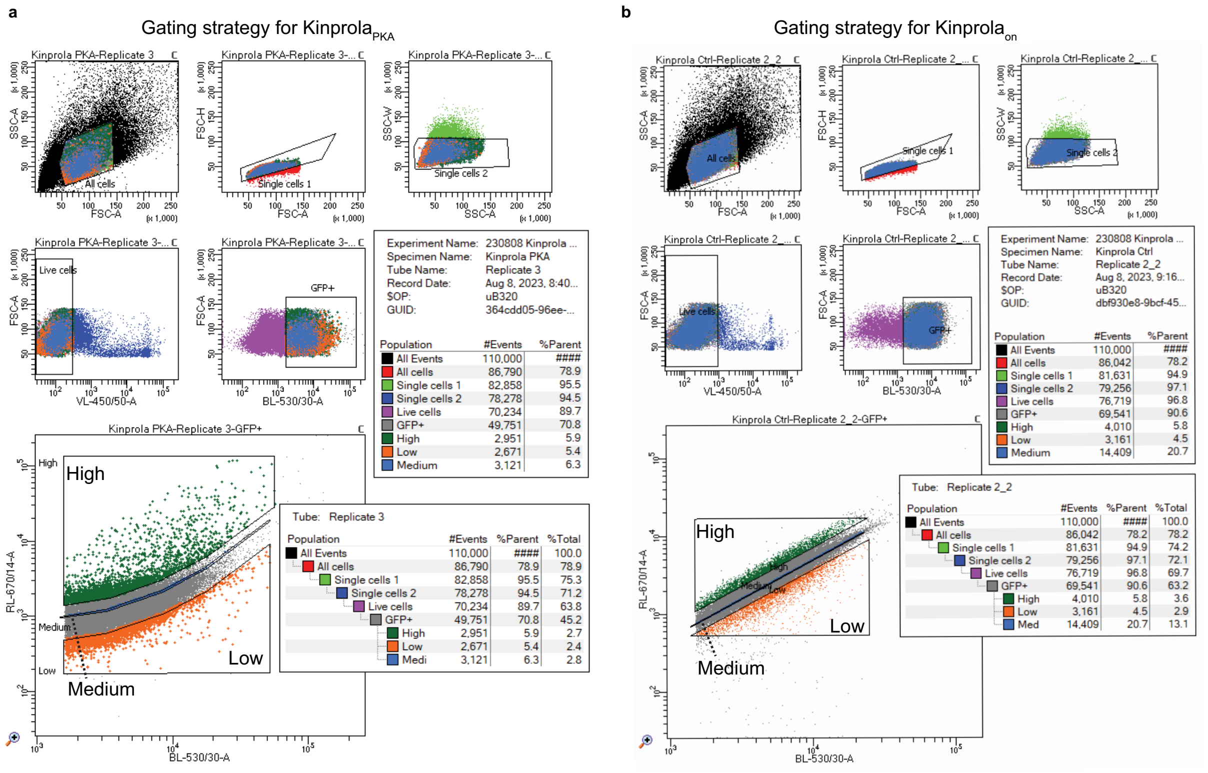


**Supplementary Fig. 4 | (related to Fig. 2) Gating and sorting strategies of Kinprola-expressing glioblastoma cells (GBCs) for RNA-Seq.** (**a**) Gating and sorting strategies of CPY-CA labeled Kinprola$_{PKA}$-expressing GBCs. (**b**) Gating and sorting strategies of CPY-CA labeled Kinprola$_{on}$-expressing GBCs.

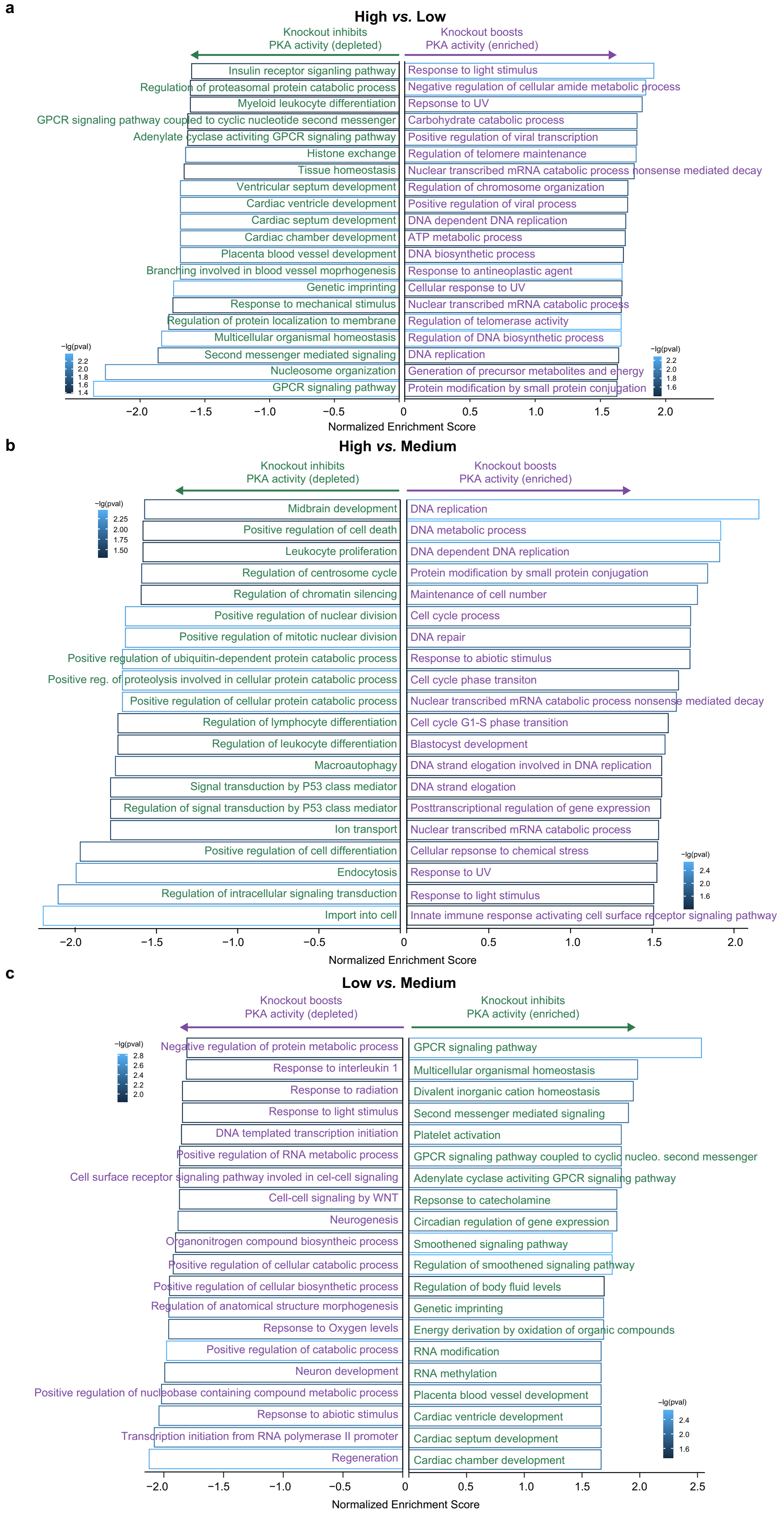
a
High vs. Low
Knockout inhibits PKA activity (depleted)
Knockout boosts PKA activity (enriched)
Insulin receptor siganling pathway
Regulation of proteasomal protein catabolic process
Myeloid leukocyte differentiation
GPCR signaling pathway coupled to cyclic nucleotide second messenger
Adenylate cyclase activiting GPCR signaling pathway
Histone exchange
Tissue homeostasis
Ventricular septum development
Cardiac ventricle development
Cardiac septum development
Cardiac chamber development
Placenta blood vessel development
Branching involved in blood vessel moprhogenesis
Genetic imprinting
Response to mechanical stimulus
Regulation of protein localization to membrane
Multicellular organismal homeostasis
Second messenger mediated signaling
Nucleosome organization
GPCR signaling pathway
Response to light stimulus
Negative regulation of cellular amide metabolic process
Repsonse to UV
Carbohydrate catabolic process
Positive regulation of viral transcription
Regulation of telomere maintenance
Nuclear transcribed mRNA catabolic process nonsense mediated decay
Regulation of chromosome organization
Positive regulation of viral process
DNA dependent DNA replication
ATP metabolic process
DNA biosynthetic process
Response to antineoplastic agent
Cellular response to UV
Nuclear transcribed mRNA catabolic process
Regulation of telomerase activity
Regulation of DNA biosynthetic process
DNA replication
Generation of precursor metabolites and energy
Protein modification by small protein conjugation
−lg(pval)
Normalized Enrichment Score
b
High vs. Medium
Knockout inhibits PKA activity (depleted)
Knockout boosts PKA activity (enriched)
Midbrain development
Positive regulation of cell death
Leukocyte proliferation
Regulation of centrosome cycle
Regulation of chromatin silencing
Positive regulation of nuclear division
Positive regulation of mitotic nuclear division
Positive regulation of ubiquitin-dependent protein catabolic process
Positive reg. of proteolysis involved in cellular protein catabolic process
Positive regulation of cellular protein catabolic process
Regulation of lymphocyte differentiation
Regulation of leukocyte differentiation
Macroautophagy
Signal transduction by P53 class mediator
Regulation of signal transduction by P53 class mediator
Ion transport
Positive regulation of cell differentiation
Endocytosis
Regulation of intracellular signaling transduction
Import into cell
DNA replication
DNA metabolic process
DNA dependent DNA replication
Protein modification by small protein conjugation
Maintenance of cell number
Cell cycle process
DNA repair
Response to abiotic stimulus
Cell cycle phase transiton
Nuclear transcribed mRNA catabolic process nonsense mediated decay
Cell cycle G1-S phase transition
Blastocyst development
DNA strand elogation involved in DNA replication
DNA strand elogation
Posttranscriptional regulation of gene expression
Nuclear transcribed mRNA catabolic process
Cellular repsonse to chemical stress
Response to UV
Response to light stimulus
Innate immune response activating cell surface receptor signaling pathway
−lg(pval)
Normalized Enrichment Score
c
Low vs. Medium
Knockout boosts PKA activity (depleted)
Knockout inhibits PKA activity (enriched)
Negative regulation of protein metabolic process
Response to interleukin 1
Response to radiation
Response to light stimulus
DNA templated transcription initiation
Positive regulation of RNA metabolic process
Cell surface receptor signaling pathway involed in cel-cell signaling
Cell-cell signaling by WNT
Neurogenesis
Organonitrogen compound biosyntheic process
Positive regulation of cellular catabolic process
Positive regulation of cellular biosynthetic process
Regulation of anatomical structure morphogenesis
Repsonse to Oxygen levels
Positive regulation of catabolic process
Neuron development
Positive regulation of nucleobase containing compound metabolic process
Repsonse to abiotic stimulus
Transcription initiation from RNA polymerase II promoter
Regeneration
GPCR signaling pathway
Multicellular organismal homeostasis
Divalent inorganic cation homeostasis
Second messenger mediated signaling
Platelet activation
GPCR signaling pathway coupled to cyclic nucleo. second messenger
Adenylate cyclase activiting GPCR signaling pathway
Repsonse to catecholamine
Circadian regulation of gene expression
Smoothened signaling pathway
Regulation of smoothened signaling pathway
Regulation of body fluid levels
Genetic imprinting
Energy derivation by oxidation of organic compounds
RNA modification
RNA methylation
Placenta blood vessel development
Cardiac ventricle development
Cardiac septum development
Cardiac chamber development
−lg(pval)
Normalized Enrichment Score

1

**Supplementary Fig. 5 | (related to Fig. 3) (a-c) Gene set enrichment analysis (GSEA) comparisons of cell subpopulations selected during CRISPR screening.** GSEA top 20 categories among "high" *vs.* "low", "high" *vs.* "medium" and "low" *vs.* "medium" comparisons. Bar graphs are color-coded by *p* values.

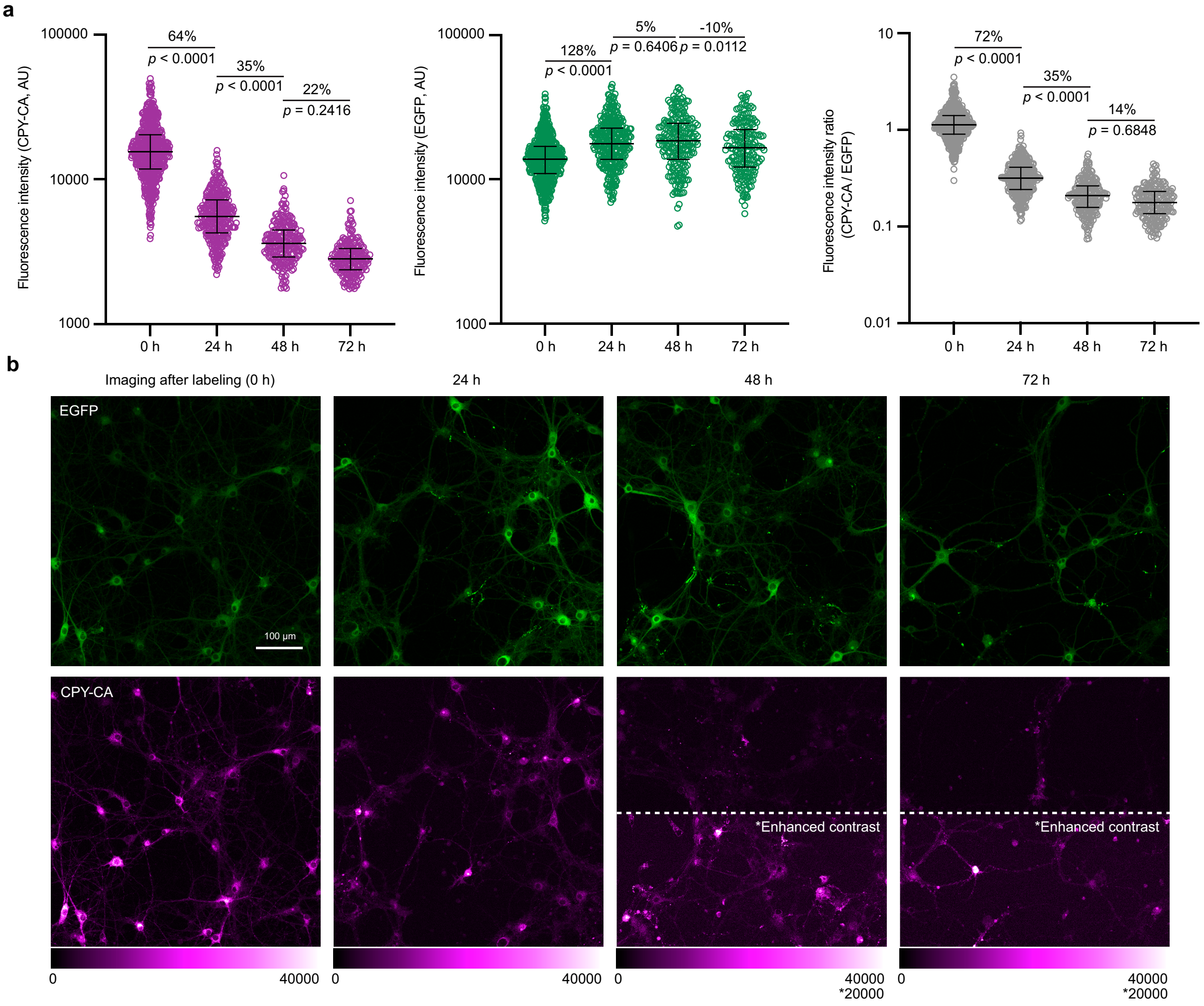


**Supplementary Fig. 6 | (related to Fig. 4) Stability of Kinprola$_{PKA}$ labeling signal in cultured primary hippocampal neurons over time.** (**a**) Dot plots showing the changes in fluorescence intensities (EGFP and CPY-CA) and fluorescence intensity ratios (CPY-CA/EGFP) of Kinprola$_{PKA}$-expressing primary hippocampal neurons over time. Neurons were labeled with 125 nM CPY-CA for 1 h without external stimulation. Images were acquired immediately after labeling (0 h) and subsequently every 24 h over a period of 3 days under live conditions. Error bars indicate median with interquartile range ($n \geq 205$ cells per group from three 24-well plates). Statistical significance was calculated with one-way ANOVA with Tukey's Post hoc test and $p$ values are given for comparison. (**b**) Representative images from the experiments described in (**a**). The contrast of images acquired at 48 h and 72 h was enhanced (CPY-CA channel, bottom half) to highlight that Kinprola$_{PKA}$ labeling can still be reliably detected after an extended period. Scale bar: 100 µm.

**Supplementary Table 1 | Protein melting temperature measured by NanoDSF in this study.**

| Purified protein | Average melting temperature (°C) |
|---|---|
| Kinprola$_{PKA}$ | 45.3 |
| Phosphorated Kinprola$_{PKA}$ | 49.8 |
| Kinprola$_{PKA_T/A}$ | 44.8 |
| Kinprola$_{on}$ | 45.8 |
| FHA1 | 52.3 |
| cpFHA1 | 37.2 |
| cpFHA1$^{N49Y}$ | 43.8 |

**Supplementary Table 2 | Biochemical characterization of Kinprola$_{PKA}$ *in vitro*.**

| **Fluorescent HaloTag substrates** | **Kinprola$_{PKA}$ $k_{PKAcat+ATP}$ ($M^{-1}s^{-1}$)** | **Kinprola$_{PKA}$ $k_{buffer}$ ($M^{-1}s^{-1}$)** | **$k_{PKAcat+ATP}$ / $k_{buffer}$** |
|---|---|---|---|
| CPY-CA | $2.56 \times 10^5$ ($2.54 - 2.58 \times 10^5$) | 54.68 (54.55 - 54.80) | 4682 (4656 - 4708) |
| TMR-CA | $1.37 \times 10^5$ ($1.36 - 1.38 \times 10^5$) | 63.48 (63.37 - 63.60) | 2158 (2146 - 2170) |
| $JF_{549}$-CA | $7.32 \times 10^4$ ($7.26 - 7.38 \times 10^4$) | 60.90 (60.79 - 61.00) | 1202 (1194 - 1210) |
| $JF_{552}$-CA | $5.46 \times 10^4$ ($5.44 - 5.49 \times 10^4$) | 57.02 (56.92 - 57.12) | 958 (956 - 961) |
| $JF_{669}$-CA | $3.84 \times 10^4$ ($3.81 - 3.87 \times 10^4$) | 136.74 (136.32 - 137.16) | 281 (279 - 282) |

**Supplementary Table 3 | Reagents and resource used in the study.**

| Reagents | Source | Identifier |
|---|---|---|
| *E.coli* strain 10G | Lucigen | 60108 |
| *E. coli* strain BL21(DE3) | Novagen | 69451 |
| NEB stable competent *E. coli* | NEB | C3040H |
| One Shot Stbl3 chemically competent *E. coli* | ThermoFisher Scientific | C737303 |
| Q5 high-fidelity DNA polymerase | NEB | M0491S |
| KOD-hot-start DNA polymerase master mix | Sigma-Aldrich | 71086 |
| Q5 site-directed mutagenesis kit | NEB | E0554S |
| In-Fusion Cloning | Takara Bio | 639650 |
| NucleoSpin Gel and PCR Clean-up, Mini kit | Macherey&Nagel | 740609.50 |
| QIAquick PCR Purification Kit | QIAGEN | 28106 |
| QIAprep Spin Miniprep Kit | QIAGEN | 27104 |
| GeneJET Endo-Free Plasmid Maxiprep Kit | ThermoFisher Scientific | K0861 |
| Quick Ligation Kit | NEB | M2200S |
| Ultra-15 Centrifugal Filter Unit | Amicon | UFC901024D |
| Quick CIP | NEB | M0525S |
| BsrGI-HF | NEB | R3575S |
| BfuAI | NEB | R0701S |
| Isopropyl-β-D-thiogalactopyranoside (IPTG) | Carl Roth | CN084 |
| Phenylmethylsulfonyl fluoride (PMSF) | ThermoFisher Scientific | 36978 |
| Lysozyme | ThermoFisher Scientific | 89833 |
| HisPur Ni-NTA Superflow Agarose | Thermo Scientific | 25217 |
| DMEM high glucose + GlutaMAX | Gibco | 31966021 |
| DMEM F12 | Gibco | 11330-032 |
| DMEM high glucose, phenol red-free | Gibco | 31053-028 |
| RPMI + GlutaMAX-I | Gibco | 61870-036 |
| Neurobasal medium | Gibco | 12348-017 |
| PBS pH 7.4 | Gibco | 10010-015 |
| TrypLE Express Enzyme | Gibco | 12604-013 |
| Accutase | Gibco | A1110501 |
| Opti-MEM | Gibco | 31985-054 |
| GlutaMAX | Gibco | 35050061 |
| B-27 | Gibco | 17504044 |
| B-27 Supplement, minus vitamin A | Gibco | 12587-010 |
| Insulin | Sigma-Aldrich | I9278 |
| Heparin | Sigma-Aldrich | H4784 |
| Human EGF Recombinant Protein | Gibco | PHG0311 |
| Human FGF-basic (FGF-2/bFGF) (aa 10-155) Recombinant Protein | Gibco | PHG0021 |
| Penicillin-Streptomycin (Pen/Strep) | Gibco | 15140122 |
| Blasticidin | InvivoGen | ant-bl-1 |
| Puromycin | biomol | Cay13884 |
| Hygromycin B | Carl Roth | 250-545-5 |
| Polybrene | Sigma-Aldrich | TR-1003-G |
| Poly-D-lysine | Sigma-Aldrich | A-003-E |
| Lipofectamine 3000 reagent | ThermoFisher Scientific | L3000015 |
| TransIT-LT1 transfection reagent | VWR | 731-0027 |
| jetOPTIMUS transfection reagent | Polyplus | 101000006 |
| Methanol-free formaldehyde | ThermoFisher Scientific | 28908 |
| DL-2-Amino-5-phosphonovaleric acid (APV) | Sigma-Aldrich | A5282 |
| NBQX disodium salt | Abcam | ab120046 |
| Forskolin (Fsk) | TCI | F0855 |
| H-89 dihydrochloride hydrate | Sigma-Aldrich | B1427 |

| | | |
|---|---|---|
| 3-Isobutyl-1-methylxanthine (IBMX) | Alfa Aesar | J64598 |
| Isoproterenol (Iso) | Sigma-Aldrich | I6504 |
| N6,2'-O-Dibutyryladenosine 3',5'-cyclic monophosphate sodium salt ($Bt_2cAMP$) | Sigma-Aldrich | D0627 |
| (−)-Epinephrine (Epi) | Sigma-Aldrich | E4250 |
| L-(−)-Norepinephrine (+)-bitartrate salt monohydrate | Sigma-Aldrich | A9512 |
| (±)-Propranolol hydrochloride | Sigma-Aldrich | 40543 |
| Prostaglandin E1 (PGE1) | Avanti Neutral Lipids | 900100P |
| Rolipram (Rol) | TCI | R0110 |
| 2-Deoxy-D-glucose (2-DG) | Sigma-Aldrich | D6134 |
| Ionomycin (Iono) | Abcam | Ab120116 |
| SBI-0206965 | Sigma-Aldrich | SML1540 |
| Thapsigargin | ThermoFisher Scientific | T7458 |
| Anisomycin | Sigma-Aldrich | A9789 |
| JNK Inhibitor VIII | Sigma-Aldrich | 420135 |
| Phorbol 12-myristate 13-acetate (PMA) | Sigma-Aldrich | P1585 |
| Gö 6983 | Sigma-Aldrich | G1918 |
| Adenosine 5'-triphosphate magnesium salt | Sigma-Aldrich | A9187 |
| Fluoromount-G | SouthernBiotech | 0100-01 |
| Pluronic F-127 | ThermoFisher Scientific | P3000MP |
| SKF-81297 hydrobromide | MCE | HY-12236 |
| 0.45 µm PES membrane | Millipore | HPWP04700 |
| Falcon 875cm² Rectangular Straight Neck Cell Culture Multi-Flask, 5-layer with Vented Cap | Corning | 353144 |
| Arcturus PicoPure Frozen RNA Isolation Kit | ThermoFisher Scientific | KIT0204 |
| RNase-Free DNase Set | Qiagen | 79254 |
| Kapa HiFi HS RM (6.25 mL) | Roche | 07958935001 |
| KAPA NGS Library Quantification Kit - Illumina for 480 Light Cycler | Roche | 07960298001 |
| DNA High Sensitivity Kit | Agilent | 5067-4626 |
| QIAamp DNA Blood Maxi Kit | Qiagen | 51192 |
| QIAquick Gel Extraction Kit | Qiagen | 12578 |
| Protease inhibitor cocktail | Sigma-Aldrich | P1860 |
| Phosphatase inhibitor cocktail | Sigma-Aldrich | P5726 |
| SYTOX blue dead cell stain | ThermoFisher Scientific | S34857 |
| Pierce BCA Protein Assay Kit | ThermoFisher Scientific | 23225 |
| PKA Colorimetric Activity Kit | ThermoFisher Scientific | EIAPKA |

**Supplementary Table 4 | Composition of common buffers used in the study.**

| Buffer name | Composition |
|---|---|
| Activity buffer | 50 mM HEPES, 50 mM NaCl, pH 7.3 |
| Kinase assay buffer | 50 mM Tris-HCl, 10 mM $MgCl_2$, 0.1 mM EDTA, 2 mM DTT, 0.5 mg $mL^{-1}$ BSA, pH 7.5 |
| IMAC lysis buffer | 50 mM $KH_2PO_4$, 150 mM NaCl, 5 mM imidazole, 1 mM PMSF,0.25 mg $mL^{-1}$ lysozyme, pH 8.0 |
| IMAC wash buffer | 50 mM $KH_2PO_4$, 300 mM NaCl, 10 mM imidazole, pH 7.5 |
| IMAC elution buffer | 50 mM $KH_2PO_4$, 300 mM NaCl, 500 mM imidazole, pH 7.5 |
| TNT extraction buffer | 20 mM Tris, pH 7.5, 150 mM NaCl, 1% (vol/vol) Triton X-100,10 mM $MgCl_2$ |
| Activated cell lysis buffer | Cell lysis buffer provided by PKA Colorimetric Activity Kit, with 0.1% (vol/vol) protease inhibitor cocktail, 1 mM PMSF, 1% (vol/vol) phosphatase inhibitor cocktail |
| Slicing buffer | 110 mM Choline-Cl, 2.5 mM KCl, 7 mM $MgCl_2$, 1 mM $NaH_2PO_4$, 0.5 mM $CaCl_2$, 25 mM $NaHCO_3$, 25 mM glucose, pH 7.4 |
| Artificial cerebrospinal fluid (ACSF) | 125 mM NaCl, 2.5 mM KCl, 1.3 mM $MgCl_2$, 1 mM $NaH_2PO_4$, 2 mM $CaCl_2$, 25 mM $NaHCO_3$, 25 mM glucose, pH 7.4 |

**Supplementary Table 5 | Plasmids and stable cell lines used in the study.**

| Construct | Purpose | Addgene # | Gene of interest | Stable cell line |
|---|---|---|---|---|
| PKAcat | Expressing PKA catalytic subunit alpha | #14921, gift from Susan Taylor | PKA catalytic subunit alpha | n.a. |
| pMD2.G | VSV-G envelope expressing plasmid | #12259, gift from Didier Trono | VSV G | n.a. |
| psPAX2 | 2nd generation lentiviral packaging plasmid | #12260, gift from Didier Trono | n.a. | n.a. |
| pET-51b(+) FHA1 | FHA1 protein production in *E .coli* | n.a. | FHA1 | n.a. |
| pET-51b(+) cpFHA1 | cpFHA1 protein production in *E .coli* | n.a. | cpFHA1 | n.a. |
| pET-51b(+) cpFHA1$^{N49Y}$ | cpFHA1$^{N49Y}$ protein production in *E .coli* | n.a. | cpFHA1$^{N49Y}$ | n.a. |
| pET-51b(+)-His10_TEVsite_NES-Kinprola$_{PKA}$-mEGFP | Kinprola$_{PKA}$ protein production in *E .coli* | n.a. | NES-Kinprola$_{PKA}$-mEGFP | n.a. |
| pET-51b(+)-His10_TEVsite_NES-Kinprola$_{PKA\ T/A}$-mEGFP | Kinprola$_{PKA\ T/A}$ protein production in *E .coli* | n.a. | NES-Kinprola$_{PKA\ T/A}$-mEGFP | n.a. |
| pET-51b(+)-His10_TEVsite_NES-Kinprola$_{on}$-mEGFP | Kinprola$_{PKA}$ protein production in *E .coli* | n.a. | NES-Kinprola$_{on}$-mEGFP | n.a. |
| pCDNA5/FRT_CMV_NES-Kinprola$_{PKA}$-mEGFP | Mammalian cell expression of Kinprola$_{PKA}$ in cytosol | TBD | NES-Kinprola$_{PKA}$-mEGFP | HEK293, HeLa |
| pCDNA5/FRT_CMV_NES-Kinprola$_{PKA_T/A}$-mEGFP | Mammalian cell expression of Kinprola$_{PKA_T/A}$ in cytosol | TBD | NES-Kinprola$_{PKA_T/A}$-mEGFP | HEK293, HeLa |
| pCDNA5/FRT_CMV_NES-Kinprola$_{on}$-mEGFP | Mammalian cell expression of Kinprola posCTRL in cytosol | TBD | NES-Kinprola$_{on}$-mEGFP | HEK293, HeLa |
| pCDNA5/FRT_CMV_NES-Kinprola$_{off}$-mEGFP | Mammalian cell expression of Kinprola negCTRL in cytosol | TBD | NES-Kinprola$_{off}$-mEGFP | HEK293, HeLa |
| pCDNA5/FRT_CMV_NES-Kinprola$_{PKC}$-mEGFP | Mammalian cell expression of Kinprola$_{PKC}$ in cytosol | TBD | NES-Kinprola$_{PKC}$-mEGFP | n.a. |
| pCDNA5/FRT_CMV_NES-Kinprola$_{PKC_T/A}$-mEGFP | Mammalian cell expression of Kinprola$_{PKC_T/A}$ in cytosol | TBD | NES-Kinprola$_{PKC_T/A}$-mEGFP | n.a. |
| pCDNA5/FRT_CMV_NES-Kinprola$_{JNK}$-mEGFP | Mammalian cell expression of Kinprola$_{JNK}$ in cytosol | TBD | NES-Kinprola$_{JNK}$-mEGFP | n.a. |
| pCDNA5/FRT_CMV_NES-Kinprola$_{JNK_T/A}$-mEGFP | Mammalian cell expression of Kinprola$_{JNK_T/A}$ in cytosol | TBD | NES-Kinprola$_{JNK_T/A}$-mEGFP | n.a. |
| pCDNA5/FRT_CMV_NES-Kinprola$_{AMPK}$-mEGFP | Mammalian cell expression of Kinprola$_{AMPK}$ in cytosol | TBD | NES-Kinprola$_{AMPK}$-mEGFP | n.a. |
| pCDNA5/FRT_CMV_NES-Kinprola$_{AMPK_T/A}$-mEGFP | Mammalian cell expression of Kinprola$_{AMPK_T/A}$ in cytosol | TBD | NES-Kinprola$_{AMPK_T/A}$-mEGFP | n.a. |
| pAAV_CAG_NES-Kinprola$_{PKA}$-mTagBFP2_WPRE-SV40 | Mammalian cell expression of Kinprola$_{PKA}$ in cytosol | TBD | NES-Kinprola$_{PKA}$-mTagBFP2 | n.a. |
| pAAV_CAG_Kinprola$_{PKA}$-mEGFP-NLS3×_WPRE-SV40 | Mammalian cell expression of Kinprola$_{PKA}$ in nucleus | TBD | Kinprola$_{PKA}$-mEGFP-NLS3× | n.a. |
| pAAV_hSyn_NES-Kinprola$_{PKA}$-mEGFP_WRPE-SV40 | Neuronal expression of Kinprola$_{PKA}$ in cytosol | TBD | NES-Kinprola$_{PKA}$-mEGFP | n.a. |
| pAAV_hSyn_NES-Kinprola$_{PKA_T/A}$-mEGFP_WRPE-SV40 | Neuronal expression of Kinprola$_{PKA_T/A}$ in cytosol | TBD | NES-Kinprola$_{PKA_T/A}$-mEGFP | n.a. |
| pAAV_hSyn_NES-Kinprola$_{on}$-mEGFP_WRPE-SV40 | Neuronal expression of Kinprola$_{on}$ in cytosol | TBD | NES-Kinprola$_{on}$-mEGFP | n.a. |
| pLKO.1-puro_CMV_NES-Kinprola$_{PKA}$-mEGFP | Glioblastoma expression of Kinprola$_{PKA}$ in cytosol | TBD | NES-Kinprola$_{PKA}$-mEGFP | GBC S24 |
| pLKO.1-puro_CMV_NES-Kinprola$_{on}$-mEGFP | Glioblastoma expression of Kinprola$_{on}$ in cytosol | TBD | NES-Kinprola$_{on}$-mEGFP | GBC S24 |
| pLenti_EF1α-NES-Kinprola$_{PKA}$-mEGFP_bGH-PA term_EF1α_FLAG-SV40NLS-Cas9-NLS-T2A-BSD-WPRE-SV40 | RKO expression of Kinprola$_{PKA}$ in cytosol and Cas9 in nucleus | TBD | NES-Kinprola$_{PKA}$-mEGFP, FLAG-SV40NLS-Cas9-NLS-T2A-BSD | RKO |
| pLenti_EF1α-NES-Kinprola$_{PKA_T/A}$-mEGFP_bGH-PA term_EF1α_FLAG-SV40NLS-Cas9-NLS-T2A-BSD-WPRE-SV40 | RKO expression of Kinprola$_{PKA_T/A}$ in cytosol and Cas9 in nucleus | TBD | NES-Kinprola$_{PKA_T/A}$-mEGFP, FLAG-SV40NLS-Cas9-NLS-T2A-BSD | RKO |
| pLenti_EF1α-NES-Kinprola$_{on}$-mEGFP_bGH-PA term_EF1α_FLAG-SV40NLS-Cas9-NLS-T2A-BSD-WPRE-SV40 | RKO expression of Kinprola$_{on}$ in cytosol and Cas9 in nucleus | TBD | NES-Kinprola$_{on}$-mEGFP, FLAG-SV40NLS-Cas9-NLS-T2A-BSD | RKO |
| HD CRISPR library sub-library A | CRISPR screen | n.a. | genome-scale sgRNA library | n.a. |
| HDCRISPRv1_U6_sg*FZR1*-1 | sgRNA expression for *FZR1* knockout | n.a. | sg*FZR1*-1 | n.a. |
| HDCRISPRv1_U6_sg*FZR1*-2 | sgRNA expression for *FZR1* knockout | n.a. | sg*FZR1*-2 | n.a. |
| HDCRISPRv1_U6_sg*TRIM33*-1 | sgRNA expression for *TRIM33* knockout | n.a. | sg*TRIM33*-1 | n.a. |
| HDCRISPRv1_U6_sg*TRIM33*-2 | sgRNA expression for *TRIM33* knockout | n.a. | sg*TRIM33*-2 | n.a. |
| HDCRISPRv1_U6_sg*PES1*-1 | sgRNA expression for *PES1* knockout | n.a. | sg*PES1*-1 | n.a. |
| HDCRISPRv1_U6_sg*PES1*-2 | sgRNA expression for *PES1* knockout | n.a. | sg*PES1*-2 | n.a. |
| HDCRISPRv1_U6_sg*NOP2*-1 | sgRNA expression for *NOP2* knockout | n.a. | sg*NOP2*-1 | n.a. |
| HDCRISPRv1_U6_sg*NOP2*-2 | sgRNA expression for *NOP2* knockout | n.a. | sg*NOP2*-2 | n.a. |
| HDCRISPRv1_U6_sg*VPRBP*-1 | sgRNA expression for *VPRBP* knockout | n.a. | sg*VPRBP*-1 | n.a. |

| | | | | |
|---|---|---|---|---|
| HDCRISPRv1_U6_sg*VPRBP*-2 | sgRNA expression for *VPRBP* knockout | n.a. | sg*VPRBP*-2 | n.a. |
| HDCRISPRv1_U6_sg*Non-targeting*-1 | sgRNA expression for non-targeting sequence | n.a. | sg*Non-targeting*-1 | n.a. |
| HDCRISPRv1_U6_sg*Non-targeting*-2 | sgRNA expression for non-targeting sequence | n.a. | sg*Non-targeting*-2 | n.a. |

**Supplementary Table 6 | sgRNA sequences selected from HD CRISPR sub-library A.**

| Gene ID | Gene name | sgRNA sequence 1 | sgRNA sequence 2 |
|---|---|---|---|
| ENSG00000105325 | *FZR1* | CCGCTCAGACCAGCCCACGG | CAGCAGCTCATTCTTGAGCA |
| ENSG00000197323 | *TRIM33* | CTTGCAGAGCCGGCGTGAGG | TTACTAAAGATCACTTGATC |
| ENSG00000100029 | *PES1* | GCATGAACTCCACAGTGAGC | GCTGTGCCGCCGGCTCACTG |
| ENSG00000111641 | *NOP2* | GTATTGGTCCGGAGGGTGAC | GAAGATGGTATGGTGAACCA |
| ENSG00000145041 | *VPRBP* | ACATGGTACCTATCCTTACC | TATTCATACCTGGTAAGGAT |
| n.a. | *Non-targeting* | GTGACTAGACCCTTACGCGG | GATCGGCGGGTTACCTCTGA |

**Supplementary Table 7 | Imaging acquisition parameters for fluorescence microscopy.**

| Figure | Construct | Label | Microscope | Objective | Excitation (nm) | Emission (nm) | Fluorophore concentration |
|---|---|---|---|---|---|---|---|
| **1c** | NES-Kinprola$_{PKA}$-mEGFP | mEGFP | Leica Stellaris 5 confocal | 20× water | 488 | 498-550 | - |
| | | CPY | | 0.75-NA | 610 | 620-720 | 25 nM |
| **1e** | NES-Kinprola$_{PKA}$-mEGFP | mEGFP | Leica Stellaris 5 confocal | 20× water | 488 | 498-550 | - |
| | | CPY | | 0.75-NA | 610 | 620-720 | 25 nM |
| **1h** | NES-Kinprola$_{PKA}$-mEGFP | mEGFP | Leica Stellaris 5 confocal | 20× water | 488 | 498-550 | - |
| | | JF$_{552}$ | | 0.75-NA | 550 | 560-600 | 25 nM |
| | | CPY | | | 610 | 620-660 | 100 nM |
| | | JF$_{669}$ | | | 669 | 679-750 | 100 nM |
| **4a** | NES-Kinprola$_{PKA}$-mEGFP | mEGFP | Leica Stellaris 5 confocal | 20× water | 488 | 498-550 | - |
| | NES-Kinprola$_{PKA_T/A}$-mEGFP | CPY | | 0.75-NA | 610 | 620-720 | 25 nM |
| | NES-Kinprola$_{on}$-mEGFP | | | | | | |
| **4c** | NES-Kinprola$_{PKA}$-mEGFP | mEGFP | Leica Stellaris 5 confocal | 20× water | 488 | 498-550 | - |
| | | CPY | | 0.75-NA | 610 | 620-720 | 125 nM |
| **5b,c** | NES-Kinprola$_{PKA}$-mEGFP | mEGFP | ZEISS LSM 710 upright confocal | 20× water | 488 | 498-550 | - |
| | NES-Kinprola$_{PKA_T/A}$-mEGFP | CPY | | 0.5-NA | 633 | 638-747 | 250 nM |
| **5d** | NES-Kinprola$_{PKA}$-mEGFP | mEGFP | Leica SP8X | 40× oil | 488 | 498-550 | - |
| | NES-Kinprola$_{PKA_T/A}$-mEGFP | CPY | | 1.3-NA | 610 | 620-720 | 250 nM |
| **5h** | NES-Kinprola$_{PKA}$-mEGFP | mEGFP | Leica SP8X | 20× dry | 488 | 498-550 | - |
| | | CPY | | 0.7-NA | 610 | 620-720 | 100 nmol |
| **Extended Data Fig. 5** | NES-Kinprola$_{PKA}$-mEGFP | mEGFP | Leica Stellaris 5 confocal | 20× water | 488 | 498-550 | - |
| | | JF$_{552}$ | | 0.75-NA | 550 | 560-600 | 25 nM |
| | | CPY | | | 610 | 620-660 | 100 nM |
| | | JF$_{669}$ | | | 669 | 679-750 | 100 nM |
| **Extended Data Fig. 6d** | NES-Kinprola$_{PKA}$-mTagBFP2 | mTagBFP2 | Leica Stellaris 5 confocal | 40× water | 405 | 415-480 | - |
| | Kinprola$_{PKA}$-mEGFP-NLS3× | mEGFP | | 1.1-NA | 488 | 498-550 | - |
| | | CPY | | | 612 | 620-700 | 50 nM |
| **Extended Data Fig. 9a,e** | NES-Kinprola$_{PKA}$-mEGFP | mEGFP | Leica Stellaris 5 confocal | 20× water | 488 | 498-550 | - |
| | | CPY | | 0.75-NA | 610 | 620-720 | 125 nM |
| **Extended Data Fig. 9c** | NES-Kinprola$_{PKA}$-mEGFP | mEGFP | Leica Stellaris 5 confocal | 20× water | 488 | 498-550 | - |
| | | CPY | | 0.75-NA | 610 | 620-720 | 25 nM |
| **Extended Data Fig. 10a** | NES-Kinprola$_{PKA}$-mEGFP | mEGFP | Leica SP8X | 20× dry | 488 | 498-550 | - |

| | | | | | | | |
|---|---|---|---|---|---|---|---|
| | | CPY | | 0.7-NA | 610 | 620-720 | 100 nmol |
| **Supplementary Fig. 2** | NES-Kinprola$_{PKA}$-mEGFP | mEGFP | Leica Stellaris 5 confocal | 20× water | 488 | 498-550 | - |
| | | CPY | | 0.75-NA | 610 | 620-720 | 25 nM |
| **Supplementary Fig. 6b** | NES-Kinprola$_{PKA}$-mEGFP | mEGFP | Leica Stellaris 5 confocal | 20× water | 488 | 498-550 | - |
| | | CPY | | 0.75-NA | 610 | 620-720 | 125 nM |

**Supplementary Note 1 | Protein sequences**

>Kinprola$_{PKA}$
Elements: NES-cpHaloΔ-PKA substrate-cpFHA1-Hpep-mEGFP
MLQNELALKLAGLDINKTGGSDVGRKLIIDQNVFIEGTLPMGVVRPLTEVEMDHYREPFLNPVDREPL
WRFPNELPIAGEPANIVALVEEYMDWLHQSPVPKLLFWGTPGVLIPPAEAARLAKSLPNCKAVDIGPG
LNLLQEDNPDLIGSEIARWLSTLEIGGTGGSGGTGGSGGSIGTGFPFDPHYVEVLGERMHYVDVGPRD
GTPVLFLHGNPTSSYVWRNIIPHVAPTHRCIAPDLIGMGKSDKPDLGYFFDDHVRFMDAFIEALGLEE
VVLVIHDWGSALGFHWAKRNPERVKGIAFMEFIRPIPTWDEWAPGSLRRATLVDPPPPPPPPPPPPPP
PPPPPPPPPPPPPPPPGGSACDYHLGNISRLSNKHFQILLGEDGNLLLNDISTNGTWLNGQKVEKNSY
QLLSQGDEITVGVGVESDILSLVIFINDKFKQCLEQNKVDRGGGMHKFSQEQIGENIVCRVICTTGQI
PIRDLSADISQVLKEKRSIKKVWTFGRNPGSAKETFQAFRSGGSMVSKGEELFTGVVPILVELDGDVN
GHKFSVSGEGEGDATYGKLTLKFICTTGKLPVPWPTLVTTLTYGVQCFSRYPDHMKQHDFFKSAMPEG
YVQERTIFFKDDGNYKTRAEVKFEGDTLVNRIELKGIDFKEDGNILGHKLEYNYNSHNVYIMADKQKN
GIKVNFKIRHNIEDGSVQLADHYQQNTPIGDGPVLLPDNHYLSTQSKLSKDPNEKRDHMVLLEFVTAA
GITLGMDELYK

>Kinprola$_{PKA_T/A}$
Elements: NES-cpHaloΔ-PKA substrate T/A-cpFHA1-Hpep-mEGFP
MLQNELALKLAGLDINKTGGSDVGRKLIIDQNVFIEGTLPMGVVRPLTEVEMDHYREPFLNPVDREPL
WRFPNELPIAGEPANIVALVEEYMDWLHQSPVPKLLFWGTPGVLIPPAEAARLAKSLPNCKAVDIGPG
LNLLQEDNPDLIGSEIARWLSTLEIGGTGGSGGTGGSGGSIGTGFPFDPHYVEVLGERMHYVDVGPRD
GTPVLFLHGNPTSSYVWRNIIPHVAPTHRCIAPDLIGMGKSDKPDLGYFFDDHVRFMDAFIEALGLEE
VVLVIHDWGSALGFHWAKRNPERVKGIAFMEFIRPIPTWDEWAPGSLRRAALVDPPPPPPPPPPPPPP
PPPPPPPPPPPPPPPPGGSACDYHLGNISRLSNKHFQILLGEDGNLLLNDISTNGTWLNGQKVEKNSY
QLLSQGDEITVGVGVESDILSLVIFINDKFKQCLEQNKVDRGGGMHKFSQEQIGENIVCRVICTTGQI
PIRDLSADISQVLKEKRSIKKVWTFGRNPGSAKETFQAFRSGGSMVSKGEELFTGVVPILVELDGDVN
GHKFSVSGEGEGDATYGKLTLKFICTTGKLPVPWPTLVTTLTYGVQCFSRYPDHMKQHDFFKSAMPEG
YVQERTIFFKDDGNYKTRAEVKFEGDTLVNRIELKGIDFKEDGNILGHKLEYNYNSHNVYIMADKQKN
GIKVNFKIRHNIEDGSVQLADHYQQNTPIGDGPVLLPDNHYLSTQSKLSKDPNEKRDHMVLLEFVTAA
GITLGMDELYK

>Kinprola$_{on}$
Elements: NES-cpHalo-PKA substrate-cpFHA1-Hpep-mEGFP
MLQNELALKLAGLDINKTGGSFARETFQAFRTTDVGRKLIIDQNVFIEGTLPMGVVRPLTEVEMDHYR
EPFLNPVDREPLWRFPNELPIAGEPANIVALVEEYMDWLHQSPVPKLLFWGTPGVLIPPAEAARLAKS
LPNCKAVDIGPGLNLLQEDNPDLIGSEIARWLSTLEIGGTGGSGGTGGSGGSIGTGFPFDPHYVEVLG
ERMHYVDVGPRDGTPVLFLHGNPTSSYVWRNIIPHVAPTHRCIAPDLIGMGKSDKPDLGYFFDDHVRF
MDAFIEALGLEEVVLVIHDWGSALGFHWAKRNPERVKGIAFMEFIRPIPTWDEWAPGSLRRATLVDPP
PPPPPPPPPPPPPPPPPPPPPPPPPPPPGGSACDYHLGNISRLSNKHFQILLGEDGNLLLNDISTNGT
WLNGQKVEKNSYQLLSQGDEITVGVGVESDILSLVIFINDKFKQCLEQNKVDRGGGMHKFSQEQIGEN
IVCRVICTTGQIPIRDLSADISQVLKEKRSIKKVWTFGRNPGSAKETFQAFRSGGSMVSKGEELFTGV
VPILVELDGDVNGHKFSVSGEGEGDATYGKLTLKFICTTGKLPVPWPTLVTTLTYGVQCFSRYPDHMK
QHDFFKSAMPEGYVQERTIFFKDDGNYKTRAEVKFEGDTLVNRIELKGIDFKEDGNILGHKLEYNYNS
HNVYIMADKQKNGIKVNFKIRHNIEDGSVQLADHYQQNTPIGDGPVLLPDNHYLSTQSKLSKDPNEKR
DHMVLLEFVTAAGITLGMDELYK

>Kinprola$_{off}$
Elements: NES-cpHaloΔ-PKA substrate-cpFHA1-mEGFP
MLQNELALKLAGLDINKTGGSFARETFQAFRTTDVGRKLIIDQNVFIEGTLPMGVVRPLTEVEMDHYR
EPFLNPVDREPLWRFPNELPIAGEPANIVALVEEYMDWLHQSPVPKLLFWGTPGVLIPPAEAARLAKS
LPNCKAVDIGPGLNLLQEDNPDLIGSEIARWLSTLEIGGTGGSGGTGGSGGSIGTGFPFDPHYVEVLG
ERMHYVDVGPRDGTPVLFLHGNPTSSYVWRNIIPHVAPTHRCIAPDLIGMGKSDKPDLGYFFDDHVRF
MDAFIEALGLEEVVLVIHDWGSALGFHWAKRNPERVKGIAFMEFIRPIPTWDEWAPGSLRRATLVDPP
PPPPPPPPPPPPPPPPPPPPPPPPPPPPGGSACDYHLGNISRLSNKHFQILLGEDGNLLLNDISTNGT

WLNGQKVEKNSYQLLSQGDEITVGVGVESDILSLVIFINDKFKQCLEQNKVDRGGGMHKFSQEQIGEN
IVCRVICTTGQIPIRDLSADISQVLKEKRSIKKVWTFGRNPGGSMVSKGEELFTGVVPILVELDGDVN
GHKFSVSGEGEGDATYGKLTLKFICTTGKLPVPWPTLVTTLTYGVQCFSRYPDHMKQHDFFKSAMPEG
YVQERTIFFKDDGNYKTRAEVKFEGDTLVNRIELKGIDFKEDGNILGHKLEYNYNSHNVYIMADKQKN
GIKVNFKIRHNIEDGSVQLADHYQQNTPIGDGPVLLPDNHYLSTQSKLSKDPNEKRDHMVLLEFVTAA
GITLGMDELYK

>mTagBFP2 (C-terminally fused)
MVSKGEELIKENMHMKLYMEGTVDNHHFKCTSEGEGKPYEGTQTMRIKVVEGGPLPFAFDILATSFLY
GSKTFINHTQGIPDFFKQSFPEGFTWERVTTYEDGGVLTATQDTSLQDGCLIYNVKIRGVNFTSNGPV
MQKKTLGWEAFTETLYPADGGLEGRNDMALKLVGGSHLIANAKTTYRSKKPAKNLKMPGVYYVDYRLE
RIKEANNETYVEQHEVAVARYCDLPSKLGHKLN

>Cas9 expression
Elements: FLAG-SV40 NLS-Cas9-nucleoplasmin NLS-T2A-BSD
MDYKDDDDKMAPKKKRKVGIHGVPAADKKYSIGLDIGTNSVGWAVITDEYKVPSKKFKVLGNTDRHSI
KKNLIGALLFDSGETAEATRLKRTARRRYTRRKNRICYLQEIFSNEMAKVDDSFFHRLEESFLVEEDK
KHERHPIFGNIVDEVAYHEKYPTIYHLRKKLVDSTDKADLRLIYLALAHMIKFRGHFLIEGDLNPDNS
DVDKLFIQLVQTYNQLFEENPINASGVDAKAILSARLSKSRRLENLIAQLPGEKKNGLFGNLIALSLG
LTPNFKSNFDLAEDAKLQLSKDTYDDDLDNLLAQIGDQYADLFLAAKNLSDAILLSDILRVNTEITKA
PLSASMIKRYDEHHQDLTLLKALVRQQLPEKYKEIFFDQSKNGYAGYIDGGASQEEFYKFIKPILEKM
DGTEELLVKLNREDLLRKQRTFDNGSIPHQIHLGELHAILRRQEDFYPFLKDNREKIEKILTFRIPYY
VGPLARGNSRFAWMTRKSEETITPWNFEEVVDKGASAQSFIERMTNFDKNLPNEKVLPKHSLLYEYFT
VYNELTKVKYVTEGMRKPAFLSGEQKKAIVDLLFKTNRKVTVKQLKEDYFKKIECFDSVEISGVEDRF
NASLGTYHDLLKIIKDKDFLDNEENEDILEDIVLTLTLFEDREMIEERLKTYAHLFDDKVMKQLKRRR
YTGWGRLSRKLINGIRDKQSGKTILDFLKSDGFANRNFMQLIHDDSLTFKEDIQKAQVSGQGDSLHEH
IANLAGSPAIKKGILQTVKVVDELVKVMGRHKPENIVIEMARENQTTQKGQKNSRERMKRIEEGIKEL
GSQILKEHPVENTQLQNEKLYLYYLQNGRDMYVDQELDINRLSDYDVDHIVPQSFLKDDSIDNKVLTR
SDKNRGKSDNVPSEEVVKKMKNYWRQLLNAKLITQRKFDNLTKAERGGLSELDKAGFIKRQLVETRQI
TKHVAQILDSRMNTKYDENDKLIREVKVITLKSKLVSDFRKDFQFYKVREINNYHHAHDAYLNAVVGT
ALIKKYPKLESEFVYGDYKVYDVRKMIAKSEQEIGKATAKYFFYSNIMNFFKTEITLANGEIRKRPLI
ETNGETGEIVWDKGRDFATVRKVLSMPQVNIVKKTEVQTGGFSKESILPKRNSDKLIARKKDWDPKKY
GGFDSPTVAYSVLVVAKVEKGKSKKLKSVKELLGITIMERSSFEKNPIDFLEAKGYKEVKKDLIIKLP
KYSLFELENGRKRMLASAGELQKGNELALPSKYVNFLYLASHYEKLKGSPEDNEQKQLFVEQHKHYLD
EIIEQISEFSKRVILADANLDKVLSAYNKHRDKPIREQAENIIHLFTLTNLGAPAAFKYFDTTIDRKR
YTSTKEVLDATLIHQSITGLYETRIDLSQLGGDKRPAATKKAGQAKKKKASGSGEGRGSLLTCGDVEE
NPGPMAKPLSQEESTLIERATATINSIPISEDYSVASAALSSDGRIFTGVNVYHFTGGPCAELVVLGT
AAAAAAGNLTCIVAIGNENRGILSPCGRCRQVLLDLHPGIKAIVKDSDGQPTAVGIRELLPSGYVWEG

| Different kinases | Substrate | T/A mutant |
|---|---|---|
| PKC | RFRRFQTLKDKAKA | RFRRFQALKDKAKA |
| JNK | DSVKTPEDEGNPLLEQLEKK | DSVKAPEDEGNPLLEQLEKK |
| AMPK | MRRVATLVDL | MRRVAALVDL |

**Localization sequences**

>Nuclear export sequence (NES, N-terminally)
(M) LQNELALKLAGLDINKT
>Nuclear localization sequence (NLS, C-terminally, 3 copies)
KSGLRSRADPKKKRKVDPKKKRKVDPKKKRKVGSTGSR

**Purification sequences**

>Poly-histidine tag + TEV cleavage sequence (N-terminal)
HHHHHHHHHHENLYFQGG (pET-51b(+) plasmid)
